# A fast genetically encoded fluorescent sensor for faithful *in vivo* acetylcholine detection in mice, fish, worms and flies

**DOI:** 10.1101/2020.02.07.939504

**Authors:** Philip M. Borden, Peng Zhang, Amol V. Shivange, Jonathan S. Marvin, Joseph Cichon, Chuntao Dan, Kaspar Podgorski, Antonio Figueiredo, Ondrej Novak, Masashi Tanimoto, Eiji Shigetomi, Mark A. Lobas, Hyuntae Kim, Paula K. Zhu, Yajun Zhang, W. Sharon Zheng, ChengCheng Fan, Guangfu Wang, Bowen Xiang, Li Gan, Guang-Xian Zhang, Kaiming Guo, Li Lin, Yuan Cai, Andrew G. Yee, Abhi Aggarwal, Christopher P. Ford, Douglas C. Rees, Dirk Dietrich, Baljit S. Khakh, Jeremy S. Dittman, Wen-Biao Gan, Minoru Koyama, Vivek Jayaraman, Joseph F. Cheer, Henry A. Lester, J. Julius Zhu, Loren L. Looger

## Abstract

Here we design and optimize a genetically encoded fluorescent indicator, iAChSnFR, for the ubiquitous neurotransmitter acetylcholine, based on a bacterial periplasmic binding protein. iAChSnFR shows large fluorescence changes, rapid rise and decay kinetics, and insensitivity to most cholinergic drugs. iAChSnFR revealed large transients in a variety of slice and *in vivo* preparations in mouse, fish, fly and worm. iAChSnFR will be useful for the study of acetylcholine in all animals.

## Introduction

Acetylcholine (ACh) is a critical neurotransmitter in all animals. Among invertebrates, it is the most prevalent excitatory transmitter in the brain, sensory ganglia, and frequently the neuromuscular junction (NMJ). Among vertebrates, only a minority of neurons release ACh, but these signals play varying key roles. For instance, ACh signals at the NMJ, in the autonomic nervous system, and in subsets of the central nervous system, particularly projections arising from the brainstem and basal forebrain. Other cholinergic neuron populations in the brain include striatal interneurons, the stria vascularis-medial habenula-interpeduncular nucleus pathway, and sparse, incompletely characterized cell types such as intrinsic cholinergic interneurons in cortex^1^ and hippocampus^2^. ACh helps to regulate attention^3^ and wakefulness^4^, and participates in memory formation and consolidation^5^. ACh is also an important transmitter in glia, and between the nervous and immune systems^6^.

Acetylcholine is synthesized pre-synaptically from choline and acetyl-CoA by choline acetyltransferase (ChAT), then packaged into synaptic vesicles by the vesicular acetylcholine transporter (VAChT). A key, partially understood aspect of cholinergic signaling is co-release with other neurotransmitters, including GABA, ATP, and glutamate^7, 8^. To understand the role of co-release, one must measure ACh release alongside emerging measurements of other neurotransmitters.

Acetylcholine receptors are among the most diverse neurotransmitter receptor families. Humans possess five muscarinic G protein-coupled receptors (GPCRs) for ACh (mAChRs) with diverse expression in the brain and smooth, cardiac, and skeletal muscle. Vertebrate nicotinic ACh receptors (nAChRs) are pentameric ligand-gated cation channels. Humans have a total of 17 nAChR subunit genes, in five classes: 10 α, 4 β, and one each of γ, δ, and ε. nAChRs occur with many subunit combinations^9^, and others may be undiscovered. Invertebrates also have ACh-gated chloride channels. On neurons, receptors can be localized pre-, post-, and extra-synaptically, often with different isoforms in each place^10^.

ACh released at or near synapses is inactivated primarily *via* hydrolysis to choline by acetylcholinesterase (AChE)^11^. Numerous molecules, both naturally occurring and synthetic, bind to and perturb VAChT, AChRs, or AChE. Cholinergic drugs are used to treat neurodegenerative conditions such as Parkinson’s disease and Alzheimer’s disease, neuromuscular diseases like myasthenia gravis, autoimmune disorders such as asthma and rheumatoid arthritis, substance abuse disorders including nicotine addiction, and autonomic diseases such as incontinence and nausea^12^. Most cholinergic drugs have side effects stemming from the broad expression, function, and diversity of ACh signaling. Several insecticides and chemical warfare agents block AChE. A better understanding of ACh signaling is thus necessary for the design and development of improved pharmaceuticals and agricultural products.

Although ACh was discovered at the beginning of the 20^th^ century^13^, methods for ACh detection are scant. ACh receptors themselves give an imperfect spatial and temporal report on the presence of ACh, because (a) ACh receptors are located at various distances between 0.1 μm and ∼ 50 μm from the release point (at the neuromuscular junction and in cardiac muscle, respectively), and (b) because ACh receptors show a delay ranging from a few μs to several tens of ms (and nAChRs and mAChRs, respectively) after ACh is applied nearby^14^. Indeed, despite the widespread presence of many nAChR subtypes in the vertebrate CNS, direct recordings of postsynaptic responses are quite rare at nicotinic synapses stimulated pre-synaptically; and direct recording of ACh release is needed to understand cholinergic signaling. For half a century, the most sensitive assay for acetylcholine was observation of its stimulatory effect on leech muscle treated with eserine (an AChE inhibitor). Advances in chemical derivatization and chromatography permitted its direct detection in the 1960s^15^, but this technique has very poor spatial and temporal resolution. Other methods detect ACh electrochemically – an inherently slower, indirect technique without molecular specificity^16^. Improvements in ACh detection have come from the development of semisynthetic, FRET-based probes^17, 18^ and whole cell-based fluorescence reporters^19^, but these are limited by low signal change, poor spatial and temporal resolution, and difficulty in delivery.

Protein-based biosensors allow targeted expression in cells, and even sub-cellular compartments, of interest. A recently published sensor (GACh2.0)^20^ couples ACh binding to a change in the fluorescence of circularly permuted green fluorescent protein (cpGFP) inserted within an intracellular loop of the M_3_ mAChR. GACh2.0 has an apparent ACh affinity of 700 nM, a maximal fluorescence increase of ∼80% in cultured cells, and kinetics of ∼200 ms τ_rise_ and ∼700 ms τ_decay_^20^. GACh2.0 was validated in neuronal culture, organotypic hippocampal slice, and acute slice, with resolvable differences in acetylcholine release being detected on the micrometer scale of a single soma. *In vivo*, GACh2.0 can detect odor-induced acetylcholine release with 2-photon microscopy in sub-regions of the *Drosophila melanogaster* antennal lobe. It also detected acetylcholine release in mouse visual cortex. However, molecular characteristics of the GACh2.0 sensor restrict its utility. Foremost, in flies and mice GACh2.0 requires seconds for the onset and decay of ACh responses^20^. This precludes resolution of faster signaling events, hinders detection of sparse events, and thus renders GACh2.0 essentially a binary detector of ACh within a region of interest over several seconds. Moreover, the sensor responds (positively or negatively) to all molecules that normally bind the M_3_ receptor, precluding the use of GACh2.0 in combination with many cholinergic drugs. Finally, over-expression of GPCRs perturbs the function of endogenous proteins such as G-proteins^21, 22^ and can disrupt cellular behavior^23^, creating concerns that GACh2.0-expressing cells may have altered ACh and/or other signaling pathways.

In a complementary strategy, we have coupled ligand binding to fluorescence with nearly modular fusion proteins. These utilize microbial periplasmic binding proteins (PBPs) as the molecular recognition moiety and circularly permuted GFP or its analogs as the reporter moieties. The strategy has created sensors for glutamate (iGluSnFR^24^, SF-iGluSnFR^25^), GABA (iGABASnFR)^26^, glucose (iGlucoSnFR^27^, iGlucoSnFR-TS^28^), maltose^29^, and phosphonate^30^. Because of their basis on microbial proteins, they are bio-orthogonal to endogenous signaling pathways in model organisms, permitting both long-term *in vivo* expression and use alongside most relevant pharmaceuticals. Sensors derived from PBPs generally have rise and decay kinetics on a millisecond time scale approaching that of postsynaptic ligand-gated ion channels, thus permitting useful imaging at >1 kHz^24, 30, 31^, appropriate to millisecond-scale neurotransmission.

Here we extend the collection of PBP-based sensors to include an intensity-based <u>a</u>cetyl<u>ch</u>oline <u>s</u>e<u>n</u>sing <u>f</u>luorescent <u>r</u>eporter, iAChSnFR. iAChSnFR detects ACh with low-micromolar affinity and 12-fold maximal fluorescence change *in vitro* (4.5-fold change in neuronal culture), has millisecond-level activation and decay kinetics, is orthogonal to most cholinergic drugs, and is available as both green and yellow fluorescent versions. iAChSnFR detects ACh release *in vivo* in the brains of mice, fish, and flies, and at the worm NMJ. In mouse cortex *in vivo*, iAChSnFR tracked running-dependent cholinergic modulation from basal forebrain, localizing ACh transients with spatial and temporal resolutions of tens of milliseconds and <1 μm). Imaging in visual cortex confirmed these results, and further demonstrated depth-dependent modulation by isoflurane anesthesia. Acetylcholinesterase inhibitors increased signals, and a non-binding sensor showed no response, together demonstrating that iAChSnFR specifically measures ACh. iAChSnFR will be useful for numerous experiments across model organisms.

## Results

### Sensor engineering

We previously developed a generalizable technique for developing intensity-based fluorescence sensors from solute-binding proteins that undergo a Venus flytrap-like conformational change^29^. Several such proteins that bind choline and/or betaine have been characterized biochemically and structurally. We first developed a sensor based on the acetylcholine/choline-binding protein ChoX from the soil bacterium *Sinorhizobium meliloti*^31^, but choline-dependent fluorescence increases were unacceptably slow (minutes; not shown). We developed another sensor from the *Bacillus subtilis* choline-binding protein OpuBC^32^, but when expressed in mammalian cells with membrane-targeting motifs, it aggregated in the endoplasmic reticulum (not shown).

We identified a hyperthermophilic homologue of *B. subtilis* OpuBC, from *Thermoanaerobacter sp. X513* (38% overall identity; 4/9 key ligand-binding residues conserved; 5/9 similar; **Supp. Fig. S1**). Isothermal titration calorimetry of purified X513-OpuBC confirmed binding of choline (*K*_d_ = 8 µM) and acetylcholine (*K*_d_ = 95 µM) (**Supp. Fig. S2**). This thermophilic scaffold provided the basis for iAChSnFR.

Insertion of circularly permuted Superfolder GFP (SFGFP)^33^ after residue Gly106, followed by optimization of the binding protein-GFP linkers, yielded an initial variant (iAChSnFR0.4; **Supp. Fig. S3**) with an apparent *K*_d_ of 220 ± 16 µM for ACh and a total fluorescence increase (ΔF/F)_max_ of ∼0.7, but with tighter affinity for choline and betaine (not shown). A crystal structure of the unliganded/open form of iAChSnFR0.4 showed the expected structural homology with *B. subtilis* OpuBC^34^ and with several other choline-binding PBPs (PDB 1SW1, 3R6U, 2REG, 2RIN).

Guided by the ligand-free crystal structure of this initial variant (**Fig. 1a**), we extensively screened for variants with increased specificity and affinity for ACh and increased ΔF/F upon saturation with ACh. Intermediate variants included iAChSnFR0.6 and iAChSnFR0.7; **Supp. Fig. S3**). The final variant, iAChSnFR0.9 (hereon referred to as iAChSnFR) has five binding-site mutations to the starting scaffold. iAChSnFR also has three mutations on the periphery of the binding site, two mutations in the “hinge” to shift the open-closed equilibrium^35^, four mutations to the proto-interface between X513 and cpSFGFP, and seven mutations to the junctions between the binding protein and cpSFGFP (**Supp. Fig. S3**). We solved the crystal structure of the ACh-bound/closed state of X513 OpuBC, without the GFP (**Fig. 1b**), revealing the importance of mutations such as Phe219Trp and Glu174Phe (aromatic cage around choline), Arg178Gly (creates space for acetyl moiety), and Lys39Ile (creates space for and hydrophobic contact with acetyl). iAChSnFR binds acetylcholine with an apparent *K*_d_ of 1.3 ± 1.0 µM, (ΔF/F)_max_ of ∼ 12, much weaker sensitivity to choline (apparent *K*_d_ 45 ± 1.1 µM) (**Fig. 1c**), and no detectible affinity for betaine nor a panel of neurotransmitters (except serotonin, which it binds with >1 mM affinity; **Supp. Fig. S4**). iAChSnFR also detects the AChE inhibitors neostigmine, physostigmine, and rivastigmine, but not structurally divergent inhibitors such as tacrine or donepezil (**Supp. Fig. S5a**). It is actually a rather sensitive sensor for nicotine and oxotremorine (**Supp. Fig. S5b**) and differs by only a few amino acids from the simultaneously developed nicotine sensor, iNicSnFR^36^. For control experiments, we made a non-binding iAChSnFR mutant (iAChSnFR-NULL; mutation Tyr357Ala; no response > 100 µM ACh, choline *K*_d_ > 50 mM; **Supp. Fig. S6**). Consistent with its construction from a hyperthermophilic binding protein, iAChSnFR is extremely stable, unfolding at ∼90°C (**Supp. Fig. S7**).

**Fig. 1.**
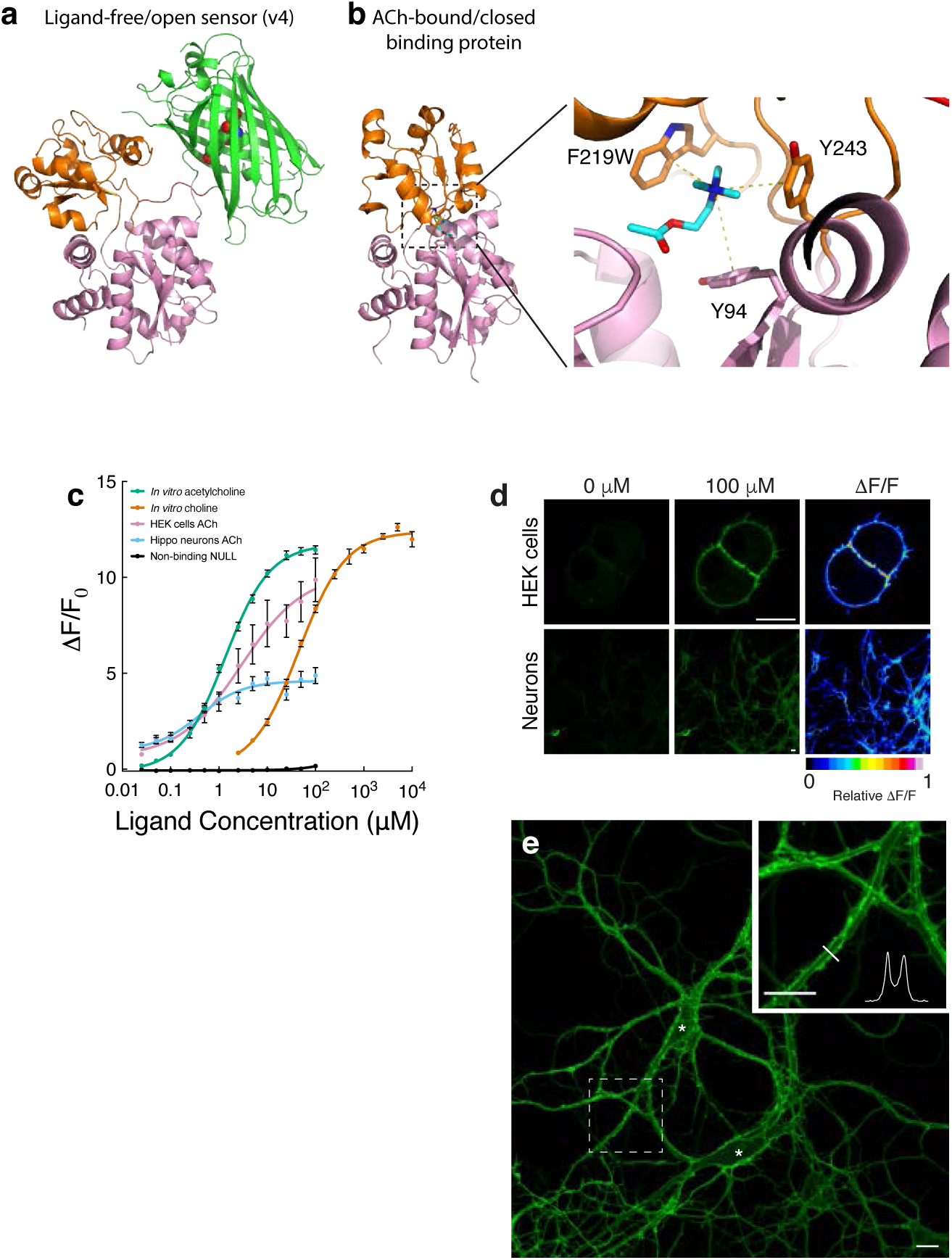
Sensor structure and *in vitro* characterization. (**a**) Overall iAChSnFR0.4 structure as determined by x-ray crystallography (PDB ID: 6URU). *Orange and pink*, periplasmic binding domain (PBP), colored to illustrate Venus-flytrap-like domain arrangement; *green*, circularly permuted superfolder GFP, with chromophore represented by spheres. Acetylcholine binding occurs in the cleft between the two domains of the PBP. (**b**) Structure of the periplasmic-binding protein component <u>only</u> from iAChSnFR (PDB ID: 6V1R), with zoom in of the binding pocket and residues forming cation-π interactions with acetylcholine. Aromatic residues Tyr94, Trp219, and Tyr243 correspond to residues labeled β, ε, and η by (ref. 34), respectively. (**c**) Titrations of purified iAChSnFR protein (soluble *in vitro*), and on the surface of cultured HEK293 cells and neurons. Data are means ± std.dev., n=3 for each experiment, fit with a sigmoidal dose-response curve. (**d**) Representative images of HEK293 cells and hippocampal neuron cultures in buffer alone, with 100 µM acetylcholine, and transformation of ΔF/F as a heat map. Scale bar, 200 µm. (**e**) Image of hippocampal neuron cultures infected with AAV2/1.*hSynapsin1*.iAChSnFR. For two neurons, somata are indicated by asterisks. Inset, magnification of boxed region. Plotted profile is the fluorescence intensity across the indicated line. Scale bars, 200 nm.

### In vitro characterization

Stopped-flow kinetic analysis of purified iAChSnFR shows k_on_ = 0.62 µM^-1^s^-1^ and k_off_ = 0.73 s^-1^ (**Supp. Fig. S8**). pH titrations indicated an apparent pK_a_ of 7.3 in the acetylcholine-bound state and 9.0 in the ligand-free state (**Supp. Fig. S9**). Fluorescence lifetime measurements showed no significant differences between the apo and ligand-bound forms (**Supp. Fig. S10**). Spectroscopy showed excitation and emission similar to other GFP-based sensors (**Supp. Fig. S11**). Introducing mutations from Venus^37^ yielded a yellow sensor y-iAChSnFR, with two-photon excitation compatible with 1030 nm fiber lasers (**Supp. Fig. S11**)^38^.

When cloned into a derivative of the pDisplay vector (Invitrogen) containing an IgG secretion sequence and a single transmembrane helical domain, and transiently expressed in HEK293 cells, iAChSnFR localizes almost exclusively to the plasma membrane, with very little intracellular accumulation (**Fig. 1d**). The sensor responded robustly to perfusion using an HBSS-acetylcholine solution, with apparent *K*_d_ = 2.9 ± 1.6 µM, and (ΔF/F)_max_ = 10 ± 1.4 (**Fig. 1c**).

When cloned into an adeno-associated virus (AAV) vector, and expressed under the human *Synapsin-1* promoter in cultured dissociated rat hippocampal neurons (**Fig. 1e**), iAChSnFR showed an apparent *K*_d_ of 0.4 ± 0.4 µM with (ΔF/F)_max_ = 4.5 ± 0.2 (**Fig. 1c**).

### Validation of iAChSnFR in cultured brain slices

We first expressed iAChSnFR in CA1 pyramidal neurons in cultured rat hippocampal slices with a lentiviral construct (**Supp. Fig. S12a**). Two-photon imaging showed iAChSnFR expression throughout, localized to the plasma membrane of somata, dendrites, and spines of CA1 neurons (**Supp. Fig. S12b-c**). We then expressed iAChSnFR in CA1 pyramidal neurons in cultured rat hippocampal slices using Sindbis virus (**Supp. Fig. S13a; Methods**). Epifluorescence imaging showed that 500-ms puffs of ACh evoked robust iAChSnFR responses, whereas puffs of artificial cerebrospinal fluid (aCSF) vehicle did not (**Supp. Fig. S13b-d; Movie S1**). As a second control, we expressed iAChSnFR-NULL in other hippocampal slices; no responses to ACh puff were seen (**Supp. Fig. S13c-d**). iAChSnFR also responded to oxotremorine (mAChR agonist^39^) and nicotine, with the latter exhibiting slower apparent kinetics (**Supp. Fig. S13b,e-h; Movie S2**).

We next used the same preparation to study the responses of CA3 neurons by simultaneous imaging and patch-clamp electrophysiology (**Supp. Fig. S14a**). CA3 neurons express high levels of cholinergic receptors^40^ and respond robustly to ACh puffs (**Supp. Fig. S14b**). Patch-clamp recordings showed that ACh-induced inward currents rise to a peak of hundreds of pA in a few tens of ms, then decrease with a time constant of several hundred ms, as expected from desensitization of nAChRs. Importantly, the growth of ACh-induced iAChSnFR fluorescence waveform matched growth of the ACh-induced currents, showing that iAChSnFR has the temporal resolution to reveal ACh presence as it activates nAChRs (**Supp. Fig. S14d**). A later phase of ACh-induced current grows and decays with kinetics of tens of seconds and has peak amplitudes of just a few pA, presumably due to mAChR activation^41^. Cells expressing iAChSnFR, and control, non-fluorescent cells nearby, showed no significant differences in their voltage-gated properties at rest (**Supp. Fig. S14b-c**). Furthermore, iAChSnFR-expressing cells showed statistically identical latencies from ACh puff to electrophysiological and fluorescence responses (**Supp. Fig. S14d**), indicating that iAChSnFR does not appreciably buffer ACh at this expression level and detects ACh as rapidly as endogenous cholinergic receptors. Notably, iAChSnFR fluorescence responses had a signal-to-noise ratio (SNR) > 50, at least as great as that of the ACh-induced currents (**Supp. Fig. S14e**). In response to repeated ACh puffs, iAChSnFR responses were constant, while the faster phase of ACh-induced currents ran down (**Supp. Fig. S14f-g**), as expected from many previous studies of nAChR desensitization^41^. ACh-induced currents had the same amplitude, latency, SNR, desensitization, rise time and decay time in iAChSnFR-expressing and control non-expressing neurons, indicating that expressed iAChSnFR neither exerts a kinetic buffering effect on ACh, nor affects ACh-induced currents on its own, under these conditions.

### Imaging properties of cholinergic transmission in acute brain slices

The sensor was then expressed in acute mouse hippocampal brain slice by AAV transduction. In experiments with electrical stimuli on CA1 pyramidal cells, both tetrodotoxin (TTX) and cobalt ions (Co^2+^), which respectively block voltage-gated Na^+^ and Ca^2+^ channels, and thus vesicular transmitter release, eliminated iAChSnFR signals, confirming that the sensor is detecting vesicular ACh release (**Supp. Fig. S15**). In acute slices of medial entorhinal cortex (MEC, **Fig. 2a**)– a region rich in cholinergic projections from nucleus basalis (NB)^5^– iAChSnFR on the surface of layer 2 (L2) stellate cells responds to ACh released by electrical stimulation of axons of NB neurons in layer 1 (**Fig. 2b**). Stimuli delivered at 10-second intervals resulted in essentially identical waveforms with a 10-90% rise time of ∼80 ms, a peak of ∼10% fluorescence increase, and a decay time constant of ∼1.9 sec (**Fig. 2b,d**). This demonstrates both reliable cholinergic transmission from basal forebrain, and reliable iAChSnFR response to release events at modest stimulus rates. At higher stimulus rates, time-dependent rundown of iAChSnFR signal occurred, providing perhaps the clearest evidence to date that in this preparation, decreased pre-synaptic ACh release underlies the observed frequency depression in post-synaptic cholinergic currents^42^ (**Fig. 2c, Movie S3**). Electrical stimulus-evoked iAChSnFR signals were often restricted to cellular sub-regions (**Movie S3**).

**Fig. 2.**
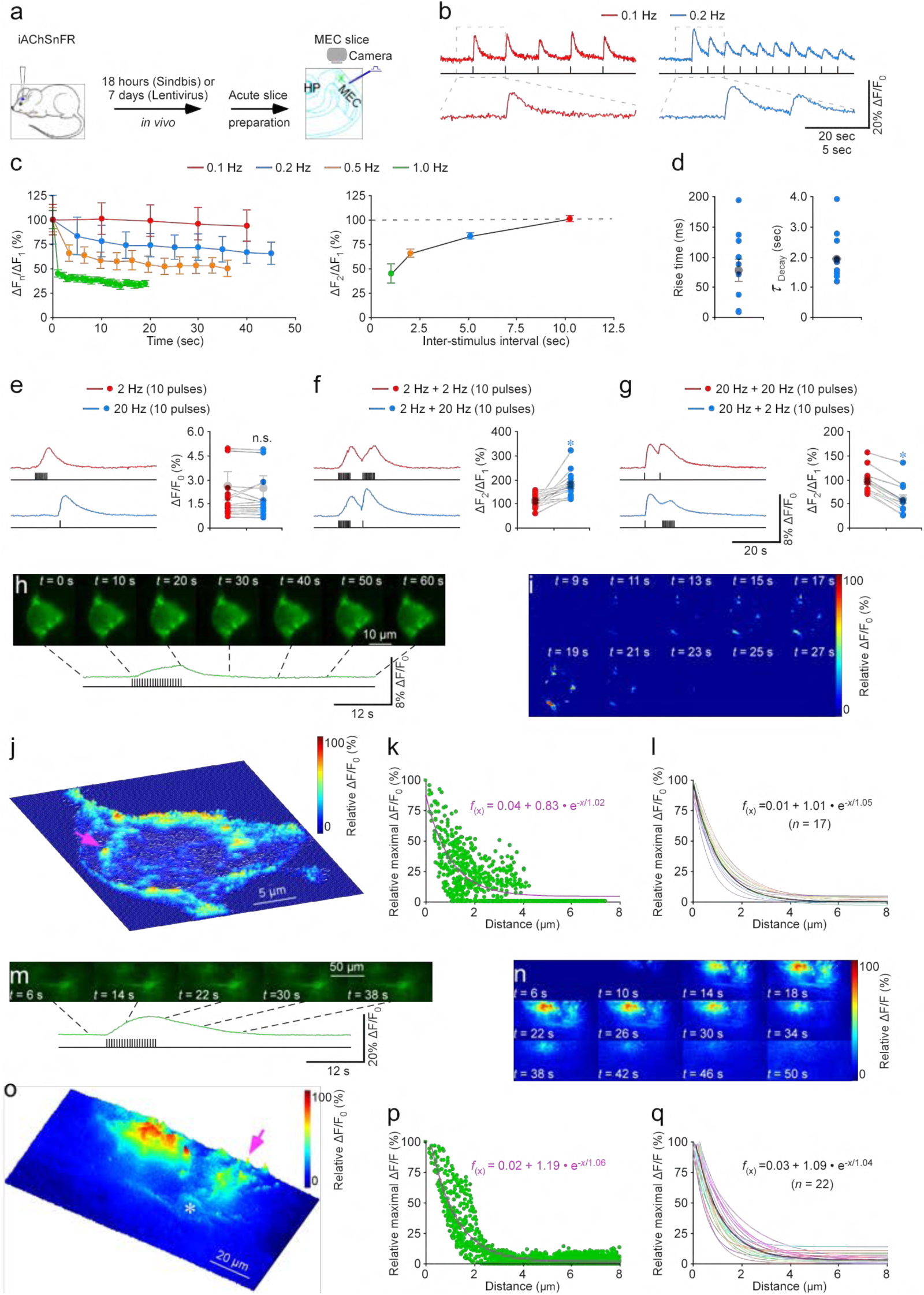
iAChSnFR illustrates endogenous MEC cholinergic transmission. (**a**) Schematic drawing of expression and stimulation-imaging experiments acute mouse MEC slices. (**b**) Representative fluorescence responses to trains of electrical pulses at 0.5 Hz. Responses in the dashed boxes are expanded below. (**c**) Peak fluorescence responses to trains of electrical pulses (normalized to the first response in each train). Red, 0.1 Hz, 5 pulses (n=13 from 12 animals); blue, 0.2 Hz, 10 pulses (n=15 from 12 animals); orange, 0.5 Hz, 12 pulses (n=10 from 10 animals); green, 1.0 Hz, 20 pulses (n=10 from 10 animals). Right, paired-pulse ratio for ACh release (peak ΔF_2_)/(peak ΔF_1_) *vs.* inter-stimulus interval. Dashed line indicates 100%. (**d**) Values for the 10-90% rise time and decay time constant of fluorescence waveforms for the first stimulus (rise time: 77±18 ms; *n*=11 from 10 animals; decay time constant: 1.92±0.25 s; *n*=11 from 4 animals). (**e-g**) Analysis of pulse trains at higher stimulus frequencies. (**e**) Left, summation of fluorescence responses during trains of 10 electrical pulses at 2 (red) or 20 Hz (blue). Right, peak values after trains (2 Hz: 2.6±0.9%; 20 Hz: 2.5±0.7%; *Z*=-0.604; *p*=0.55; *n*=19 from 11 animals). (**f**) Left, fluorescence responses after two consecutive 10-pulse trains. Ten pulses at 2 Hz were followed, after a 5 sec pause, by a train of 10 pulses at 2 Hz (red) or 20 Hz (blue). Right, paired-train ratio for 2+2 Hz and 2+20 Hz stimuli (2+2 Hz: 1.1±0.08; 2+20 Hz: 1.8±0.1; *Z*=3.296; *p*=0.001; *n*=14 from 10 animals). (**g**) Left, responses to a 10-pulse train at 20 Hz followed, after a 5 sec pause, by a 10-pulse train at 2 Hz (red) or 20 Hz (blue). Right, paired-train ratio for 20+2 Hz and 20+20 Hz stimuli (20+20 Hz: 1.0±0.07; 20+2 Hz: 0.6±0.09; *Z*=-3.059; *p*=0.002; *n*=12 from 10 animals). Large gray dots indicate average responses and asterisks indicate *p*<0.05 (Wilcoxon tests) in **e**-**g**. Scale bars in **g** apply to **e**-**g**. (**h**) Fluorescence snapshots for an iAChSnFR-expressing stellate cell responding to local electrical stimuli. (**i**) Temporal and spatial dependence of ΔF/F_0_. Images are Gaussian filtered to enhance visualization of release hot spots. (**j**) Three-dimensional spatial and temporal ΔF/F_0_ profile of the iAChSnFR-expressing cell, shown as heat map. An isolated release site chosen for further analysis is indicated by pink arrow. (**k**) Fluorescence responses at the isolated release site indicated by pink arrow in **j** at ∼180 nm/pixel resolution over 60-second imaging period creates a plot of relative maximal ΔF/F_0_ of each pixel against its distance to the pixel of largest maximal ΔF/F_0_. Fitting with a single exponential decay function (pink line) yields an estimated volume spread length constant of 1.02 µm. (**l**) Superimposed fits for exponential spread from 17 putative single release sites; the average length constant of 1.05±0.05 µm for cholinergic transmission at MEC (n=17 sites on 8 neurons from 8 animals). (**m-q**) As (h-l), for an astrocyte. In **o**, note that for the astrocyte, the predominant responses occur at distal processes, with one isolated release site indicated by pink arrow, not at the cell body (white asterisk). In **p**, fit is 1.06 µm; in **q**, fit is 1.04±0.08 µm (*n*=22 from 10 neurons from 8 animals). Note the average single exponential decay function fitting curves in black in **l** and **q**.

In awake animals, basal forebrain cholinergic neurons may alternate between tonic (∼0.5-4 Hz) or bursts of higher-frequency (∼theta rhythmic) action potentials^43, 44^. In trains of 10 stimuli at > 2 Hz, individual responses in slice were poorly resolved, and instead summed during the trains. When the train ended, fluorescence began to decay within a few sec (**Fig. 2e**). When 2 Hz and 20 Hz trains were delivered alternately, the 20 Hz trains produced a higher (ΔF/F)_max_ (**Fig. 2f,g**).

We next employed nanoscopic imaging and analysis algorithms^45^ (**Methods**) to visualize the distribution of iAChSnFR fluorescence changes on the surface of stellate cells at the micron scale (**Fig. 2h-i, Movie S4**). Electrical stimulation elicited ACh release at multiple, resolvable loci (**Fig. 2j**); release sites often clustered together, although individual release sites were sometimes seen. The distribution of pixel-wise relative (ΔF/F)_max_ revealed the spatial spread of ACh after release (**Fig. 2k**); fitting with a single exponential showed that ACh spread with a length constant of ∼1.0 µm (**Fig. 2k-l**).

Bath application of the non-competitive, reversible AChE inhibitor donepezil (Aricept^®^; does not affect iAChSnFR function, see above) increased iAChSnFR responses and slowed down decay kinetics resulting from electrical stimulation (**Supp. Fig. S16**). This indicates that AChE is, unsurprisingly, at least partially, responsible for terminating the observed sensor waveform.

Finally, we explored cholinergic signaling onto astrocytes. ACh receptors are highly expressed at distal processes^46^, but the functional role of such receptors is poorly understood. First, iAChSnFR was expressed on the surface of astrocytes in acute mouse striatal brain slice with AAV under the glial fibrillary acidic protein (*gfaABC_1_D*) promoter (**Supp. Fig. S17a,b**). Electrical stimulation and ACh application (**Supp. Fig. S17c,d,g**) showed clear responses, with ACh puff much larger. Application of TTX ablated ACh release (**Supp. Fig. S17e**), whereas donepezil application strengthened it (**Supp. Fig. S17f**). Responses at somata and processes were similar (**Supp. Fig. S17d**).

Next, we expressed iAChSnFR on astrocytes in acute MEC brain slice using lentivirus. Nanoscopic imaging showed prominent electrical stimulus-evoked ACh release at distal processes, but not somata (**Fig. 2m-o, Movie S5**). Single, isolated release sites were also seen on entorhinal astrocytes (**Fig. 2o**), with (ΔF/F)_max_ ∼2-fold larger than those on entorhinal neurons (**Supp. Fig. S18**). As before, an ACh spread length constant of ∼1.0 µm was seen (**Fig. 2p-q**). Importantly, (ΔF/F)_max_ of iAChSnFR-expressing neurons and astrocytes had only a weak negative correlation with basal fluorescence F_0_ (**Supp. Fig. S19**), suggesting that ΔF/F_0_ responses are largely independent of iAChSnFR expression levels (cf.^47^).

Finally, we sought to image iAChSnFR activity while directly recording electrophysiological signals of cholinergic signaling in an intact circuit. For this, we co-expressed iAChSnFR together with the G protein-coupled inwardly-rectifying potassium channel GIRK2 (*a.k.a.* K_ir_3.2), which couples signaling through GPCRs – including the mAChR4 (“M_4_”) expressed on striatal dorsal medium spiny neurons (dMSNs) ^48^ – to a distinctive K^+^ current, validated *in vivo* in dMSNs^42, 49^. After successful co-transduction of dMSNs with AAV2/9-*hSynapsin1*.iAChSnFR and AAV2/9-*hSynapsin1*.tdTomato.T2A.mGIRK2-1-A22A.WPRE.bGH, we directly compared iAChSnFR imaging with electrophysiological recordings in detecting endogenous ACh signals in acute striatal slices (**Supp. Fig. S20a**). In response to the same electrical stimulation (20-30 μA; 0.5 ms), iAChSnFR fluorescence responses had faster activation kinetics, but slightly slower decay, compared to presumptive M_4_-mediated inhibitory post-synaptic currents (IPSCs; **Supp. Fig. S20b-g**). Application of the voltage-gated Na^+^ channel blocker tetrodotoxin (TTX; 500 nM) or perfusion of Ca^2+^-free aCSF eliminated both iAChSnFR responses and M_4_-mediated IPSCs (p < 0.01, Wilcoxon matched-pairs signed-rank test, n = 10 for both) (**Supp. Fig. S20d,f**).

### In vivo imaging of worm NMJ activity

Most NMJs in the nematode *Caenorhabditis elegans* signal with ACh, although some use GABA^50^. To directly image ACh function at the worm NMJ, we expressed iAChSnFR0.7 (with the signal sequence replaced by a worm-specific sequence; **Methods**) post-synaptically in body-wall and egg-laying muscles using the muscle-specific promoter *myo-3*. iAChSnFR expressed well in both muscle types (**Fig. 3a**), and immobilized worms were imaged on a confocal microscope. We first imaged a small region of the dorsal nerve cord, where motor neurons synapse onto body-wall muscles; iAChSnFR fluorescence intensity was generally higher along the nerve cord, relative to the muscle arms or body (**Fig. 3a**). We observed iAChSnFR increases at punctate spots reminiscent of worm presynaptic structures (**Fig. 3b**). In egg-laying muscles (vm1/2 of the vulva), iAChSnFR increases were observed at the outer muscle margins, presumably reflecting the sites of cholinergic innervation by VC motor neurons (**Fig. 3c**). Spontaneous iAChSnFR transients occurred with a frequency of ∼0.05 Hz (**Fig. 3d**), similar to activity patterns of VC and HSN motor neurons during egg-laying behaviors^51^. Thus, iAChSnFR can read out motor neuron activity at both body-wall and egg-laying NMJs in living worms, with both spatial and temporal activity consistent with literature reports.

**Fig. 3.**
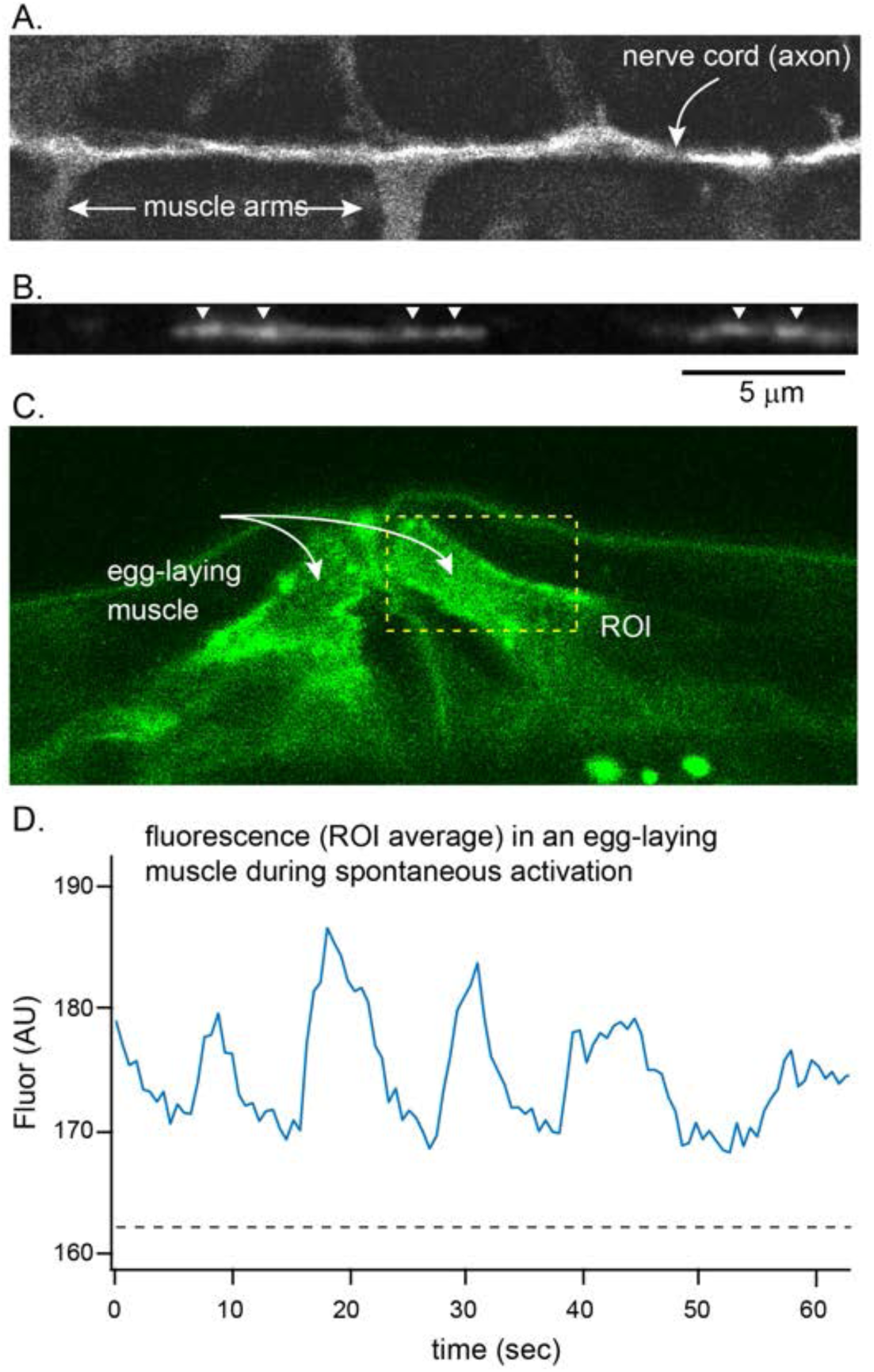
iAChSnFR responses at the *C. elegans* neuro-muscular junction. (**a**) Longitudinal image of a small region of the dorsal nerve cord, where motor neurons synapse onto body-wall muscles; also shown are the muscle arms. (**b**) iAChSnFR0.7 fluorescence increases at puncta reminiscent of presynaptic structures (arrowheads). (**c**) At the vulva, egg-laying muscles (vm1/2) expressing iAChSnFR0.7. Region of interest (ROI) quantified in (d) indicated. (**d**) Fluorescence changes at the ROI in (c).

### In vivo detection of acetylcholine release in zebrafish

Zebrafish have two large Mauthner cells (one in each side of the hindbrain) that control the fast escape reflex^52, 53^. Mauthner axons excite both motor neurons and inhibitory interneurons, ensuring that contralateral muscles relax^52^. Primary motor neurons in the spinal cord are known to receive cholinergic transmission from these cells at their ventral processes^54, 55^.

We established a fictive swimming preparation using an immotile zebrafish line (**Methods**). We replaced the mammalian IgG secretion sequence in iAChSnFR with the signal peptide from zebrafish Nodal-related 1 (Ndr1, squint). We sparsely expressed iAChSnFR in spinal motoneurons by microinjection of 10xUAS:iAChSnFR plasmid in a transgenic line expressing GAL4 under the VAChT promoter (TgBAC(*vacht*:GAL4)). We imaged primary motoneurons in spinal cord during stimulus-induced fictive escape to see cholinergic inputs to motoneurons including that from a Mauthner cell (**Fig. 4a,b**). Changes in fluorescence were observed in 4 cells (in 3 fish). The greatest change in fluorescence was ∼7%, observed in an axon proximal to the soma, while other sub-cellular locations showed smaller changes (**Fig. 4c,d**).

**Fig. 4.**
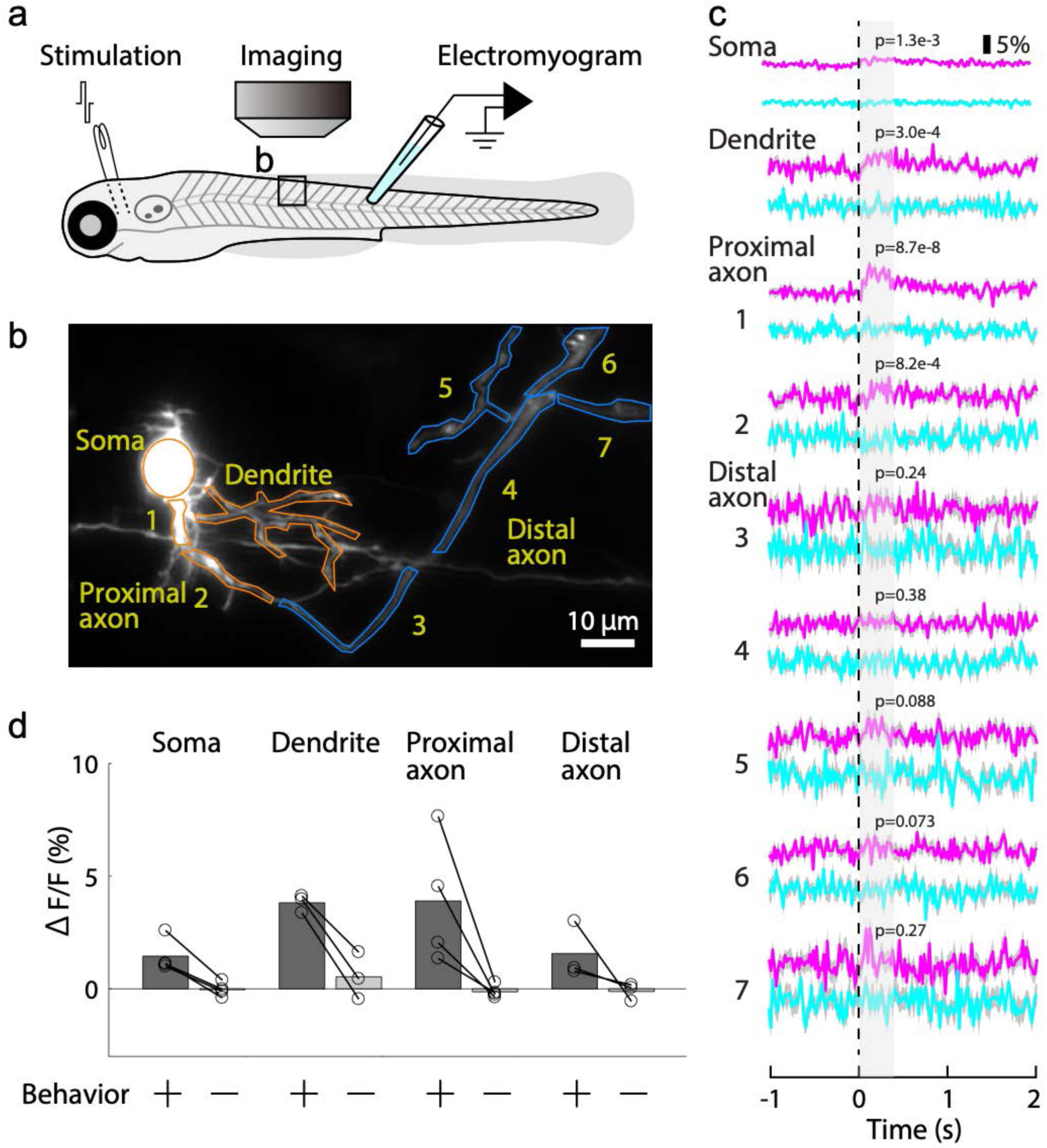
Behavior-related iAChSnFR responses in larval zebrafish. (**a**) Schematic of experimental setup. (**b**) 2-Photon fluorescence image of iAChSnFR-expressing primary motor neuron in the spinal cord. (**c**) *Magenta*, trial-averaged iAChSnFR responses to electrical shock with fictive motor activity observed. *Cyan*, average of 11 waveforms without observed fictive motor activity. *Dashed line*, electrical shock. *Grey*, period for statistical analysis. P-values from n=13 (with motor activity) and 11 (without motor activity) trials, two-sample t-test shown. (**d**) Average peak iAChSnFR responses during behavior, at several somatic, dendritic, and axonal ROIs (n = 4 cells from 3 fish).

### In vivo imaging of acetylcholine in Drosophila melanogaster

In flies and many other invertebrates, ACh, rather than glutamate, is the predominant excitatory neurotransmitter. Although the entire *Drosophila melanogaster* brain has been recently mapped to the synapse level^56^, the details of synaptic communication *in vivo* are not yet fully understood. Combined with myriad genetic and molecular tools already available, visualization of ACh in *Drosophila* would further facilitate mechanistic understanding of neuronal and circuit physiology in this model system.

*Drosophila melanogaster* has about 50 glomeruli in the antennal lobe, a central brain structure similar to the mammalian olfactory bulb and formed by olfactory sensory neuron (OSN) axons, olfactory projection neuron (OPN) dendrites and local interneurons^57, 58^. We replaced the mammalian IgG secretion sequence from iAChSnFR with the signal peptide from *Drosophila* immunoglobulin binding chaperone protein (BiP)^59^, and expressed it in OPNs under the *GH146*-GAL4 driver^57^. We recorded the fluorescence signal of iAChSnFR in OPN dendrites with two-photon excitation in whole-mount adult *Drosophila*, while stimulating their antennae with odorants. iAChSnFR reported acetylcholine release in response to individual odor trials (**Fig. 5a**) with (ΔF/F)_max_ of 43% ± 4.5% (mean ± s.e.m., *n* = 4 flies, 6 trials per fly) with a mean signal-to-noise ratio of 6.88 ± 0.86.

**Fig. 5.**
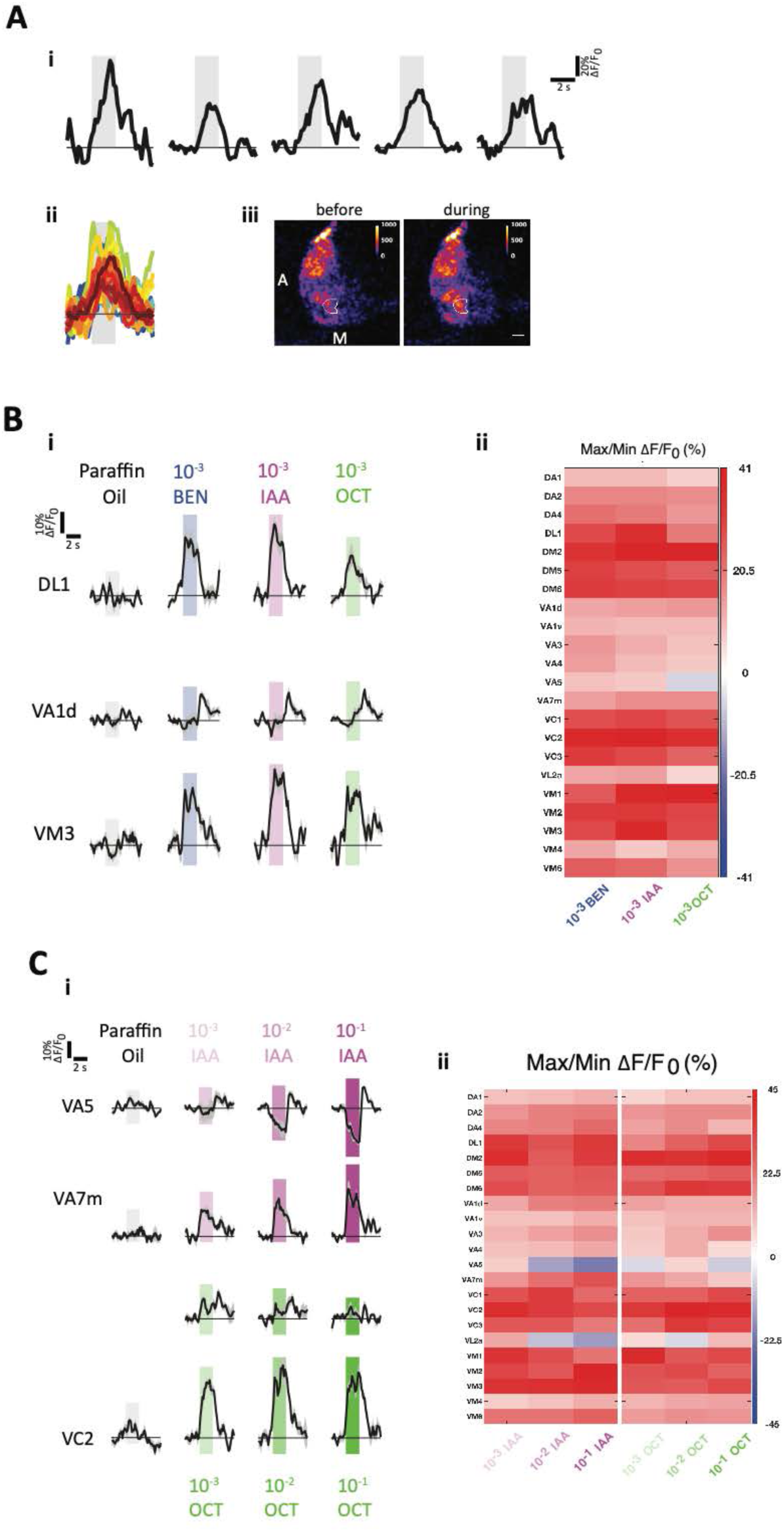
iAChSnFR responses to odor stimulation in *Drosophila melanogaster*. (**a**) iAChSnFR can report single-trial acetylcholine release in response to odor stimulation in the antennal lobe olfactory projection neuron dendrites. i) Five example traces of single-trial responses to 10^-2^ 3-octanol in the VC2 glomerulus. Traces show 2 sec before odor, 2 sec of odor stimulation (shaded), and 3 sec after odor offset. Scales for time and ΔF/F_0_ are shown at top right. ii) VC2 response traces from all individual trials (24 trials from 4 flies) with 10^-2^ 3-octanol stimulation. Each colored line represents an individual trial. Same scales for time and ΔF/F_0_ as in Ai. iii) Pseudo-colored images of the right antennal lobe before and during 10^-2^ 3-octanol stimulation. Each image is the average intensity projection of 8 frames (1.2 s) either before or during stimulation. The color scales for the fluorescence intensity are shown in the top right corners. Left is anterior, down is medial (“A” and “M” in the left image). The VC2 glomerulus is identified from the average intensity projection of all trials and shown with the white outline. Scale bar in the right image is 5 μm. (**b**) iAChSnFR reports varied acetylcholine received across glomeruli and time, depending on odor identity. i) Mean response traces of 3 example glomeruli to paraffin oil (negative control), 10^-3^ benzaldehyde (BEN), 10^-3^ isoamyl acetate (IAA), and 10^-3^ 3-octanol (OCT). N = 4 flies for each odorant-glomerulus combination. Gray shaded envelops around the traces indicate standard errors of the mean. Rectangular color shaded regions indicate the periods of odor stimulation. Scales for time and ΔF/F_0_ are shown at top left. ii) Heat map of the maximum responses across glomeruli and odorants. Positive values are for excitatory responses, negative for inhibitory. Color scale for ΔF/F_0_ (%) is to the right. (**c**) iAChSnFR responses demonstrate varied tuning profiles across odor concentrations and glomerulus identity. i) Mean response traces of 3 example glomeruli to paraffin oil (negative control) and increasing concentrations of isoamyl acetate (10^-3^, 10^-2^, 10^-1^ IAA) and 3-octanol (10^-3^, 10^-2^, 10^-1^ OCT). N = 4 flies for each odorant-glomerulus combination. Gray shaded envelops around the traces indicate standard errors of the mean. Rectangular color shaded regions indicate the periods of odor stimulation. Darker shades indicate higher odor concentrations. Scales for time and ΔF/F_0_ are shown at top left. ii) Heat map of the maximum responses across glomeruli and odor concentrations. Positive for excitatory responses, negative for inhibitory. Color scale for ΔF/F_0_ (%) is to the right.

OPNs encode odor identity by activating different combinations of olfactory glomeruli in response to odorants^60, 61^. To determine if iAChSnFR can capture the combinatorial complexity of odorant-induced glomerular activation, we analyzed iAChSnFR responses in 22 glomeruli to 3 different odorants diluted to 0.1% in mineral oil: benzaldehyde (BEN), isoamyl acetate (IAA), and 3-octanol (OCT). iAChSnFR had variable (ΔF/F)_max_ across different glomeruli and odorants, ranging from −6% (BEN in VA1d) to 40% (IAA in VM3) (**Fig. 5b**). Rates of change were also variable, including a delayed response in VA1d (**Fig. 5b(i)**, second row). Across the 22 glomeruli, each odorant elicited a different response signature (**Fig. 5b(ii)**), corroborating previous reports from calcium indicators and electrodes^61, 62^.

Since varying concentrations of odorants can be represented as varied amplitudes and combinations of glomeruli responses, we further analyzed the iAChSnFR responses from the 22 identified glomeruli to 2 odorants at 3 concentrations spanning 2 orders of magnitude (0.1%, 1%, and 10%; **Fig. 5c**). As with calcium imaging and electrode recordings^61, 63^, varying odor concentration resulted in a wide range of iAChSnFR response amplitudes across different combinations of identified glomeruli. Fluorescence was positively correlated with IAA concentrations in some glomeruli, but negatively in others (as low as −21% below baseline, **Fig. 5c(i)**). This is consistent with previous reports of OSN inhibition by certain odors^64^, and demonstrates the utility of iAChSnFR in reporting not only increases, but also decreases, in acetylcholine release.

Overall, we demonstrate that iAChSnFR expressed in the *Drosophila* brain is both sensitive enough to report single-trial responses and has sufficient dynamic range to report complex odor coding in terms of the combinatorial code, including both excitation and inhibition. We believe it will be an invaluable tool in the investigation of synaptic and circuit dynamics *in vivo*.

### In vivo detection of acetylcholine in mouse motor cortex

Acetylcholine released in mammalian cortex produces biphasic responses in excitatory pyramidal cells (PCs) comprising fast hyperpolarization followed by slow depolarization^65, 66^. This fast inhibition is mediated by both nAChRs and mAChRs, which increase the excitability and firing rate of PC dendrite-targeting GABAergic interneurons, primarily somatostatin (*Sst*)^+^ and vasoactive intestinal polypeptide (*VIP*)^+^ cells^67, 68^. Slow depolarization is mediated by mAChR closure of M-type (KCNQ, *a.k.a.* Kv7) potassium channels in pyramidal neurons, enhancing their excitability^5, 69^. Motor cortex receives active cholinergic projections originating from the NB^70^, here revealed by calcium imaging in mouse primary forelimb motor cortex (M1) with GCaMP6s (ref. ^71^; **Supp. Fig. S21**). Further experiments showed that direct electrical stimulation of NB induces a fast activation of *Sst*^+^ interneurons in layer 2/3 (L2/3) followed by delayed (5.5 s later) activation of pyramidal cells (**Supp. Fig. S22**). Thus, M1 provides a suitable platform for testing iAChSnFR responses in cell types responsible for biphasic cortical cholinergic signaling.

To examine acetylcholine release from NB onto cortical pyramidal cells and interneurons (**Fig. 6a**), M1 of *Sst*-Cre mice (for interneurons) was infected with Cre-dependent AAV expressing iAChSnFR (AAV2/1.*CAG*.FLEX-iAChSnFR), and M1 of wild-type mice (for pyramidal cells) was infected with AAV2/1.*hSynapsin1*.iAChSnFR. Two weeks after AAV infection, *Sst*^+^ neurons showed strong fluorescence in both pyramidal cells and *Sst*^+^ neurons, with good membrane localization (**Supp. Fig. S23a,b**). Awake mice were head-restrained and positioned on a free-floating, linear treadmill (**Methods**) while iAChSnFR fluorescence was imaged during both quiet resting and forward running. Two-photon imaging under resting conditions detected fluorescence transients in subsets of layer 2/3 (L2/3) dendritic segments (**Fig. 6b-e**). Fluorescence decay appeared to be faster than the upper limit of our acquisition scan speed (10 Hz). (ΔF/F)_max_ of these transients was higher during running than resting (53% ± 3% *vs*. 37% ± 2%, *P* < 0.001, two-way ANOVA followed by Bonferroni’s test, **Fig. 6d**). Local application of a cholinesterase inhibitor (100 µM eserine, ESR) increased (ΔF/F)_max_ during both activities (**Fig. 6d**).

**Fig. 6.**
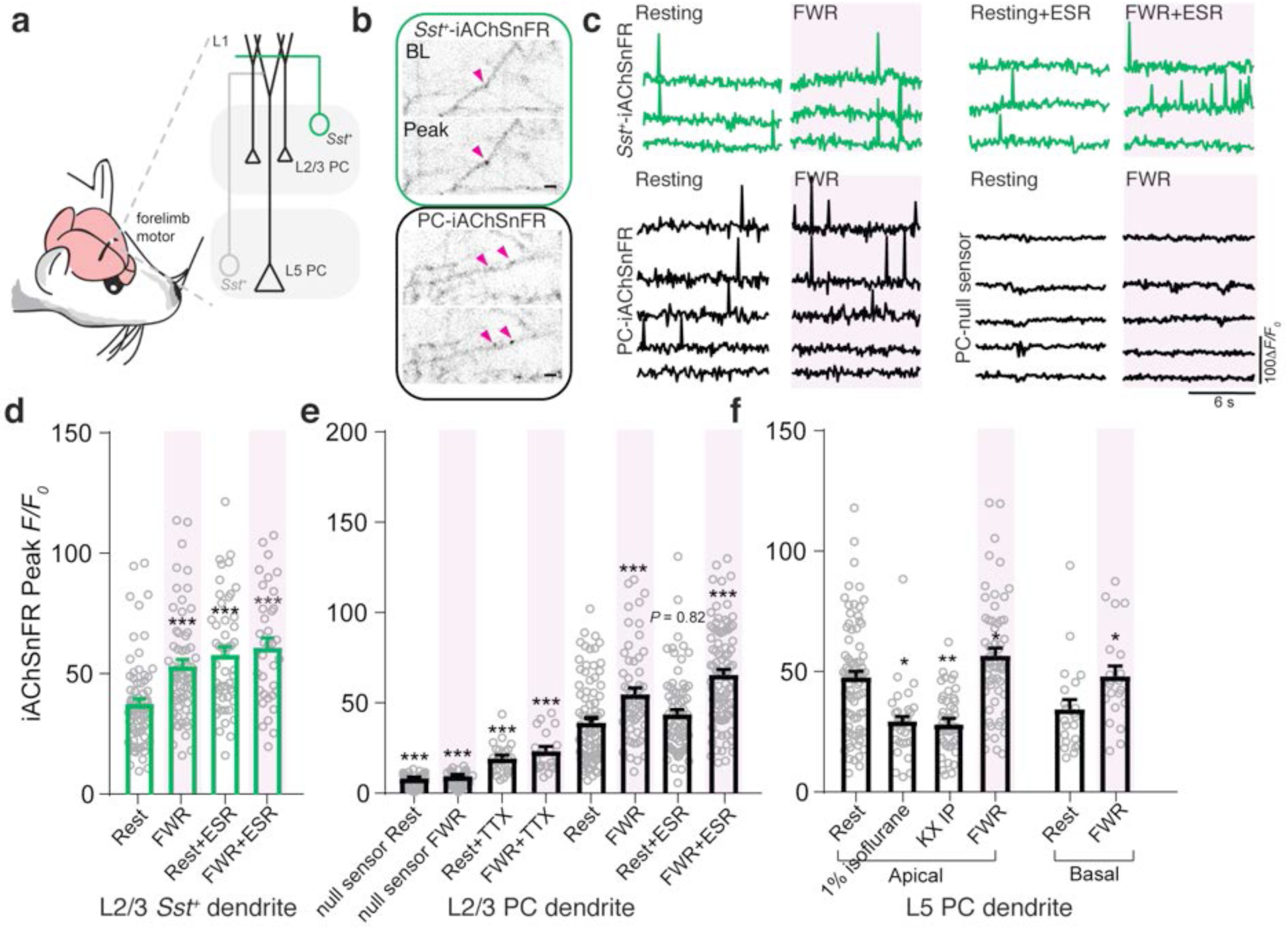
iAChSnFR responses in Somatostatin interneurons and pyramidal neurons in mouse motor cortex. (**a**) Schematic of iAChSnFR expression in different cell types within M1. (**b**) Representative 2-photon images (display inverted) of L2/3 dendrites of *Sst^+^* neurons (top) and PCs (bottom). Baseline (BL) and peak signals are shown. ROIs indicated by arrowheads. Scale bars, 2 µm. (**c**) Fluorescence traces of iAChSnFR and iAChSnFR-NULL from different ROIs over 12 sec trials during quiet resting, forward running (FWR), and with local application of eserine (ESR). (**d**) Average peak Δ*F/F* iAChSnFR signals from *Sst^+^* apical dendrites during resting and forward running (rest: 37 ± 2%, *n* = 72; run: 53 ± 3%, *n* = 63; *P* < 0.001, t=4.08, df=215), and with local ESR (rest: 58 ± 3%, *n* = 49, *P* < 0.001, t=5.11, df=215; run: 61 ± 4%, *n* = 35, *P* < 0.001; t=5.31, df=215). (**e**) Average peak iAChSnFR signals from PC apical dendrites during quiet resting and FWR (rest: 40 ± 2%, *n* = 86; run: 55 ± 3%, *n* = 58). iAChSnFR-NULL showed minimal signals (rest: 8 ± 1%, *n* = 75, *P* < 0.001, t=18.3, df=269; run: 10 ± 1%, *n* = 73, *P* < 0.001, t=17.2, df=269). TTX addition greatly lowered signals (rest: 19 ± 2%, *n* = 24, *P* < 0.001, t=6.63, df=165; run: 23 ± 2%, *n* = 20, *P* < 0.001, t=5.21, df=165). ESR addition enhanced signals during FWR but not resting (rest: 44 ± 3%, *n* = 74, *P* = 0.82, t=1.09, df=305; run: 65 ± 3%, *n* = 91, *P* < 0.001, t=7.03, df=305). (**f**) Average peak iAChSnFR signals from apical tuft dendrites (rest: 47 ± 3%, *n* = 78; run: 56 ± 3%, *n* = 58, *P* < 0.03, t=2.22, df=134) and basal dendrites (rest: 34 ± 4%, *n* = 22; run: 48 ± 4%, *n* = 21, *P* < 0.02, t=2.33, df=41) of L5 PCs. Anesthesia with 1% isoflurane (29 ± 3%, n = 31, *P* < 0.02, t=2.58, df=158) and ketamine/xylazine (KX) (29 ± 2%, *n* = 44, *P* < 0.01, t=2.89, df=158) weakened iAChSnFR signals. Data are presented as means ± s.e.m. *P < 0.05, **P < 0.01, ***P < 0.001, unpaired Student’s t-test in **f** (basal dendrites) and two-way ANOVA followed by Bonferroni’s test in **d**-**f** (apical dendrites). Representative images and traces from experiments carried out on at least 3 animals per group.

We next examined iAChSnFR responses on L2/3 and layer 5 (L5) PC dendrites (**Fig. 6b,e,f**). In awake, quietly resting animals, we observed numerous transient cholinergic events in L2/3 apical tuft dendrites with (ΔF/F)_max_ of 40 ± 2% (**Fig. 6e**). Forward running significantly increased fluorescence response (55% ± 3%, *P* < 0.001, two-way ANOVA followed by Bonferroni’s test, **Fig. 6e**). These responses, like those observed in *Sst*^+^ dendrites, were distributed across the dendritic shaft and quite focal (<0.5 µm; **Fig. 6b**). Responses were strongly modulated by electrical stimulation of NB (R^2^=0.33, *P* < 0.001, **Supp. Fig. S24c**) and by hindlimb foot shock (no shock: 45% ± 2%; foot shock: 59% ± 4%, *P* < 0.001, unpaired Student’s *t*-test, **Supp. Fig. S24d,e**). iAChSnFR signal increased with strength of NB current injection, from 200-800 μA, indicating a wide dynamic range of both recruitment of ACh release from NB fibers and of iAChSnFR response.

Next, AAV2/1.*CAG*.FLEX-iAChSnFR was injected into L5 in *Rbp4*-Cre mice (L5-specific)^72^. In *Rbp4*-Cre mice, iAChSnFR showed good membrane targeting in all dendritic compartments, including thin branches of the apical tuft, apical trunk, and basal dendrites (**Supp. Fig. S23c**). iAChSnFR fluorescence changes in response to resting and forward running were obvious across the dendritic arbor, even below 450 µm in basal dendrites (**Fig. 6f-right**). General anesthesia decreased peak responses (**Fig. 6f-left**), and eserine enhanced responses (**Fig. 6e**). Notably, iAChSnFR responses detected on both *Sst*^+^ interneurons and pyramidal cells were comparable to those observed with iGluSnFR^23, 24^.

### Kilohertz imaging of cortical acetylcholine release

Cholinergic neurons from the basal forebrain (NB) also project to visual cortex (V1), where ACh signaling regulates attention and visual learning^73^. To visualize dynamics of NB-derived acetylcholine signals with micron spatial and millisecond temporal resolution, we used two-photon Scanned Line Angular Projection (SLAP) imaging^38^. Yellow-shifted y-iAChSnFR was expressed in L2/3 cortical neurons with AAV2/1.*hSynapsin1*.y-iAChSnFR, and an electrode implanted in NB (**Fig. 7a**). Previous work showed that 300 μm below the pia, brief stimulation (1 ms, 500 μA) of NB elicits punctate (<2 μm) and rapid (∼10 ms) iAChSnFR transients, consistent with synaptic ACh release events^38^. Here we compared dynamics of stimulation-evoked ACh transients recorded from neuropil at different cortical depths in response to pulse trains of different lengths (1, 3, or 10 pulses at 20 Hz). Evoked neuropil transients 300 μm below the pia showed sharply phasic responses aligned to each stimulus, superposed on a slowly decaying response (**Fig. 7b-bottom**). This persistent component is consistent with seconds-long evoked transients recorded with functionalized microelectrodes^74^, although such methods have so far lacked the time resolution to detect the sharp component. Signals in superficial layers (<250 μm below the pia) lacked the phasic component (**Fig. 7b-top**), suggesting that ACh is degraded and/or released more slowly in superficial cortex, and demonstrating that iAChSnFR enables precise spatiotemporal analysis of neurotransmission (**Fig. 7c**).

**Fig. 7.**
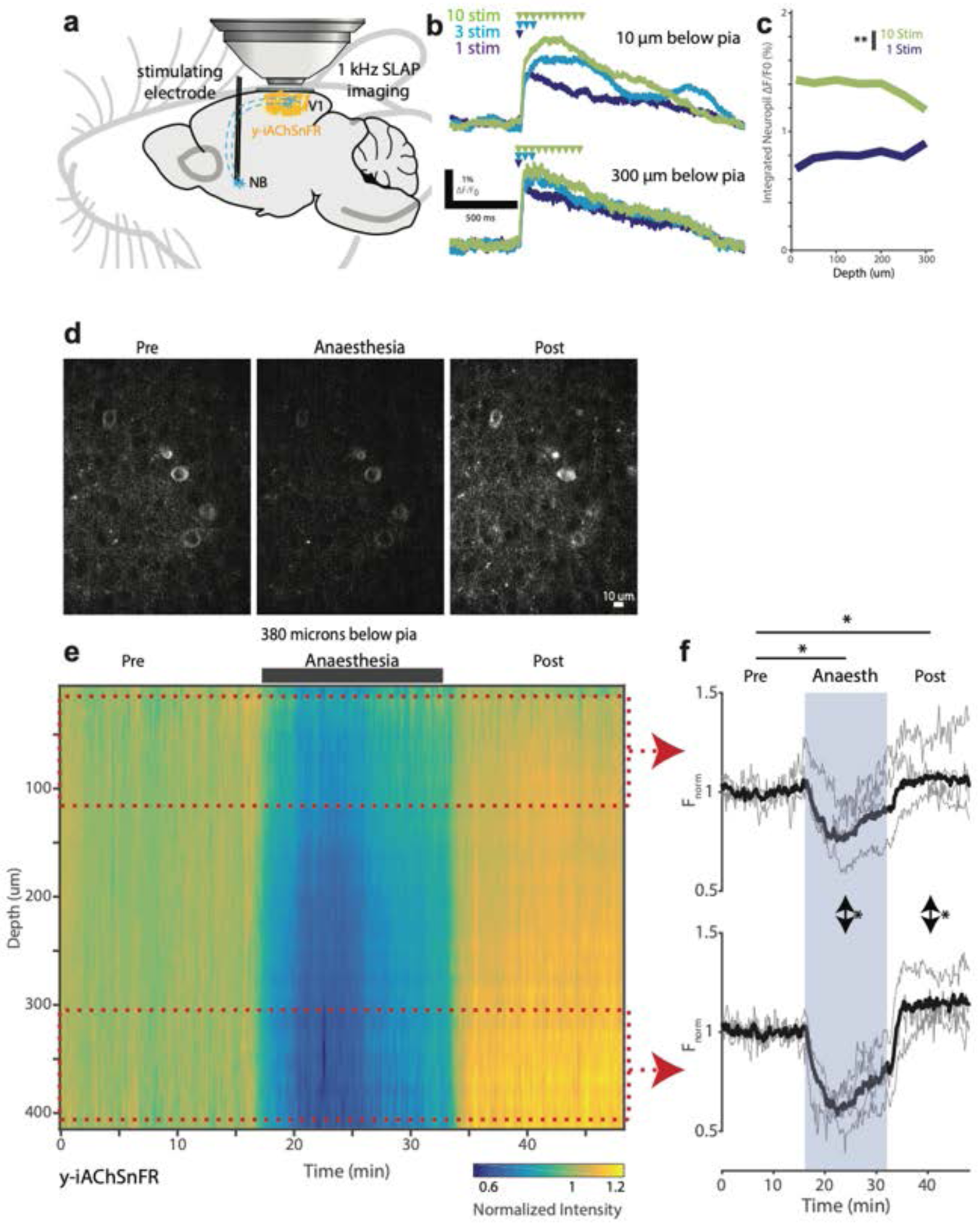
Depth-resolved responses to electrical stimulation of nucleus basalis and anaesthesia. (**a-c**) Depth-dependent y-iAChSnFR responses to electrical stimulation of nucleus basalis (NB). (**a**) Experimental schematic. Visual cortex was densely labeled with AAV2/1.*hSynapsin1*.y-iAChSnFR. y-iAChSnFR signals were recorded from ∼100 μm x 100 μm of neuropil at defined depths using SLAP 2-photon imaging, while electrically stimulating NB. (**b**) Example neuropil responses to trains of 1, 3, and 10 stimuli, 10 µm (top) and 300 µm (bottom) below pia. Arrowheads mark time of stimuli. (**c**) Response amplitudes averaged for the period between 500 and 1000 ms following onset of stimulation. Stimuli consisted of 1 ms, 500 µA pulses at 20 Hz. Mean of *n*=20 trials per condition. (**d-f**) Depth-resolved response of y-iAChSnFR to transient anaesthesia. (**d**) Example image plane (380 μm depth) before, during, and shortly after transient anaesthesia. (**e**) Average y-iAChSnFR intensity in a 156 x 156 x 400 μm deep region of visual cortex, at each Z-plane. 1.5% isoflurane anaesthesia applied from 16 to 32 min. Intensity was normalized to the mean during the waking period, per plane. (**f**) Normalized intensity of the most superficial (**top**) and deepest (**bottom**) 100 μm regions (20-110 μm, 310-400 μm below pia). Shaded region denotes period of anaesthesia. N = 4 sessions shown in gray traces; black trace is mean. *: Mean response for this period differed from awake (“pre”); ↕*: Mean response for this period varied with depth, p<0.05, paired two-sample Student’s *t*-test.

### Depth-resolved recording of anesthesia effects on cholinergic signaling

During anesthesia, cortical ACh levels are known to decrease^75^. However, due to limitations of existing ACh detection methodology (*i.e.* microdialysis), this phenomenon has never been explored with high spatiotemporal resolution. To directly observe ACh changes during anesthesia, we injected AAV2/1.*hSynapsin1*.y-iAChSnFR into V1 and performed volumetric raster imaging (10-410 μm below pia, 12 volumes/min). y-iAChSnFR fluorescence across V1 decreased upon isoflurane application (**Fig. 7d**) and recovered following anesthesia removal, indicating a decrease in cholinergic tone during anesthesia, consistent with reports^76^. Superficial cortical layers showed moderate depression of iAChSnFR signal during anesthesia, while deeper regions showed stronger depression followed by overshoot following anesthesia offset (**Fig. 7e,f**)^76^.

Cholinergic release sites and AChE expression are patterned within the cortex on scales from nanometers to millimeters^77^. iAChSnFR will help address how this patterning drives cholinergic signal dynamics on micron spatial and millisecond temporal scales, and how anesthesia affects these patterns^76^.

### Monitoring cholinergic reward signals in mouse hippocampus

As a final demonstration of the *in vivo* utility of iAChSnFR, we used it to measure another aspect of cholinergic signaling: its contribution to reward signals^78^, specifically in the hippocampus^79^. Hippocampal lesions, particularly in CA1, impair acquisition and performance of operant (*i.e.* reward- and punishment-driven) tasks^5^. The medial septum is the source of most ACh released in the hippocampus, and projects multiple cholinergic fiber types to CA1^80^. Accordingly, iAChSnFR was expressed in CA1 pyramidal cells by viral transduction with AAV2/1.*hSynapsin1*.iAChSnFR (**Fig. 8a**). First, we sought to calibrate iAChSnFR response in this circuit by artificially driving ACh release onto CA1 through electrical stimulation of the medial septum (**Fig. 8a**). Electrical stimulation, at several frequencies, led to robust iAChSnFR signals in CA1 (100 ± 24 average Z-scores above baseline, *n*=3 repeats at each of 4 frequencies in each of 4 mice; **Fig. 8b**).

**Fig. 8.**
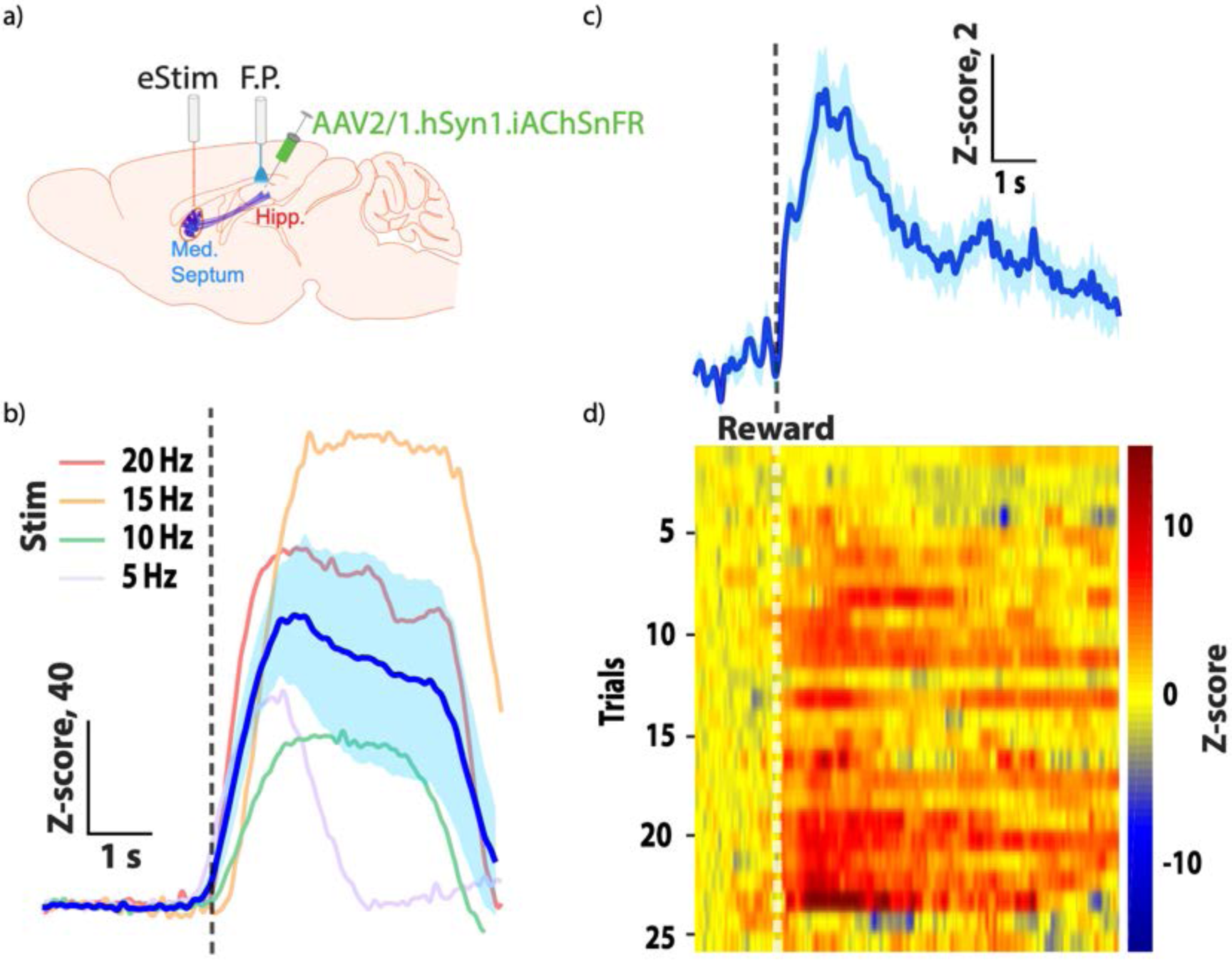
Monitoring hippocampal reward related ACh transmission *in vivo* during an operant task. a) Schematic of the experiment. eStim, electrical stimulation in the medial septum. F.P., fiber photometry in hippocampal CA1 region. b) Response to electrical stimulation. Representative individual Z-scores of iAChSnFR signals following electrical stimulation (twenty 8-msec pulses of 200 µA intensity at 5, 10, 15, or 20 Hz, for 4.0, 2.0, 1.33, and 1.0 sec, respectively) in the medial septum (*n*=3 repeats at each frequency in each of 4 mice). Mean of all responses (dark blue; average of every repeat of every stimulus frequency in every mouse) Z-score ± s.e.m. (light blue shading). c) Mean Z-score ± s.e.m. during a progressive ratio operant task (*n*=26 trials in a single, representative mouse). d) Heatmap illustration of Z-scores on each trial during the operant task.

During behavioral testing, an increase in the fluorescence signal was observed immediately following reward consumption at each prescribed ratio within the progressive ratio operant task. The average peak intensity was 5.1 ± 0.8 (*n* = 26 trials) Z-scores greater than the average baseline signal prior to the reward event (**Fig. 8c**). Importantly, these increases in the fluorescence signal were observed in the large majority of trials (**Fig. 8d**).

## Discussion

Here we design, optimize and exploit a genetically encoded biosensor, iAChSnFR, for the ubiquitous neurotransmitter acetylcholine. The sensor is based on a bacterial choline-binding protein, and it shows no apparent effects on neurons following stable expression in rodent brain slice and in living worms, fish, flies, and mice – both in the brain and at the neuromuscular junction (NMJ). We further created both green and yellow variants, the latter enabling kilohertz frame-rate imaging in mouse cortex *in vivo* with SLAP microscopy^38^. The sensor also allows detection of both increases and decreases of acetylcholine signaling from tonic levels.

The new method for ACh detection is less cumbersome than previous techniques using microdialysis, derivatization and chromatography, or choline electrochemistry; the new method also has superior spatial and temporal resolution. A genetically encoded acetylcholine fluorescent indicator (GACh2.0)^20^ was recently published, based on insertion of circularly permuted GFP into the M_3_R muscarinic acetylcholine receptor. iAChSnFR constitutes an important complementary tool to GACh2.0 (and improved versions: GACh3.0 has about twice the ΔF/F as GACh2.0, with similar kinetics^81^).

We directly compared iAChSnFR and GACh2.0 in side-by-side in acute cortical brain slices. Direct sensor comparison at the same entorhinal stellate neurons shows that GACh2.0 is capable of detecting full cholinergic signals at a low frequency range (≤1 Hz), but not higher frequencies, and it fails to faithfully follow the time courses of signals at all these frequencies (**Supp. Fig. S25a-c**). Analysis confirms that iAChSnFR has several fold faster kinetics, resolving peak responses at ∼1 Hz and summed responses at higher frequencies (**Supp. Fig. S25d**). (The soluble nature of iAChSnFR allows relevant characterization *via* stopped-flow experiments; activation has a time constant of ∼ 140 ms at 10 μM ACh and just a few ms at 1 mM ACh; **Supp. Fig. S8**). iAChSnFR also showed larger (ΔF/F)_max_ than GACh2.0 (**Supp. Fig. S25c,d**). Membrane targeting of both sensors was good, across cell bodies, dendrites, and spines

In general, G protein-coupled receptor-based sensors and microbial periplasmic binding protein-based sensors have different advantages and shortcomings. GPCRs have evolved appropriately high affinity for measuring cognate molecules, while PBPs, which have evolved to help microbes navigate nutrient gradients, frequently have lower affinities. GPCRs also have excellent specificity relative to other endogenous molecules (in this case ACh relative to choline), whereas PBPs often must be tuned by protein engineering (our scaffold protein initially had tighter affinity for choline than ACh). However, GPCRs can be difficult to overexpress, can cause problems when they are^21, 22^, and are more difficult to target to organelles than soluble PBPs. Furthermore, GPCRs and other receptors contain many specific localization/export/retention/binding sequences (*e.g.* endoplasmic reticulum, Golgi, membrane; sequences occur at the N- and C-termini and in intracellular loops) and will not necessarily express well in distantly related organisms. Here, we have shown that PBP-based sensors need no adaptation for successful function in animals as diverse as mouse, fly, fish, and worm – we did tailor the N-terminal secretion peptide for each organism, but this is modular and does not obviously affect sensor performance.

Soluble sensors are readily amenable to purification for use in experiments such as vesicle fusion studies^82^, and also to express in the cytoplasm or organellar lumen to monitor transporter function^83^. PBPs are also readily crystallizable, which facilitates structure-guided optimization, and amenable to high-throughput screening as purified proteins or bacterial lysates, which allows the screening of thousands of mutants simultaneously. Here, we readily solved high-resolution crystal structures of both the full sensor and the mutated binding protein, which fed back into our protein design cycle, allowing us to target specific protein residues and amino acid mutations.

Receptor-based sensors also recognize a host of drug molecules, precluding their simultaneous use with imaging. Meanwhile, PBP-based indicators have few such restrictions, which has allowed use of TBOA, ketamine, NBQX, and other drugs with iGluSnFR^24, 25, 84, 85^, vigabatrin and isoflurane with iGABASnFR^26^, and here, tacrine, donepezil, oxotremorine and eserine with iAChSnFR.

Beginning with a hyperthermophilic binding-protein scaffold was likely critical to successful engineering of iAChSnFR, wherein up to 8 binding-pocket residues were mutated, while still retaining thermodynamic stability, proper sub-cellular targeting, and large fluorescence responses (>10x). In addition to serving as an excellent sensor in its own right, iAChSnFR also provides a promising scaffold for developing additional sensors through targeted mutagenesis. We have developed iAChSnFR into a sensor specific for serotonin by machine learning-guided protein redesign and directed evolution^86^. In parallel with development of iAChSnFR, we also mutated OpuBC to recognize nicotine or the smoking-cessation drug varenicline^36^ (PDB 6EFR). Further experiments produced variants that detect other neural drugs, including the rapidly acting antidepressant S-ketamine^87^, selective serotonin reuptake inhibitor antidepressants, and opioids^88^. In the original OpuBC and related PBPs^89^, and in the better characterized sensors, ligand binding is mediated by cation-π interactions between the protonated secondary, protonated tertiary, or quaternary amine of the ligand and aromatic residues.

In this preliminary report describing the tool, iAChSnFR was able to illuminate a few biological points in various preparations. The high-resolution localization of individual ACh release sites with iAChSnFR lays the groundwork for the study of the fundamental features of acetylcholine transmission, including the transmitter diffusion extent, number of release sites, release pool size, release probability, quantal size and refilling rate^45^.

Even at the NMJ, where postsynaptic nAChR responses have been studied kinetically, pharmacologically, and computationally for decades, it has been necessary to infer the diffusional spread of ACh within the synaptic cleft. Simulations suggest that [ACh] reaches several mM for < 1 ms and that when AChE is inhibited, ACh diffusion is kinetically buffered by rebinding to receptors^90^. Because the stopped-flow kinetics (**Supp. Fig. S8**) show that iAChSnFR activation kinetics approach 100 s^-1^ at such concentrations, one can hope to test these conclusions.

The sensor also enabled new observations about the microphysiology of transmission at muscarinic synapses. Previous data show that when G_i_-coupled endogenous mAChRs (*i.e.* M2 or M4) on neurons or cardiac myocytes are subjected to a step of [agonist] or a decrease of [antagonist] on a time scale of ∼1 ms, recorded GIRK channel currents show a period of zero slope, followed by a current that rises to a peak in a few hundred ms. This resembles responses after presynaptic stimulation, suggesting that coupling to G_i_ and activation of G_i_ protein effectors limits the time course of muscarinic transmission. iAChSnFR now enables us to compare the waveform of the postsynaptic response with that of pre-synaptically released ACh. Our data (*e.g.* **Supp. Fig. S20d**) show conclusively that ACh release is complete on a time scale of 10 ms or less; in fact, [ACh] has already begun to decay when the GIRK current reaches its peak.

The microphysiological knowledge empowered by iAChSnFR also has clinical implications. For example, acetylcholinesterase (AChE) inhibitor treatment, an important therapy for Alzheimer’s disease^91^, produces sub-optimal cognitive improvement and introduces medication termination-associated irreversible, accelerated deterioration^92^. The properties of cholinergic transmission resolvable by iAChSnFR can account for these clinical observations. For example, acetylcholinesterase inhibitors could cause less precise cholinergic transmission (cf.^93^), explaining the modest cognition improvement. Moreover, long-term application of AChE inhibitors could lead to up-regulation of AChE levels in Alzheimer’s patients and/or down-regulation presynaptic ACh release^91^, explaining the accelerated deterioration. The new insights into regulation and precision of cholinergic transmission not only suggests presynaptic cholinergic regulatory mechanisms to be effective Alzheimer’s intervention targets, but also sets the normal baseline for testing of candidate therapies. Likewise, analysis of cholinergic transmission at other cell types may shed new light on mechanisms and treatments of various other diseases, including diabetes^94^, immune deficiency^95^ and tumorigenesis^96, 97^.

The initial results at the *C. elegans* neuromuscular junction will allow detailed characterization of its release properties. Some *C. elegans* NMJs use GABA, for which there are good sensors^26^; together, these indicators will allow cataloguing the neurotransmitter usage and properties of all the components of the worm brain-muscle interface. Vertebrate NMJs use ACh exclusively; iAChSnFR will allow their study as well.

In flies, iAChSnFR showed quite large responses (up to 43% ΔF/F_0_ and a signal-to-noise ratio of 7) in olfactory glomeruli in response to odor addition. In similar experiments, GACh2.0 showed only a 0.4% ΔF/F_0_ response^20^, although the glomeruli imaged and odors added differed in the two experiments. iAChSnFR also demonstrated robust negative responses to some glomerulus/odor combinations, demonstrating sufficient dynamic range to detect decreases in tonic ACh release; the utility of the GACh sensors for such bidirectional imaging has not yet been demonstrated. The excellent ΔF/F_0_ and rapid kinetics will facilitate a large number of experiments in flies and other arthropods, which appear to use ACh as their predominant excitatory neurotransmitter.

In zebrafish, the finding that the greatest iAChSnFR signal on spinal motor neurons was observed at the proximal axon and the dendritic arbor makes predictions testable by forthcoming datasets such as the spinal wiring diagram of the larval zebrafish from serial-section electron microscopy^98^. More importantly, with the rapid growth of imaging modalities and expression lines for the fish, iAChSnFR enables direct visualization of cholinergic signaling in targeted cell types or whole-brain, which will complement results from other techniques.

In mouse, iAChSnFR enabled a number of interesting findings. In forelimb motor cortex, running greatly increased iAChSnFR responses, but signals at rest were large, indicative of substantial tonic cholinergic signaling. iAChSnFR signals at *Sst^+^* interneuron dendrites preceded those at pyramidal cell dendrites, consistent with observed biphasic ACh responses^65, 66^. On both cell types, iAChSnFR signals were highly spatially localized (sub-micron), consistent with our results in brain slice, and supportive of a model of ACh transmission that prominently features fast, local, phasic signaling, perhaps also bolstered by slow, broad, volume transmission. Results in visual cortex, observed iAChSnFR signals were also mostly fast and sub-micron; this preparation enabled us to study the depth-dependence of ACh transmission properties. We found that deep cortical layers showed more phasic transmission aspects, whereas shallow layers exhibited mostly slower responses. Future experiments will explore this phenomenon more deeply, and elucidate the contribution of differential expression of AChE and VAChT, specific isoforms of ChAT, synaptic release properties, and diverse nicotinic and muscarinic receptors to it.

iAChSnFR also revealed that both isoflurane and ketamine/xylazine anesthesia decreased cholinergic tone in the cortex. Further, we observed a depth-dependence in anesthetic effects, with deeper cortical layers being affected earlier and longer. The mechanism of general anesthesia largely remains a pharmacological mystery, although ACh is known (by microdialysis) to decrease in the cortex^75, 76^. The effects of anesthesia on ACh signaling can now be studied at greater spatial resolution and can be compared among anesthetics.

Finally, our preliminary data reveal that increases in hippocampal ACh are exquisitely time-locked with procurement of reward during operant conditioning. These findings suggest that ACh release in the hippocampus accompanies adaptive effortful responding and can be accessed to overcome economic constraints associated with escalating costs of a commodity. However, further work is required to better understand the precise causal role that sub-second hippocampal ACh release plays in orchestrating the pursuit of reward when its associated costs vary as a function of response requirement.

The spatial and temporal precision of iAChSnFR-measured ACh concentrations will enable neuroscience to test and to improve existing concepts such as the distinctions among “point-to-point transmission”, “volume transmission”, and “neuromodulation”. It will also become possible to measure the detailed spatial and temporal aspects of co-release. At neurons that co-release ACh and GABA, iAChSnFR and iGABASnFR signals can now be compared. Likewise, at neurons that co-release ACh and glutamate, iAChSnFR and iGluSnFR release can be compared. That synapses are exquisite biophysical machines, specialized to function on a time scale of milliseconds and a distance scale of micrometers, underlies the density and speed of information processing in nervous systems. The new biosensor, iAChSnFR, allows further descriptions, on the relevant time and distance scale that complement existing, primarily postsynaptic, electrophysiological measurements (**Figs. 2, 6, S14 and S20**).

Given its broad utility in diverse organisms (in the brain and at the NMJ), excellent fluorescence change, kinetics, and photostability, utility alongside most cholinergic drugs, and apparent absence of effects on cellular function and cholinergic signaling, iAChSnFR will be a valuable tool for the investigation of synaptic and circuit dynamics *in vivo*. It is likely that further improvements can be made to the iAChSnFR scaffold, both in improving the performance of the existing green and yellow versions, and in creation of high-performance red variants.

## Materials & methods

### Sensor engineering and characterization

The methods used to generate sensor constructs and linker variant libraries are essentially the same as described previously^24, 29^. Briefly, the gene encoding OpuBC from *Bacillus subtilis* and its hyperthermophilic homologue from *Thermoanaerobacter sp. X513* were ordered as gBlocks from IDT. These were then cloned into a modified pRSET-A vector (Invitrogen, renamed pHHM) using a BglII site encoding cysteine-serine at the 5’ end and a PstI site encoding leucine-glutamine prior to the C-terminal Myc tag. Circularly permuted Superfolder-GFP was included in the gBlock, inserted after amino acid 83 in OpuBC and 106 in the *X513* variant. Linker variants, binding site mutations, hinge mutations, and interface mutations were introduced by site-saturation mutagenesis and screened in a high-throughput manner as described previously^24, 29^. Briefly, mutations were introduced using uracil template-based mutagenesis^99^, colonies were picked into deep well 96-well plates, grown overnight in auto-induction media^100^, washed 3 times with phosphate buffered saline (PBS), and frozen. Cleared lysates were then titrated with acetylcholine and fluorescence changes observed. Fluorescence measurements were acquired using a Tecan Infinite M1000 Pro plate reader, using 485 nm excitation and 515 nm emission settings. Detailed amino acid sequences are provided in **Supplementary Fig. 3**. The non-binding version of the sensor (iAChSnFR-NULL) was made by abolishing acetylcholine binding through alanine mutation of one of the key aromatic residues involved in cation-pi interactions (Tyr357).

### Protein expression and purification

pHHM.iAChSnFR plasmids were transformed into *E. coli* BL21(DE3) cells (lacking pLysS). Protein expression was induced using liquid auto-induction medium supplemented with 100 μg/mL ampicillin at 30°C. Proteins were purified by immobilized by Ni-NTA affinity chromatography in 0.1 M PBS containing 137 mM NaCl, pH 7.4. Proteins were then eluted using a 120 mL gradient from 0 mM to 200 mM imidazole. Following elution, fractions containing the sensor were pooled, concentrated using Vivaspin concentrators (Sartorius), and dialyzed extensively in PBS.

### Biochemical characterization

Acetylcholine, neurotransmitters, acetylcholinesterase inhibitors, choline, and glycine-betaine were purchased from Sigma. 200 nM of purified sensor was used for all titrations. Ligand binding data were fit to a single-binding site isotherm. pH titrations were performed using Hydrion buffers from pH 2-11 (Sigma). Circular dichroism (CD) measurements were made using 1 μM purified sensor in 0.1X PBS using a Chirascan (Applied Photophysics) CD spectrophotometer. Stopped-flow analyses were performed using 200 nM sensor in PBS at room temperature on a SX20 LED stopped-flow instrument (Applied Photophysics) using a 490 nm excitation LED and 525 nm long-pass filter.

### Protein crystal structure determination

#### iAChSnFR0.4, ligand-free structure

iAChSnFR was expressed and purified as described above, with an additional purification step consisting of size exclusion chromatography. After Ni-NTA purification, the protein was loaded onto a Superdex 200 26/600 HiLoad column (GE Healthcare) with 5 mM MOPS, 50 mM NaCl, 1 mM EDTA, pH 7.6 as the running buffer, and the protein was eluted at a flow rate of 2 mL/min. Proteins were concentrated to 10-30 mg/mL for crystallization as described above. Crystallization was carried out at room temperature by hanging-drop vapor diffusion using commercially available sparse-matrix screens (Index, Hampton Research; JCSG Plus, Joint Center for Structural Genomics) by mixing varying ratios (1:3, 1:1, 3:1) of protein and precipitant solutions. From initial screens 0.05 M MgCl_2_, 0.1 M HEPES pH 7.5, 30% v/v polyethylene glycol monomethyl ether 550 (Hampton Index Reagent 55) was identified as the precipitant yielding the best quality crystals. Crystals were collected and shipped for data collection at the Advanced Light Source, Berkeley, CA on beamline 8.2.1. X-ray data were reduced using Mosfilm^101^. Structures were solved by molecular replacement using P^102^ and the CCP4 package^103^. Iterative cycles of model building in Coot^104^ and Refmac/CCP4/Phenix^105^ produced the models. Final protein structures have been deposited in the Protein Data bank (http://www.rcsb.org).

#### iAChSnFR, periplasmic binding protein-only

Deletions of the N-terminal HA-tag, C-terminal Myc tag of the biosensor and GFP were carried out with Q5 Site-Directed Mutagenesis Kit (New England Biolabs). The plasmid encoding this engineered periplasmic acetylcholine-binding protein (sequence MHHHHHHGGAQPARSANDTVVVGSIIFTEGIIVANMVAEMIEAHTDLKVVRKLNLGGVNVNFEAIKRGGANNGIDIYVEYTGHGLVDILGFPATTDPEGAYETVKKEYKRKWNIVWLKPLGFNNTYTLTVKDELAKQYNLKTFSDLAKISDKLILGATMFFLEGPDGYPGLQKLYNFKFKHTKSMDMGIRYTAIDNNEVQVIDAWATDGLLVSHKLKILEDDKAFFPPYYAAPIIRQDVLDKHPELKDVLNKLANQISLEEMQKLNYKVDGEGQDPAKVAKEFLKEKGLILQVD) was transformed into *E. coli* BL21-gold (DE3) cells (Agilent Technologies). Protein overexpression was carried out with ZYM-5052 autoinduction media at room temperature (Studier 2005). For purification, cells were lysed with four cycles of freeze-thaw in lysis buffer containing 100 mM NaCl, 20 mM Tris, pH 7.5, lysozyme, DNase and protease inhibitors. After centrifugation at 14,000 rpm for 30 min at 4°C to remove cell debris, the supernatant was collected and loaded onto a pre-washed NiNTA column at 4°C. NiNTA wash buffer contains 100 mM NaCl, 20 mM Tris, 30 mM imidazole, pH 7.5, and elution was carried out with the same buffer containing 300 mM imidazole, pH 7.5. The eluted sample was subjected to size-exclusion chromatography with buffer containing 100 mM NaCl and 20 mM Tris, pH 7.5 using a HiLoad 16/60 Superdex 200 column (GE Healthcare). Peak fractions were pooled and concentrated to ∼50 mg/ml with Amicon Ultra 15 concentrator (Millipore) with 10 kDa cutoff.

Crystallization trials were carried out at 40 mg/ml in the presence and absence of 10 mM acetylcholine using different commercial screens. The protein crystallized in Wizard 1 (Molecular Dimensions), condition #1, with 20% PEG 8,000, 0.1 M CHES, pH 9.5. Conditions was further optimized with the Additive Screen (Hampton Research), with the final mix containing an additional 10 mM spermidine.

X-ray datasets were collected at Stanford Synchrotron Radiation Lightsource (SSRL) beamline 12-2 with a Pilatus 6M detector using a Blu-Ice interface^106^ The data was integrated with XDS^107^, and scaled with aimless^103^. Molecular replacement was carried out using Phaser in Phenix^105^ with sub-domains of iNicSnFR (PDB ID: 6EFR)^36^ as the input model. Subsequent refinement and model building cycles were carried out with phenix.refine^105^ and Coot^104^, respectively.

### Membrane anchored iAChSnFR on HEK293 cells

For mammalian expression, iAChSnFR variants were cloned into a modified pDisplay vector (Invitrogen, renamed pMinDis) lacking the hemagglutinin-A (HA) tag and only retaining the BglII site in the multiple cloning site. pMinDis.iAChSnFR variants were transiently transfected into HEK293 cells (ATCC) using Amaxa electroporation and grown in high-glucose DMEM supplemented with 10% FBS and 2 mM glutamine in 35-mm MatTek dishes. After 1 d incubation at 37°C, cells were washed extensively with Hanks Balanced Salt Solution (Invitrogen) and imaged on a Zeiss LSM 510 inverted confocal microscope using a Zeiss 20x/0.8 NA Plan-Apochromat objective, controlled with Zeiss Zen software. Images were acquired continuously at a rate of one frame per second at a resolution of 512 x 512 pixels. For acetylcholine titration experiments, images were collected for 1 minute at each concentration. The fluorescence level at each concentration was averaged over the entire minute for several regions of interest (ROIs) per experiment. Binding curves were fit as above.

### Neuronal culture characterization

Mixed cultures of neurons and glia were prepared from Sprague-Dawley rat pups (Charles River) and plated on laminin-coated MatTek dishes. Three days after plating, cultures were infected with 10^12–14^ GC/mL of AAV2/1 viral particles, carrying iAChSnFR driven by the human *Synapsin-1* promoter. One week after infection, cultures were imaged and titrated with acetylcholine as above.

### Cultured slice preparation

Cultured slices were prepared from P6−7 rats following the previous studies^47, 108^. In brief, hippocampi were dissected out in ice-cold HEPES-buffered Hanks’ solution (pH 7.35) under sterile conditions, sectioned into 400 µm slices on a tissue chopper, and explanted onto a Millicell-CM membrane (0.4-µm pore size; Millipore, MA). The membranes were then placed in 750 µL of MEM culture medium, contained (in mM): HEPES 30, heat-inactivated horse serum 20%, glutamine 1.4, D-glucose 16.25, NaHCO_3_ 5, CaCl_2_ 1, MgSO_4_ 2, insulin 1 mg/mL, ascorbic acid 0.012% at pH 7.28 and osmolarity 320. Cultured slices were maintained at 35°C, in a humidified incubator (ambient air enriched with 5% CO_2_).

### Acute brain slice preparation

Acute entorhinal cortical brain slices were prepared from P25-60 animals deeply anesthetized by xylazine-ketamine as described in our previous reports^20, 108^. The animals were decapitated and the brain block containing the entorhinal cortex or striatum was quickly removed and placed into cold (0−4°C) oxygenated physiological solution containing (in mM): 125 NaCl, 2.5 KCl, 1.25 NaH_2_PO_4_, 25 NaHCO_3_, 1 MgCl_2_, 25 D-glucose, and 2 CaCl_2_, pH 7.4. The brain blocks were sectioned into 400-μm-thick brain slices using a DSK microslicer (Ted Pella Inc.). The tissue slices were kept at 37.0 ± 0.5 °C in oxygenated physiological solution for ∼0.5−1 hour before imaging. During the recording and/or imaging the slices were submerged in a chamber and stabilized with a fine nylon net attached to a platinum ring. The recording chamber was perfused with oxygenated physiological solution. The half-time for the bath solution exchange was ∼6 s, and the temperature of the bath solution was maintained at 34.0 ± 0.5 °C. All antagonists were bath applied. For striatal experiments, the bath solutions contained DNQX (10 μM), picrotoxin (100 μM), CGP55845 (300 nM), SCH 23390 hydrochloride (1 μM) and sulpiride (200 nM) to block AMPA, GABA_A_, GABA_B_, dopamine D1 and dopamine D2-receptors. The bath solution lacking Ca^2+^ (Ca^2+^-free) was supplemented with 2 mM MgCl_2._ Distinct cell types, including L2 stellate neurons and astrocytes could be easily identified under transmitted light illumination based on their locations and morphology as characterized in the previous reports^20, 109^.

### Sindbis and lentivirus preparation and expression

iAChSnFR, iAChSnFR-NULL and its variants were sub-cloned into Sindbis and/or lentiviral constructs and viral particles were produced following our previous studies^47, 110^. In brief, iAChSnFR and its variants were sub-cloned into Sindbis viral vector pSinREP5 with the cloning sites Xba1 and Sph1, lentiviral vector pLenti-with the cloning sites BamH1 and Xho1/Not1 under a human synapsin promoter to ensure neuronal expression or under a GFAP promoter to ensure astrocyte expression. One astrocyte-targeting lentiviral construct performed best for *in vivo* expression^111^, and it was used in the experiments reported in **Figure 2**.

Expression of iAChSnFR, iAChSnFR-NULL or other variants was made as previously reported^47, 110^. Neurons in hippocampal cultured slices were infected after 8−18 days *in vitro* with Sindbis or lentivirus, and then incubated on culture media and 5% CO_2_ before experiments. For *in vivo* expression, P25−60 animals were initially anesthetized by an intraperitoneal injection of ketamine and xylazine (10 and 2 mg/kg, respectively) or isoflurane (1-3%), and then placed in a stereotaxic frame. A glass pipette was used to penetrate into the entorhinal cortex or striatum according to stereotaxic coordinates to deliver ∼50-500 nL of AAV, Sindbis or lentiviral solution by pressure injection to infect neurons or astrocytes with iAChSnFR. Experiments were typically performed within 18 ± 4 h after Sindbis viral infection or 1−2 weeks after lentiviral infection or 3-4 weeks after infection of AAV2/9-*hSynapsin1*.tdTomato.T2A.mGIRK2-1-A22A.WPRE.bGH.

### Electrophysiology

Simultaneous dual whole-cell recordings were obtained from two nearby infected and non-infected hippocampal CA3 pyramidal neurons under visual guidance using fluorescence and transmitted light illumination^47, 108^. The patch recording pipettes (4−7 MΩ) were filled with intracellular solution 120 mM potassium gluconate, 4 mM KCl, 10 mM HEPES, 4 mM MgATP, 0.3 mM Na_3_GTP, 10 mM sodium phosphocreatine and 0.5% biocytin (pH 7.25) for voltage-clamp recordings. Bath solution (29 ± 1.5°C) contained (in mM): NaCl 119, KCl 2.5, CaCl_2_ 4, MgCl_2_ 4, NaHCO_3_ 26, NaH_2_PO_4_ 1 and glucose 11, at pH 7.4 and gassed with 5% CO_2_/95% O_2_. Whole-cell recordings were made with up to two Axopatch-200B patch clamp amplifiers (Molecular Devices, Sunnyvale, CA).

### Neuronal reconstruction

To recover the morphology of recorded neurons, the slices were fixed by immersion in 3% acrolein/4% paraformaldehyde in 0.1 M PBS at 4°C for 24 hours after *in vitro* patch-clamp recordings with internal solution containing additional 1% biocytin, and then processed with the avidin-biotin-peroxidase method to reveal cell morphology as previously reported^20, 110^. The morphologically recovered cells were examined and reconstructed with the aid of a microscope equipped with a computerized reconstruction system Neurolucida (MicroBrightField, Colchester, VT).

### Fluorescence imaging of cells in cultured and acute slice preparations

Wide-field epifluorescence imaging was performed using Hamamatsu ORCA FLASH4.0 camera (Hamamatsu Photonics, Japan), and iAChSnFR- or iAChSnFR-NULL-expressing cells in cultured hippocampal slices and acutely prepared brain slices are excited by a 460-nm ultrahigh-power low-noise LED (Prizmatix, Givat-Shmuel, Israel)^47, 108^. The frame rate of FLASH4.0 camera was set to 10-50 Hz. To synchronize image capture with drug perfusion, electrical stimulation, and/or electrophysiological recording, the camera was set to external trigger mode and triggered by a custom-written IGOR Pro 6 program (WaveMetrics, Lake Oswego, OR)-based software PEPOI^108^. Agonists including acetylcholine, and nicotine, oxotremorine M (Tocris Bioscience, Bristol, UK), were puff applied with a glass pipette (∼1 µm in tip diameter) positioned ∼150 µm above the imaged neurons using 500-ms 30-kPa pressure pulses, and acetylcholinesterase inhibitor donepezil was bath applied. Cholinergic fibers in tissue slices were stimulated with a bipolar electrode placed ∼50−200 µm from imaged cells with single or a train of voltage pulses (500 µs, up to 50 V) to evoke ACh release.

Two-photon imaging was performed using a custom-built microscope operated by PEPOI^108^. The parameters of frame scan were typically set at a size of 200 × 200 pixels and a speed of 1 frame/s. The fluorescence of iAChSnFR was excited by a femtosecond Ti:Sapphire laser (Chameleon Ultra II, Coherent) at a wavelength of 950 nm. To quantify surface expression of GACh sensors, lentiviral expression of iAChSnFR was made in the CA1 region of organotypic hippocampal cultured slices. About ∼1−2 weeks after expression, iAChSnFR expressing CA1 pyramidal neurons were patch-clamp recorded and loaded with 25 µM Alexa Fluor 594 (Life Technologies) for ∼10 minutes, and two-photon images were then taken at different compartments along the somatodendritic axis. ACh-induced iAChSnFR fluorescent changes at the striatum were measured using 2-photon spot photometry (Toronado, B.W. Strowbridge), by scanning the laser around a circular path (157 nm diameter, 2 KHz) centered on a fluorescent punctum of interest. The green fluorescence was collected by a water immersion objective (60x, N.A. 1.00; Olympus), a dichroic mirror (T700LPXXR, Chroma) and filtered (ET680sp and ET525/50m-2P, Chroma).

### Nanoscopic imaging analysis

Evoked ΔF/F_0_ responses of iAChSnFR expressing cells were typically imaged for periods from seconds to minutes. To minimize drifting and fluctuating movements, a stable recording/stimulating and imaging setup was used to carry out all imaging experiments^108^. Image analysis was performed using our custom-written MATLAB-based image processing and analysis program (MIPAP), with algorithms created using MATLAB 2018a with MATLAB’s Image Processing Toolbox (Mathworks). After correcting photobleaching and auto fluorescence, all images were spatially aligned to the initial reference images using an intensity-based registration function to correct the minimal image drifts and fluctuations. Then, the fluorescence ΔF/F_0_ responses at individual pixels were quantified, rescaled and converted into pseudo-color scales to create heatmap images. The maximal electrically evoked maximal ΔF/F_0_ responses at individual pixels over time were subsequently plotted to create 3D spatial profiles for individual transmitter release sites. To reduce other sources of noise (e.g., pixelation noise and background noise), super-resolution localization microscopy analysis strategies were employed to average over multiple exposures, multiple releases and/or multiple directions of transmitter diffusion^112–114^. With these, the evoked ACh release displayed many consistent and clear nanoscopic release hotspots, and pixels with the maximal ΔF/F_0_ responses were assumed to be the centers of release. The average fluorescence ΔF/F_0_ intensity profiles normalized to the peak values fit well with a single exponential decay function, and the decay constants were extracted as spatial spread length constants. To compensate microscopic point-spread function diffraction, we obtained our microscopic point-spread function with 23 nm green GATTA beads (GATTAquant GmbH, Hiltpoltstein, Germany). Deconvolution based on this measured point-spread function yielded the true spread length constants to be 0.76 µm and 0.73 µm at entorhinal stellate neurons and astrocytes, respectively, indicating ∼35% overestimation.

### Slice experiments (Bonn)

All procedures were planned and performed in accordance with the guidelines of the University of Bonn Medical Centre Animal-Care-Committee as well as the guidelines approved by the European Directive (2010/63/EU) on the protection of animals used for experimental purposes. All efforts were made to minimize pain and suffering and to reduce the number of animals used, according to the ARRIVE guidelines. Mice were housed under a 12h light–dark-cycle (light-cycle 7 am/7pm), in a temperature (22±2°C) and humidity (55±10%) controlled environment with food/water *ad libitum* and nesting material (nestlets, Ancare, USA). Animals were allowed at least 1 week of acclimatization to the animal facility before surgery and singly housed after surgery. Juvenile C57/BL6J mice (5-7 weeks old, Charles River, Sulzfeld, Germany, male) were bilaterally injected with AAV2/1. *hSynapsin-1*.iAChSnFR following i.p. anesthesia with ketamine (100 mg/kg) and xylazine (16 mg/kg). Stereotaxic coordinates were 2.5, 3.0, and 3.0 mm (posterior, lateral, dorso-ventral) relative to Bregma. Coordinates were selected to target the CA3/CA1 region of the ventral hippocampus. Mice were treated with analgesics (0.05 mg/kg buprenorphine, s.c. and 5 mg/kg ketoprofen, s.c.) and antibiotics (5 mg/kg Baytril, s.c.) 30 minutes before the operation started. Following anesthesia, animals were positioned in the stereotaxic frame, the skin and fascia were retracted to reveal the skull, and the positions were measured. Small holes were drilled just through the bone, and the injection needle was slowly lowered to the ventral position. 1 µL of virus (undiluted, titer: 8-10 × 10^12^) was injected into each hemisphere over the course of 5 minutes. Following injection, the needle was left in place for 2 minutes, and then withdrawn. The scalp was sutured, and mice were left to recover on a heating pad until awakening from anesthesia. Mice were treated with analgesics (twice per day with 0.05 mg/kg buprenorphine, s.c. and once per day with 5 mg/kg ketoprofen, s.c.) and monitored for 3 days following surgery and then utilized in the imaging experiments 2-3 weeks following virus injection. Brain slices with strong expression in the stratum radiatum of CA1 were selected for the experiment. To quantify iAChSnFR signals, a confocal scan (488nm, Zeiss Pascal) was acquired in the stratum radiatum (emission band pass 514/40 nm). The scans were positioned next to the stimulation pipette (glass ∼1-2 MOhm resistance). During stimulation, a series of 30-50 frames of 512 x 128 pixels (0.41 µm pixel size) was acquired at 120 - 300 millisecond intervals and repeated every minute. Background-corrected ΔF/F per ROI was calculated to assess signal strength and quantify the effect of pharmacological manipulations.

### C. elegans methods

We used *Caenorhabditis elegans* strain JSD1115 (*tauEx461*), derived from strain N2 by injection with JP992 [Pmyo-3::ss(*unc-52*)-iAChSnFR0.7] at 20 ng/µL and *Pacr-2*::mCherry::RAB-3 at 10 ng/µL. For worm expression, the human signal sequence was replaced with the signal sequence from UNC-52 (perlecan, a muscle heparan sulfate proteoglycan). iAChSnFR0.7 was expressed in body-wall and egg-laying muscles using the *myo-3* promoter.

*C. elegans* were maintained on agar nematode growth media (NGM) at 20°C and seeded with OP50 bacteria as previously described^115^. We studied young-adult (4 days after hatching) hermaphrodite worms.

To image iAChSnFR fluorescence, worms were immobilized in a mixture of polystyrene beads and 8% agarose on a coverslip^116^. Images were acquired on an inverted Olympus microscope (IX81), using a laser scanning confocal imaging system (Olympus Fluoview FV1000 with dual confocal scan heads) and an Olympus PlanApo 60X 1.42 NA objective.

### Methods for iAChSnFR imaging in astrocytes in acute brain slice

AAV injection procedures have been described previously^117^. About 1 µL of AAV2/5.*GfaABC_1_D*.iAChSnFR (∼1 x 10^10^ genome copies) was injected into left dorsal striatum or CA1 region of the hippocampus of C57BL/6NTac mice (Taconic, 8-11 weeks old, male and female). Imaging experiments were performed 18-19 days after the injection. Slice procedures have been described previously^117^. Coronal (hippocampus) or para-sagittal striatal slices were prepared. Briefly, animals were deeply anesthetized with isoflurane and decapitated. The brain slices (300 µm) were cut in ice-cold modified artificial cerebrospinal fluid (aCSF) containing the following (in mM): 194 sucrose, 30 NaCl, 4.5 KCl, 1 MgCl_2_, 26 NaHCO_3_, 1.2 NaH_2_O_4_, and 10 D-glucose, saturated with 95% O_2_ and 5% O_2_. The slices were allowed to equilibrate for 30 min at ∼32 °C in normal aCSF containing (in mM): 124 NaCl, 4.5 KCl, 2 CaCl_2_, 1 MgCl_2_, 26 NaHCO_3_, 1.2 NaH_2_PO_4_, and 10 D-glucose saturated with 95% O_2_ and 5% O_2_. Slices were subsequently stored at room temperature. Slices were imaged using a confocal microscope (Fluoview 1000; Olympus) with 40X water-immersion objective lens with a numerical aperture (NA) of 0.8 and at a digital zoom of 1.5. The 488 nm line of an Argon laser was used to excite iAChSnFR. The emitted light pathway consisted of an emission band pass filter (505-605 nm) before a photomultiplier tube. A stimulus electrode (TST33A05KT, WPI) was placed at the corpus callosum. Imaging was performed at 10-210 µm away from the electrode. Analysis was performed at 100-200 µm away from the electrode. Four or eight trains (10 Hz, 0.6 mA) of electrical field stimulation (EFS) was applied. Images were analyzed by FIJI^118^. Time traces of fluorescence intensity were extracted from the ROIs and converted to ΔF/F_0_ values.

### Drosophila preparation, odor delivery, two-photon imaging and data analysis

Ten-to-fifteen day-old adult female *Drosophila melanogaster* (w^-^; GH146/+; 20xUAS-BiP-iAChSnFR-SV40 [VK00005]/+) were mounted on the bottom of a custom-made holder^119^ using UV-activated epoxy (Loctite 3972), with the top of the head capsule accessible from above and the antennae exposed to air. An adult hemolymph-like buffer^120^ was loaded into the holder, then the cuticle and trachea above the brain were carefully dissected to allow optic access to the antennal lobe.

Pure odorants were freshly diluted with paraffin oil (Sigma-Aldrich, no. 18512) and stored in glass vials (BD Vacutainer, no. 366430). The odor delivery system was driven from pressure-regulated, filtered, compressed air and the flow rate through all lines were controlled by Mass Flow Controllers (Alicat Scientific). A carrier air stream of 450 standard cubic centimeters per minute (sccm) was directed at the antennae of the fly through the end of a syringe needle. The odor delivery system consists of two stages of manifolds with ultrafast miniature semi-inert three-way solenoid valves (The Lee Company, no. LHDA1231515H) and integrated hit-and-hold control circuits (Bio-Chem Fluidics Inc., no. CoolCube-50R). The first stage selects the odor to fill the odor line, 6 seconds before the actual stimulation; the second stage injects either the control air stream (50 sccm) or the odor stream (50 sccm) into the carrier air stream. Before stimulation, the second stage directs the control air into the carrier and the odor stream away; upon stimulation, the second stage directs the odor stream into the carrier and simultaneously directs the control air away; after 2 seconds, the second stage swaps the streams back to their pre-stimulation directions. After another 6 seconds, all lines are flushed with the control air stream for 16 seconds. The air flows through the odor delivery system continuously throughout the experiment and pressure never builds up in any odor vials. For each fly, a custom-written MATLAB (MathWorks) program controlled the structure and timing of the experiment through a data acquisition card (National Instruments, no. PCIe-6323). Six blocks of odor trials were presented to each fly. Each block includes 8 individual trials, each with either the control (paraffin oil) or any of the 7 odor/concentration combinations, in a randomized sequence. The odor/concentration pairs used are 10^-3^/10^-2^/10^-1^ 3-octanol (OCT), 10^-3^/10^-2^/10^-1^ isoamyl acetate (IAA), and 10^-3^ benzaldehyde (BEN). All pure odorants are highest grade from Sigma-Aldrich.

Two-photon imaging was conducted on a modified Prairie Ultima Two-Photon Microscope (Bruker). The iAChSnFR expressed in the antennal lobes of the flies was excited with a Coherent Chameleon Vision II Ti:Sapphire femtosecond laser tuned to 920 nm and guided with one resonant galvanometer (8 kHz) along the x-axis and one non-resonant galvanometer along the y-axis. The emitted photons were collected with a GaAsP Photomultiplier Tube (Hamamatsu, no. 7422PA-40). Each imaging cycle was triggered by a TTL signal from the same NI-DAQ card that controlled odor delivery. The water-immersion objective (Olympus, LUMPlanFl/IR, 40x, 0.80 NA) was driven by a piezo so that a z-series of 4 different focal planes (17 μm between planes) was taken at each time point and repeated continuously until the end of the trial. Each plane was imaged with 2x optical zoom at 256 x 256 pixels within 16 milliseconds. The interval between neighboring frames from the same focal plane was 150 milliseconds, resulting in a rate of 6.7 volumes/s.

Image analysis was conducted after acquisition with custom ImageJ (NIH) and MATLAB routines. Briefly, image frames were assigned to their corresponding focal planes and registered with the TurboReg plug-in^121^. The regions corresponding to the background and various antennal lobe glomeruli were defined manually in ImageJ. The average pixel intensity within the background region and the regions of interest (ROIs) were computed for each frame. Olfactory response vectors were computed for each ROI after subtracting the background average intensity from the ROI average intensity for each time point. A Savitzky-Golay smoothing filter, with the polynomial order of 1 and a frame length of 3, was applied to each response vector for data shown in Figure 5, Panels B and C, and a frame length of 7 for Figure 5 Panel A. For each fly, the response vectors for the same odor-glomerulus pair over 6 repetitions were averaged as the mean response vector, then the normalized mean response vectors (Δ*F/F_0_* = (*F − F_0_*)/*F_0_*) were computed with F_0_ being the mean of the mean response vectors during the 2 seconds before stimulation. The grand mean of the normalized mean response vectors and the standard errors of the mean (N = 4 flies) were plotted in Figure 5, Panels Bi and Ci. The maximum (for excitation) and minimum (for inhibition) ΔF/F_0_ for all odor-glomerulus pairs during the 4 seconds after stimulus onset are presented in Figure 5, Panels Bii and Cii.

### Zebrafish transgenic line and imaging

All experiments were conducted in accordance with protocols approved by the Institutional Animal Care and Use and Institutional Biosafety Committees of the Howard Hughes Medical Institute, Janelia Research Campus. An immotile mutant, *relaxed*, was used to avoid need of the cholinergic blocker α-bungarotoxin for immobilization^122^. *Relaxed* fish have a homozygous loss-of-function mutation of the dihydropyridine receptor b1 gene, *Cacnb1*, which couples action potential and contraction of muscle fibers through voltage-dependent Ca^2+^ release^122, 123^. For expression in zebrafish, we replaced the mammalian IgG secretion sequence in iAChSnFR with the signal peptide from zebrafish Nodal-related 1 (Ndr1, squint). iAChSnFR was transiently expressed in spinal motor neurons by injecting plasmid containing 10xUAS:Ndr1peptide-iAChSnFR into fertilized eggs from in-crosses of heterozygous *relaxed*, TgBAC(vacht:Gal4) fish. The embryos were manually removed from the chorion with a pair of forceps at 2 days post-fertilization. By 3 days post-fertilization, immotile embryos were screened for the absence of tactile-induced escape behavior and for iAChSnFR fluorescence under an epifluorescence microscope. At 3- or 4-days post-fertilization, a screened embryo was embedded with the lateral side-up in 1.6% low-melting-point agar (MilliporeSigma, 2070-OP, MO). The agar covering the head and mid-body (from 14^th^ to 20^th^ segments) were removed. A brief bipolar electrical stimulus (1 ms in duration, 5-10 V) was delivered using a stimulus isolator (BSI-950, Dagan) through a concentric tungsten electrode placed on the bottom side of the head. Motor activity was recorded from the mid-body muscles through a glass pipette by an extracellular amplifier (EXT-2B, NPI). The inter-stimulus interval was longer than 1 min. The stimulus amplitude was adjusted to evoke strong motor activity consistently. An iAChSnFR-expressing spinal motor neuron was imaged with a custom two-photon microscope equipped with a resonant scanner (CRS 8 KHz, Cambridge Technologies) using ScanImage software (Vidrio Technologies, VA). Femtosecond laser pulses of 940 nm, 80 MHz (MKS Instruments, Mai Tai HP, Deep See, MA) were used to excite iAChSnFR. For imaging soma and neural processes at frame rate of 50 Hz (512 x 300 pixels), a Bessel beam was used to axially extend foci of excitation beam through an axicon lens module^124^. iAChSnFR fluorescence was detected by a photomultiplier tube (H11706-40, Hamamatsu) through a 25x objective lens (Leica, HCX IRAPO L 25x/ 0.95 W, Germany), a dichroic mirror (Semrock), and emission filter (Semrock). The images were analyzed in MATLAB R2016b (MathWorks). Fluorescence time course of each ROI was extracted and changes relative to baseline fluorescence was calculated.

### Mouse husbandry

All experiments were conducted in accordance with protocols approved by the Institutional Animal Care and Use and Institutional Biosafety Committees of the Howard Hughes Medical Institute, Janelia Research Campus, New York University, University of Pennsylvania, University of Colorado School of Medicine, University Clinic Bonn, University of California, Los Angeles, University of Maryland School of Medicine, or University of Virginia School of Medicine. AAV experiments were conducted with wild-type C57BL/6J (Charles River), *Rbp4*-Cre (MMRRC; stock no. 031125-UCD), *Chat*-Cre (Jackson Laboratory; stock no. 006410), and *Sst*-IRES-Cre (Jackson Laboratory; stock no. 013044), unless stated otherwise. Mice were group-housed in temperature-controlled rooms on a 12-h light-dark cycle and were randomly assigned to different treatment groups. 1-to-2-month-old animals of both sexes were used for all the experiments.

### Surgical preparation for imaging awake, head-restrained mice in motor cortex

Dendritic and somatic imaging was carried out in awake, head-restrained mice in a quiet resting state and forward running. Surgery preparation for awake animal imaging included attaching a head holder and creating a transcranial window. Specifically, mice were deeply anesthetized with an intraperitoneal injection of ketamine (100 mg/kg) and xylazine (15 mg/kg) or isoflurane (5% for induction, 1% for maintenance). The mouse head was shaved, and the skull surface was exposed. The periosteal tissue over the skull surface was removed and a head holder composed of two parallel micro-metal bars was attached to the animal’s skull with glue (Loctite 495) to help restrain the animal’s head and reduce motion-induced artifacts during imaging. A small skull region (∼0.2 mm in diameter) was located over the primary motor cortex based on stereotactic coordinates (at Bregma and 1.2 mm lateral from the midline) and marked with a pencil. This region was then thinned with a dental drill to create a thinned skull window, or a piece of skull was removed and a round glass coverslip (approximately the same size as the bone being removed) was glued to the skull. Dental acrylic cement (Lang Dental Manufacturing Co.) was applied to the surrounding area to secure the metal bars, taking care not to cover the thinned region or glass coverslip with cement. Upon awakening, mice with head mounts were habituated three times (10 min each time) in a custom-built imaging platform to minimize potential stress effects of head restraint and imaging. In this device, the animal’s head-mount was secured to metal blocks such that the head was fixed and perpendicular to the two-photon objective. The animal’s body was allowed to rest against the bottom of the solid imaging plate. Imaging experiments were started ∼12–24 h after window implantation and free from anesthetic effects.

### Treadmill running under the two-photon microscope

A free-floating treadmill was used for motor skill training in this study. This free-floating treadmill allows head-fixed mice to move their forelimbs freely to perform running tasks. Animals were positioned in a custom-made head holder device. The device enables forelimbs to rest on the belt and the rest of body to be positioned off-belt on a perch overlying the treadmill belt. During motor training, the treadmill motor (Dayton, no. 2L010) was driven by a DC power supply (Extech). At the onset of a trial, the motor was turned on and the belt speed transitioned from 0 cm s^−1^ to 8 cm s^−1^ within ∼1 s. Each running trial lasted 10 s or 30 s. After completion of each trial, the treadmill was turned off and the next trial started after a 30 s rest period.

### Sensor expression and imaging in primary motor cortex

The genetically encoded activity indicators used in this study include GCaMP6-sensitive (GCaMP6s) for Ca^2+^ imaging and iAChSnFR0.6, iAChSnFR0.6-NULL, and iAChSnFR for acetylcholine transient imaging. Sensors were expressed in L2/3 or L5 pyramidal neurons and *Sst*^+^ interneurons in M1. For L2/3 PCs, AAV2/1.*hSynapsin1*.GCaMP6s was used (>2 × 10^13^ GC/mL titer; produced by the University of Pennsylvania Gene Therapy Program Vector Core). iAChSnFR variants were expressed as AAV2/1.*hSynapsin1* virus (>2 × 10^13^ GC/mL titer; produced by the Janelia virus core). For *Sst*^+^ Cre mice, GCaMP6s was expressed with AAV under a Cre-dependent CAG promoter (AAV2/1.*CAG*.FLEX-GCaMP6s; >2 × 10^13^ GC/mL titer; produced by the University of Pennsylvania Gene Therapy Program Vector Core). iAChSnFR variant expression in *Sst*^+^ cells and L5 PCs was accomplished with AAV2/1.*CAG*.FLEX virus (>2 × 10^13^ GC/mL titer; produced by the Janelia virus core). For all AAVs, 0.1 μL of virus was injected (Picospritzer III; 15 p.s.i., 12 ms, 0.8 Hz) into L2/3 or L5 of M1 using a glass microelectrode over 10–15 min. Mice were prepared for head fixation and imaging after 2 weeks of AAV expression.

In **Supp. Fig. S21**, AAV2/1.*CAG*.FLEX-GCaMP6s was injected into NB based on stereotaxic coordinates (posterior 1 mm from Bregma and lateral 2.25 mm, at a depth of 4 mm). To evaluate the expression patterns of GCaMP6s axonal projections originating from NB, mice were fixed with 4% paraformaldehyde and the brain was sectioned at 200 μm thickness and immunostained against GFP (Abcam, no. ab13970, 1:300). Confocal microscopy (Zeiss LSM 700 confocal microscope; 10× air objective, NA 0.3) revealed numerous GCaMP6^+^/GFP^+^ axonal projections in M1.

*In vivo* two-photon imaging was performed with an Olympus Fluoview 1000 2-photon system equipped with a Ti:Sapphire laser (MaiTai DeepSee, Spectra Physics) tuned to 920 nm. The average laser power on the sample was ∼10-15 mW for L1 imaging and 15-25 mW in L2/3 of the cortex. All experiments were performed using a 25× objective immersed in an ACSF solution and with 2× (somata) or 4-6× (dendrites) digital zoom. All images were acquired at frame rates of 2–14 Hz (2 μs pixel dwell time). Image acquisition was performed using FV10-ASW v.2.0 software and analyzed post hoc using NIH ImageJ software.

### Direct NB stimulation

In **Supp. Fig. S22**, mice expressing GCaMP6s in either PCs or *Sst*^+^ cells were briefly anesthetized with isoflurane (1%) and secured in a stereotactic frame. A small area (200 μm) of skull, located 1 mm posterior from Bregma and 2.25 mm lateral, was drilled such that a bone flap was created. The bone flap was lifted, and a parallel bipolar electrode was advanced to a depth of 4.25 mm, at the position of the NB. Electrical stimulation was delivered at a frequency of 100 Hz, amplitude of 500 μA, in 0.2 ms pulses for 0.5 sec. In **Supp. Fig. S24**, mice expressing iAChSnFR0.6 were also stimulated with similar parameters except varying amplitudes (200 μA – 800 μA).

### Foot shock

Foot shocks (1 s, direct current, 0.9 mA) were delivered to the left hindpaw during recording trials.

### Motor cortex data analysis

During quiet resting, motion-related artifacts (related to the animal’s respiration and heart beat) were typically less than 2 μm as detected in our cortical measurements. Vertical movements were infrequent and minimized by habituation, the use of two micro-metal bars attached to the animal’s skull by dental acrylic, and the use of a custom-built body platform. If the animal struggled in the body platform, imaging time points from those segments were excluded from quantification. All imaging stacks were registered using NIH ImageJ plug-in StackReg.

Only cells expressing iAChSnFR or cytosolic GCaMP6s were included for quantification. Cells with apparent nuclear GCaMP6s expression^71^, although rare – typically at the center of the AAV injection site – were excluded from analysis. ROIs corresponding to visually identifiable somata (pyramidal cells and interneurons) were selected for quantification at different cortical depths from the pial surface. Imaging planes were from L2/3 and L5, corresponding to cells 200–300 μm below the pial surface and cells >500 μm below the pial surface, respectively.

The fluorescence time course in each cell body was measured with NIH ImageJ by averaging all pixels within the ROI covering the soma. The ΔF/F_0_ (as a percent change) was calculated as Δ*F/F_0_* = (*F − F_0_*)/*F_0_* × 100, where F_0_ is the baseline fluorescence signal averaged over a 0.5 s period corresponding to the lowest fluorescence signal over the recording period. In Ca^2+^ imaging experiments, we performed an integrated measurement of a cell’s output activity over 2.5 min recording, termed total integrated calcium activity, because it is difficult to report firing rates on the basis of calcium responses within individual cells.

At the depth of ∼10-100 μm below the pial surface, dendrites with iAChSnFR or GCaMP6s expression were included for analysis. For dendritic fluorescent changes, F_0_ was background-corrected, where fluorescence background was measured from a region adjacent to the dendrite. Reported transients had amplitudes more than 3 times the s.d., which was on average ∼15% Δ*F/F_0_* for iAChSnFR and ∼35% Δ*F/F_0_* for GCaMP6s.

### Local in vivo drug delivery during imaging

Tetrodotoxin (TTX) citrate (Tocris, no. 1069) and eserine (Sigma-Aldrich, no. E8375) were applied to the surface of M1 after removing a small bone flap (∼200 μm in diameter) adjacent to a thinned skull window. TTX (100 nM) and eserine (100 μM) were dissolved directly in ACSF to final concentrations. The bone flap for drug delivery was made during awake head mounting and covered with a silicone elastomer such that it could be easily removed at the time of imaging. Because small molecules diffuse rapidly in the cortex, we estimated that the drug concentration was reduced ∼10-fold in the imaged cortical region, such that the final effective concentration would be ∼10 nM for TTX and ∼10 μM for eserine.

### Visual cortex imaging during NB stimulation

Surgery and imaging preparation was performed as in ^38^. Line-scan two-photon excitation was performed by sweeping a 156 μm long reticle focus (420 nm wide, 1.6 μm deep full width at half-maximum) across the sample plane at 1016 Hz. Emitted light was collected with a silicon photomultiplier detector (Hamamatsu MPPC). This microscope uses a 1030 nm Yb:YAG fiber laser for excitation (BlueCut, Menlo Sytems, 190 fs pulses, repetition rate 5 MHz). In **Fig. 7**, F_0_ was calculated by averaging the photon count over the 300 ms prior to stimulus onset, for each trial. Line-scan two-photon excitation was performed by sweeping four 156 μm long reticle foci across the sample plane at four different angles, at 1016 Hz. The summed-intensity recorded from the entire image plane is reported. The effect of fluorophore bleaching was removed by subtracting a mono-exponential fit to the period 0-500 ms prior to and 1500-1600 ms after stimulus onset. A periodic heartbeat artifact was removed using a notch filter with a stopband of 12.5-13 Hz.

### Visual cortex imaging during short-term anesthesia

Line-scan two-photon excitation was performed by sweeping four 156 μm long reticle foci across the sample plane at four different angles, at 1016 Hz. The summed-intensity recorded from the entire image plane is reported. The effect of fluorophore bleaching was removed by subtracting a mono-exponential fit to the period 0-500 ms prior to and 1500-1600 ms after stimulus onset. A periodic heartbeat artifact was removed using a notch filter with a stopband of 12.5-13 Hz.

### Monitoring cholinergic reward signals in mouse hippocampus

#### Animals

*ChAT*-Cre^+/−^ CB1^fl/fl^ mice (90 days old, both male and female, Jackson Labs) were used for the *in vivo* fiber photometry experiments. All mice were single-housed in a temperature-controlled room on a 12-hour light/dark cycle.

#### Surgery

Mice were anesthetized using isoflurane and 500 nL of AAV2/1.*hSynapsin-1*.iAChSnFR (1.9*10^12^ GC/mL) was unilaterally injected in the CA1 region of the hippocampus (AP −2.2 mm, ML +2.0 mm, and DV −1.6 mm). Optical fibers (Thorlabs, CFMC54L02) were implanted 100 µm above the injection site two weeks later. For the electrically stimulated recordings, the mice were anesthetized using isoflurane and an electrode was positioned near the medial septum (AP + 0.6 mm, ML 0.0 mm, and DV −3.5 mm). While recording, the medial septum was stimulated with twenty 8-msec pulses at 5, 10, 15, or 20 Hz (200 µA intensity; duration 4.0, 2.0, 1.33, and 1.0 sec, respectively) to evoke acetylcholine release in the hippocampus. Each electrical stimulus was supplied n = 3 times.

#### Behavior

Animals recorded during an operant task were food-restricted to 90% of body weight and food trained with 1 hr sessions of a fixed-ratio (FR) schedule of reinforcement. During the FR schedule, animals received one sucrose pellet reward for every correct press of the active lever. Following food training, animals were switched to a progressive-ratio (PR) schedule of reinforcement where to earn each incremental reward, the number of active lever presses required increases exponentially (5 × e^(0.2 × reward number) – 5]^). Each PR session ended when the animals failed to reach the number of active lever press required for the reward within 20 mins. The breakpoint, or the total number of rewards an animal received during a session, was considered the metric of motivation for the reward. Photometric recordings were performed during the duration of the PR task following 3 consecutive sessions with the same breakpoint (± 1 reward).

#### Photometry recordings

All photometry recordings were performed using a TDT RZ5P Fiber Photometry processor, Doric Fluorescence minicube (FMC4_AE(405)_E(460-490)_F(500-550)_S), 2151 Femtowatt Silicon Photoreceiver, and TDT Synapse (2015 Edition) software. Data was analyzed using MATLAB (Mathworks, Edition 2018a) with the TDTMatlabSDK tool package provided by Tucker-Davis Technologies.

#### Immunohistochemistry

For the immunostaining, animals were anesthetized using isoflurane and perfused using 1 M phosphate-buffered saline (PBS) followed by 4% paraformaldehyde (PFA). The brains were removed and placed overnight in 4% PFA, washed with 1 M PBS, and cryoprotected in a 30% (w/v) sucrose solution. Using a cryostat, the brains were sectioned at a thickness of 40 microns. The slices were incubated with DAPI for 10 mins, washed with 1 M PBS, and mounted. Images of the hippocampus were taken using a Leica SP8 confocal microscope.

### Statistical tests

Data are presented as mean with standard error (std. err.) or standard deviation (std. dev.), as noted in each case. All values of *n* are provided; no data were excluded. For comparisons between two datasets, a two-sided Student’s t-test was used, unless otherwise noted.

## Acknowledgements

We would like to thank Deepika Walpita (Janelia) for rat neuronal culture, and Kim Ritola (Janelia) for making AAV virus. We thank Jens T. Kaiser at the Caltech Molecular Observatory and the beamline staff at SSRL 12-2 for their support during data collection. DD and HK thank Dr. Martin Fuhrmann (DZNE, Bonn, Germany) for help with virus injections.

This work was supported by the Howard Hughes Medical Institute. JJZ is supported by NIH NS104670, PZ by an Alzheimer’s Association research fellowship, CPF by NIH NS95809, PKZ by a 2019 International Science and Engineering Fair award, WSZ by a 2019 Harrison undergraduate research award, and GW by Epilepsy Foundation postdoctoral fellowship No. 310443. JJZ is the Radboud Professor and Sir Yue-Kong Pao Chair Professor. HAL and AVS are supported by NIH DA037161, NIH DA043829, California Tobacco-Related Disease Research Program 23XT-0007 and 27IP-0057, and NARSAD. JSD is supported by R01-GM095674. JFC is supported by NIH DA022340, and AF by a STAR-PREP fellowship. LL is supported by a National Natural Science Foundation of China grant NSFC81771284. We gratefully acknowledge the Gordon and Betty Moore Foundation and the Beckman Institute for their generous support of the Molecular Observatory at Caltech. Use of the Stanford Synchrotron Radiation Lightsource, SLAC National Accelerator Laboratory, is supported by the U.S. Department of Energy, Office of Science, Office of Basic Energy Sciences under Contract No. DE-AC02-76SF00515. The SSRL Structural Molecular Biology Program is supported by the DOE Office of Biological and Environmental Research, and by the NIH, NIGMS (P41GM103393). BSK is supported by NIH (NS111583, DA047444, MH104069). DD and HK are supported by DFG (SPP1757, SFB1089, DI853/3-5) and BONFOR to DD. WBG is supported by NIH R01 NS047325. ES is supported by JSPS KAKENHI #15KK0340.

## Author contributions

P.M.B., P.Z., H.A.L., J.J.Z., and L.L.L. led the project. P.M.B., A.V.S., J.S.M., H.A.L., and L.L.L. performed protein engineering. P.M.B., A.V.S., and J.S.M. performed biochemical characterization. P.M.B., J.S.M., C.F., D.C.R. acquired and refined crystal structures. P.M.B., A.A., and R.P. performed spectroscopy. P.M.B. and J.S.M. performed HEK293 and cultured neuron experiments. P.Z. and Y.Z. screened sensor variants in brain slice. P.Z., P.K.Z., Y.Z., W.S.Z., G.W., and B.X. acquired and analyzed nanoscopic imaging data. P.Z., Y.Z., L.G., G.-X.Z., K.G., and L.L. made Sindbis and lentivirus constructs and live virus. P.Z. and Y.Z. performed other slice characterization. H.K. and D.D. performed hippocampal stratum radiatum slice characterization. E.S. and B.S.K. performed hippocampal astrocyte slice characterization. Y.C., A.G.Y., and C.P.F. performed striatal slice characterization. J.S.D. acquired and analyzed the *C. elegans* data. C.D. and V.J. acquired and analyzed the *Drosophila* data. M.T. and M.K. acquired and analyzed the zebrafish data. O.N. and K.P. did SLAP imaging and experiments on the effects of anesthesia. J.C. and W.-B.G. did motor cortex experiments. A.F. and J.F.C. did hippocampal reward experiments. P.M.B., H.A.L., J.J.Z., and L.L.L. wrote the paper, with help from all authors.

## Code availability

All analysis code used in this study is available upon request.

## Data availability

All data from this study is available upon request.

## Accession codes

All constructs have been deposited at Addgene (#137950-137959). Sequences have been deposited in Genbank. PDB accession code for iAChSnFR0.4 is 6URU. PDB accession code for the binding protein-only component of iAChSnFR is 6V1R. AAV virus is available from Addgene.

---

**Supplementary Fig. S1.**
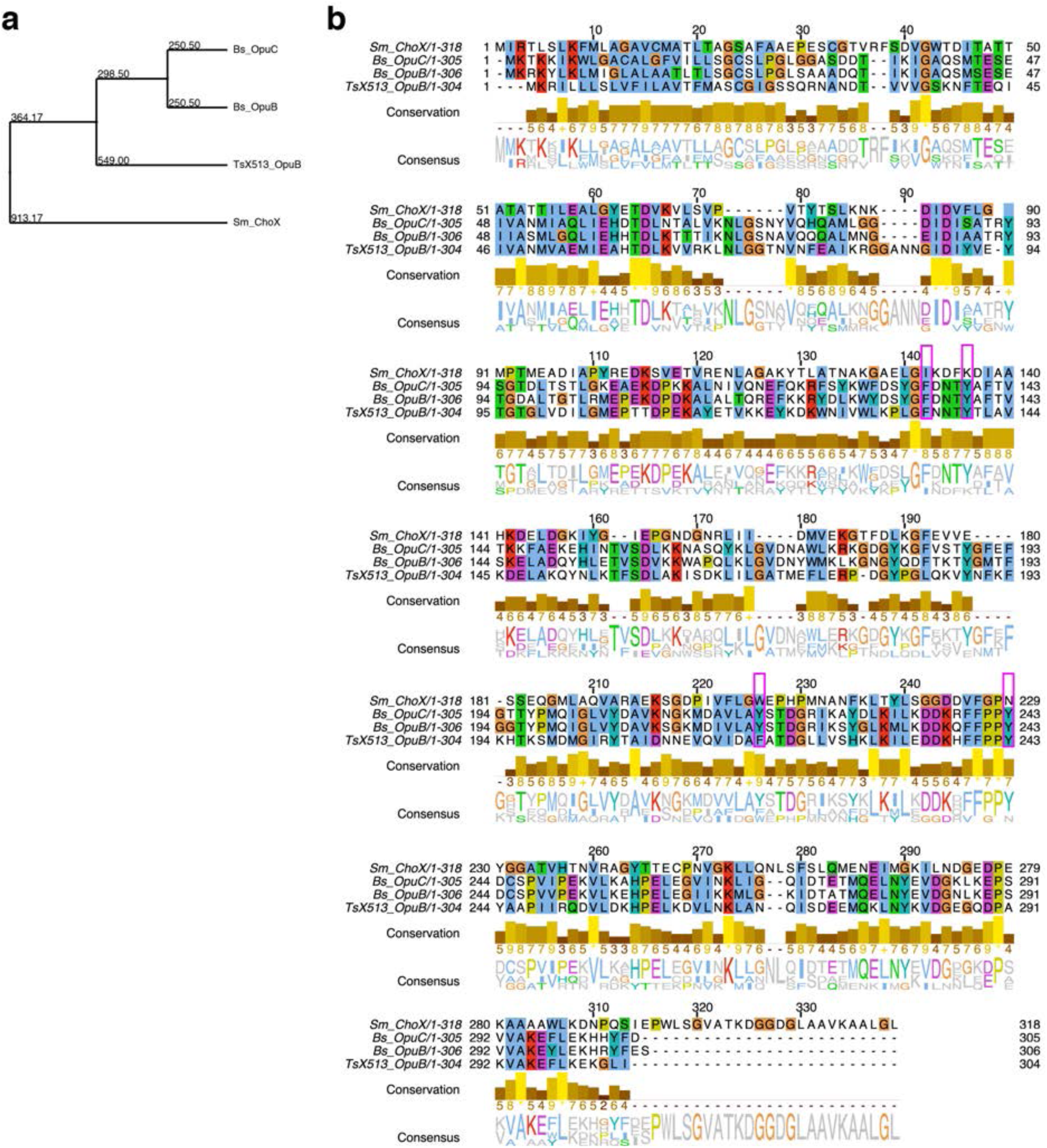
Sequence analysis of the various PBP sequences used in this study. (**a**) Phylogenic tree relating the four sequences. Numbers represent branch length. (**b**) ClustalW alignment of the four sequences. The four key aromatic residues (Tyr94, Trp219, and Tyr243) involved in cation-pi interactions are highlighted in magenta boxes; the last three correspond to residues labeled γ, ε, and η in ref. 34, respectively. Sm = *Sinorhizobium meliloti*, Bs = *Bacillus subtilis*, TsX513 = *Thermoanaerobacter sp. X513*.

**Supplementary Fig. S2.**
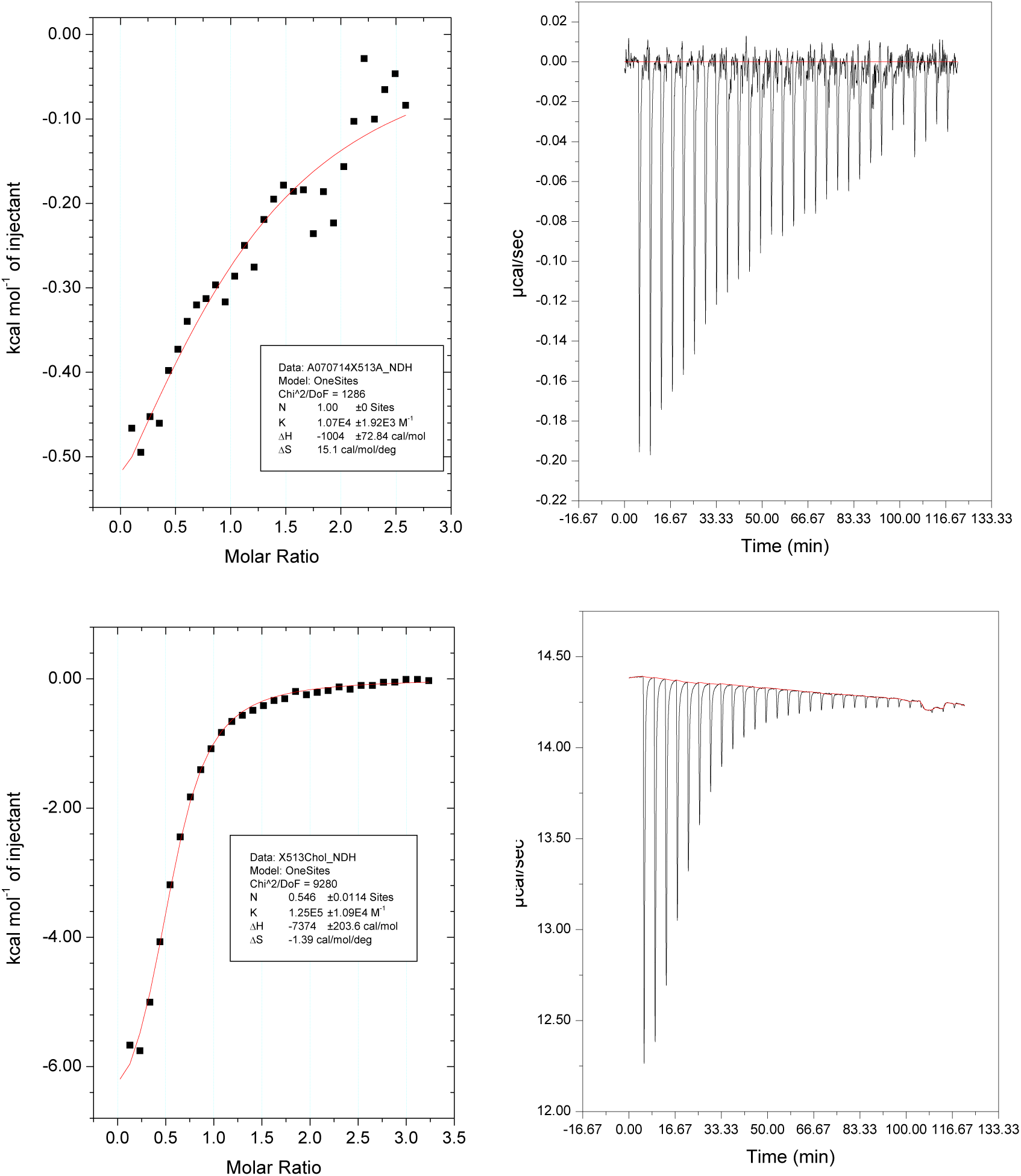
Isothermal titration calorimetry analysis of *Thermoanaerobacter sp. X513* OpuBC homologue. (*Top*) 100 μM purified protein was titrated with 1 mM ACh at 25°C in PBS. Curve fitting with a fixed stoichiometry (N) of 1 yields a calculated *K*_d_ for ACh of 95 µM (red curve; Chi^2^ for fit = 1300). Curve fitting with a variable N yields a calculated *K*_d_ of 65 µM with N=1.22 (not shown). (*Bottom*) 80 µM purified protein was titrated with 1.5 mM choline at 25°C in PBS. Curve fitting with variable stoichiometry yields a calculated *K*_d_ of 8 µM and a stoichiometry of 0.55 (red curve; Chi^2^ for fit = 9300). Each experiment was performed once.

**Supplementary Fig. S3.**
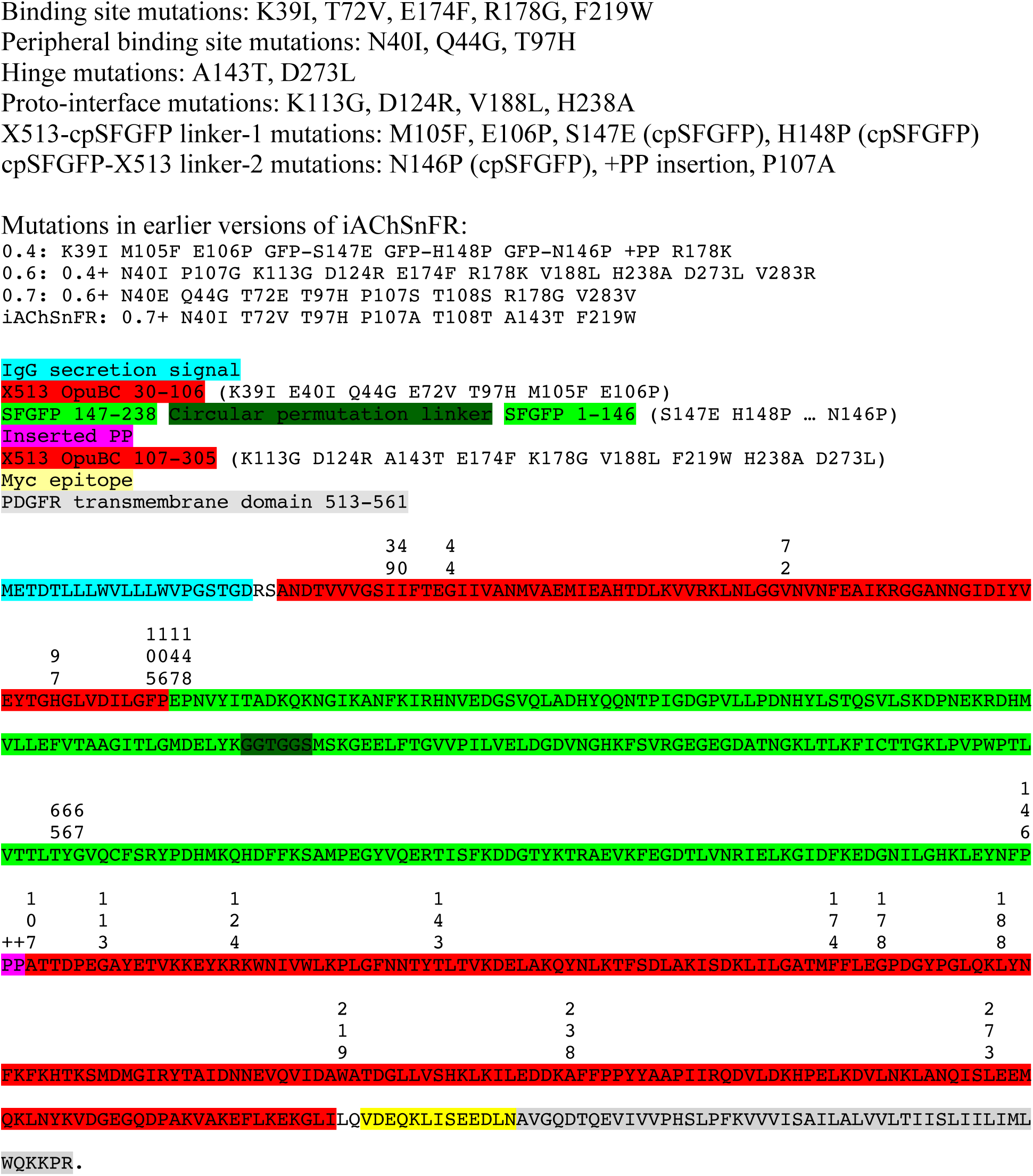
Annotated amino acid sequence of iAChSnFR and variants. Insertion of cpSFGFP is after residue E106 of X513. Residues RS near the N-terminus encode a BglII restriction site, and residues LQ at the C-terminus encode a PstI restriction site. Domains colored. Amino acid numbering matches normal N-to-C numbering of each domain. X513 OpuBC starts at 30 (the first 29 amino acids are an endogenous secretion signal, which is omitted). cpSFGFP is numbered from 147-238.GGTGGS.1-146; the chromophore (TYG) is numbered 65-67.

**Supplementary Fig. S4.**
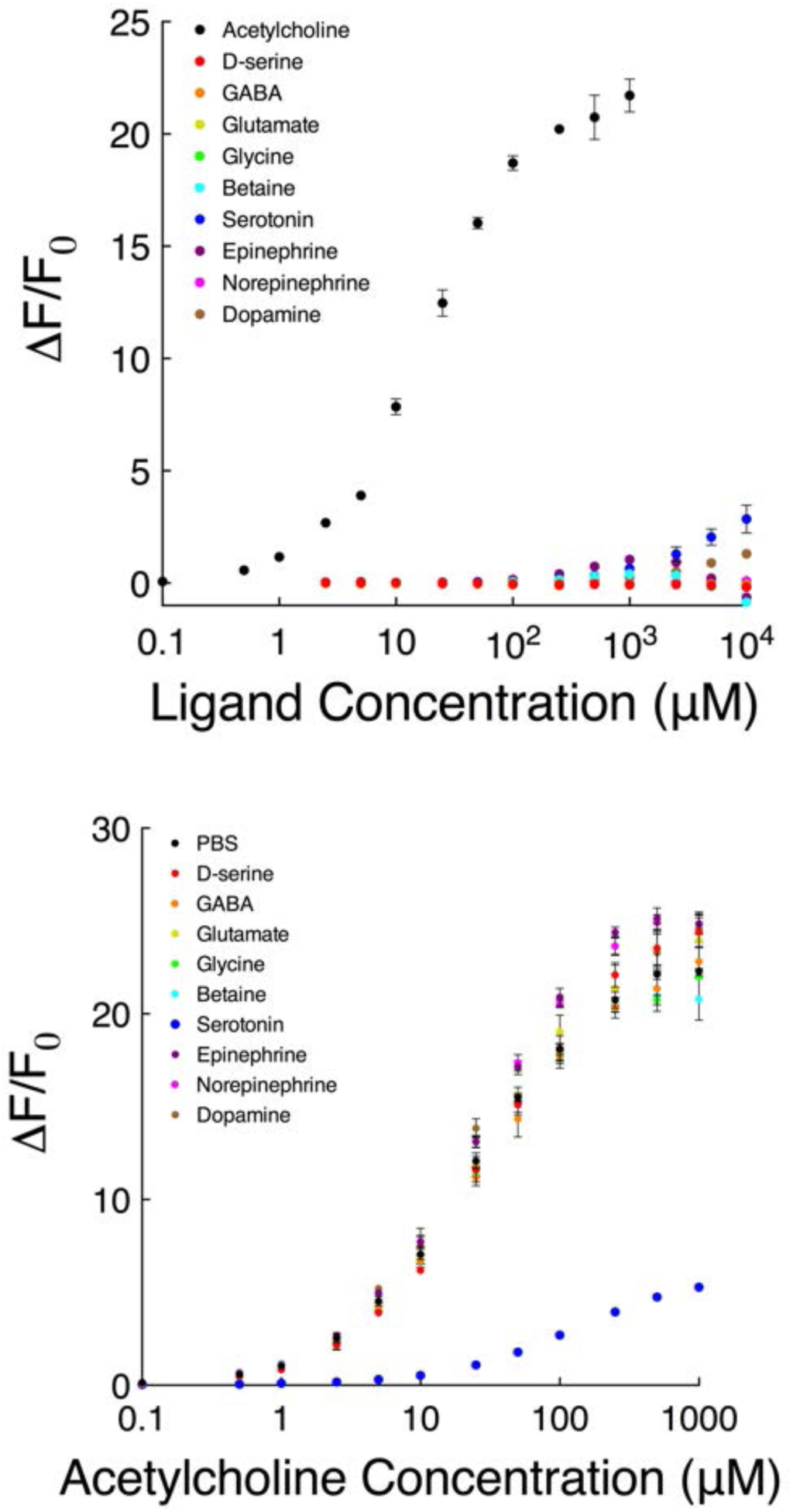
Top, responses of iAChSnFR to other known and putative neurotransmitters; 200 nM iAChSnFR was titrated with various concentrations of the ligands tested. Bottom, tests of the effects of ligands on the response of iAChSnFR to ACh; 200 nM iAChSnFR was titrated with acetylcholine in the presence of 1 mM of each of the ligands listed. Mean of n=3 technical replicates, error bars show std.dev.

**Supplementary Fig. S5.**
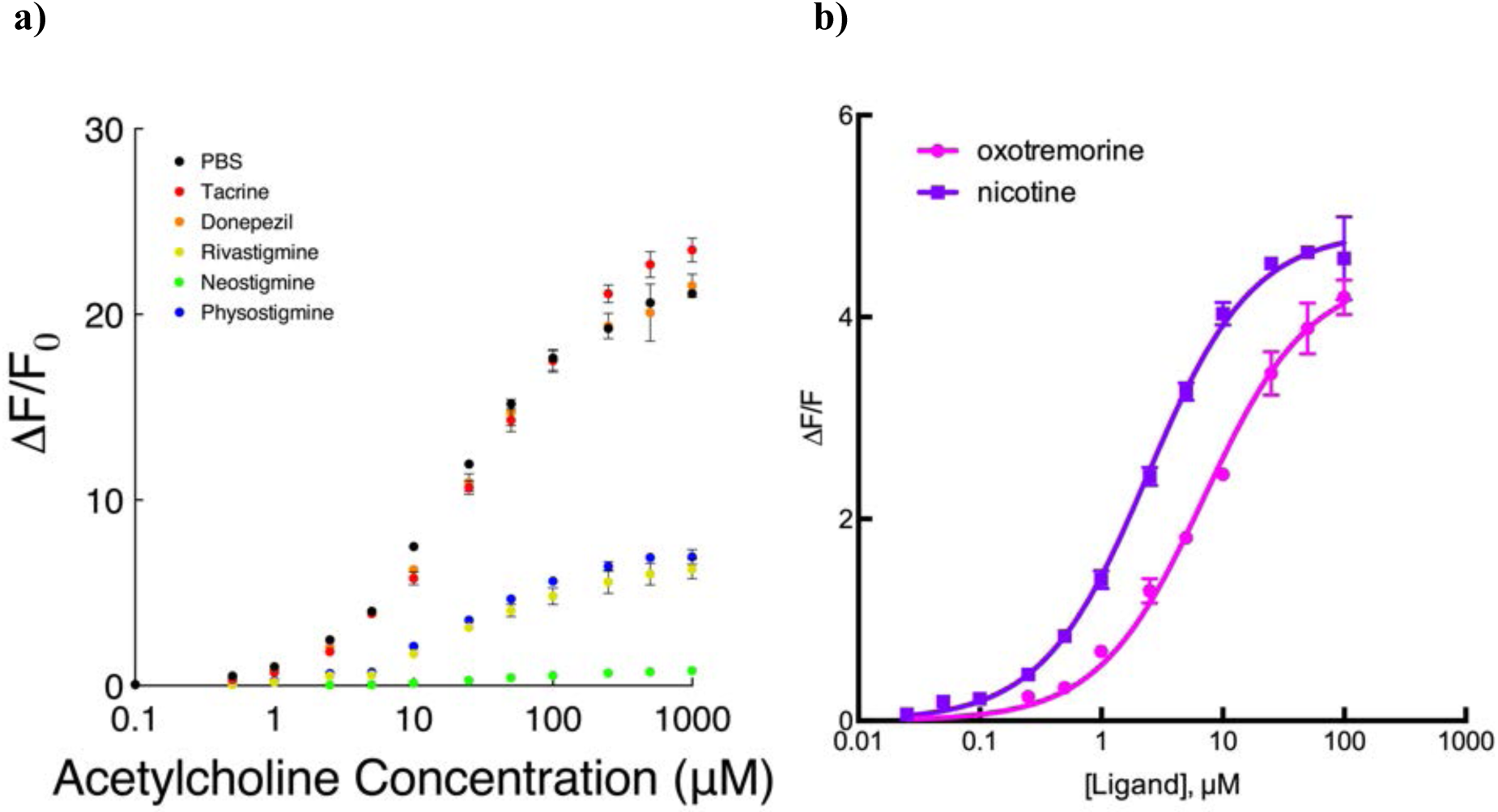
a) Influence of acetylcholinesterase inhibitors on iAChSnFR. 200 nM iAChSnFR0.7 was titrated with various concentrations of acetylcholine in the presence of 50 μM of each of the acetylcholinesterase inhibitors listed. Mean of n=3 technical replicates, error bars show S.D. b) Affinity of iAChSnFR for the nAChR and mAChR agonists nicotine and oxotremorine, respectively. 200 nM iAChSnFR was titrated with various concentrations of each compound. Mean of n=3 technical replicates, error bars are std.dev.

**Supplementary Fig. S6.**
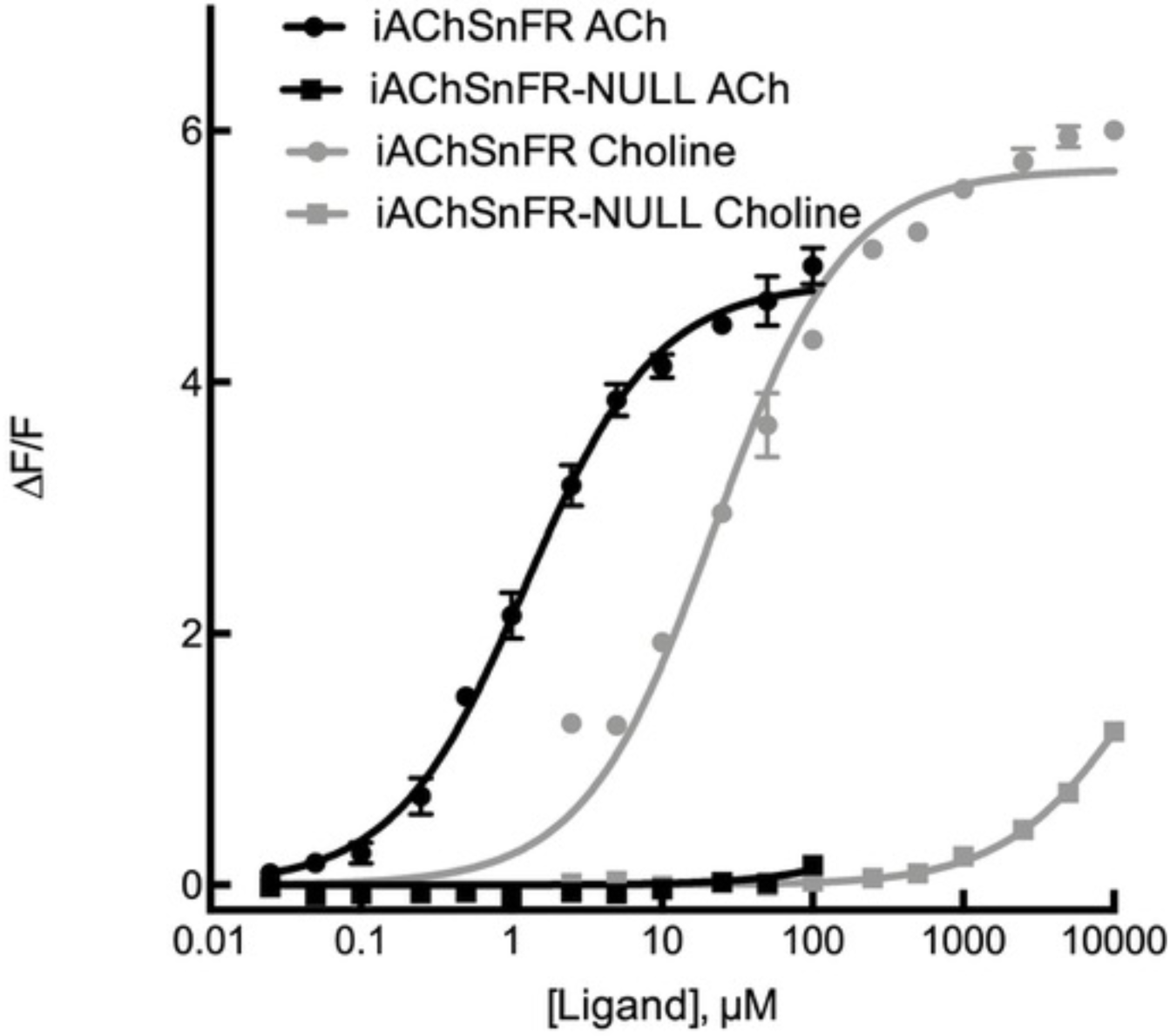
Responses of iAChSnFR and the NULL variant to acetylcholine or choline. 200 nM iAChSnFR was titrated with various concentrations of the two compounds. Mean of n = 3 technical replicates, error bars are s.e.m.

**Supplementary Fig. S7.**
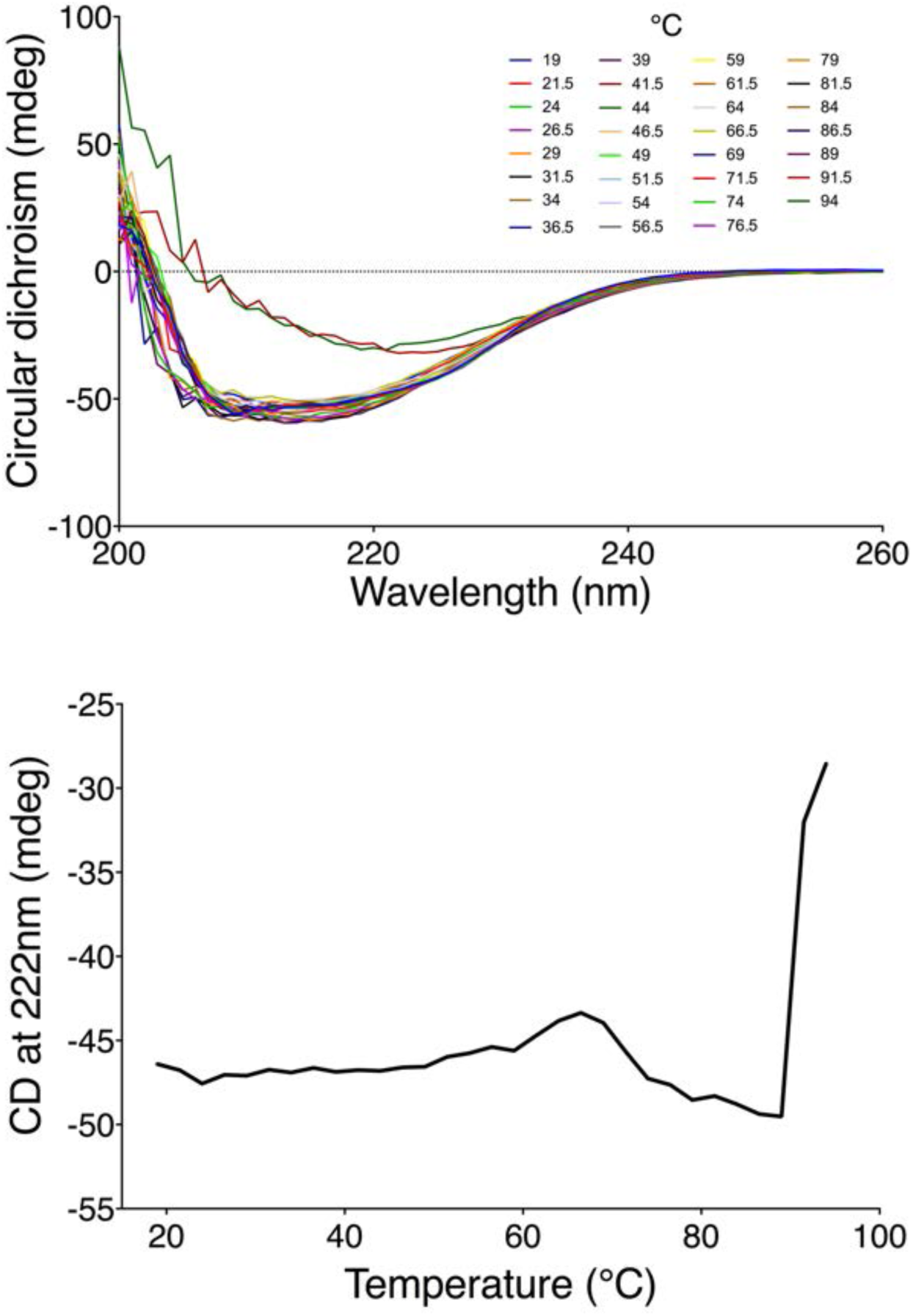
Circular dichroism (CD) measurements of iAChSnFR. Top, 1 μM of iAChSnFR was thermally denatured in 0.1X PBS. Note that the upper temperature limit for the instrument used is 95°C. Bottom, plot of CD signal at 222 nm as a function of temperature.

**Supplementary Fig. S8.**
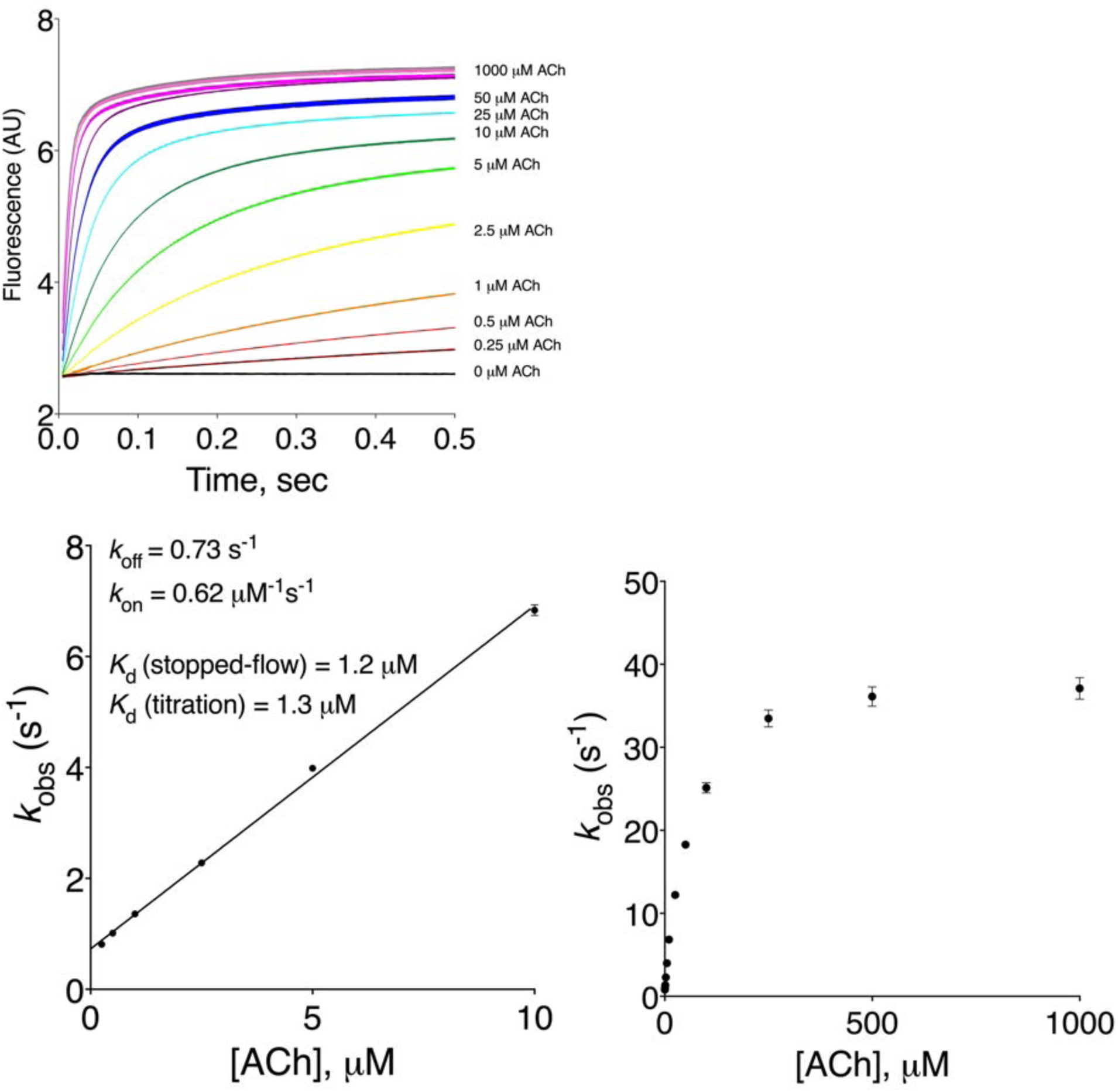
Stopped-flow kinetic analysis of iAChSnFR. Top, 200 nM iAChSnFR was mixed with varying concentrations of acetylcholine at 25⁰C to produce the indicated final [ACh] values. Bottom, *k*_obs_ plot as a function of concentration; left panel, linear fit of the data at ≤ 10 μM. Fitted kinetic parameters: *k*_on_ = 0.62 μM^-1^s^-1^, *k*_off_ = 0.73 s^-1^, apparent *K*_d_ (stopped-flow) = *k*_off_ /*k*_on_ = 1.2 μM. Mean of n = 3 technical replicates, error bars show s.e.m.

**Supplementary Fig. S9.**
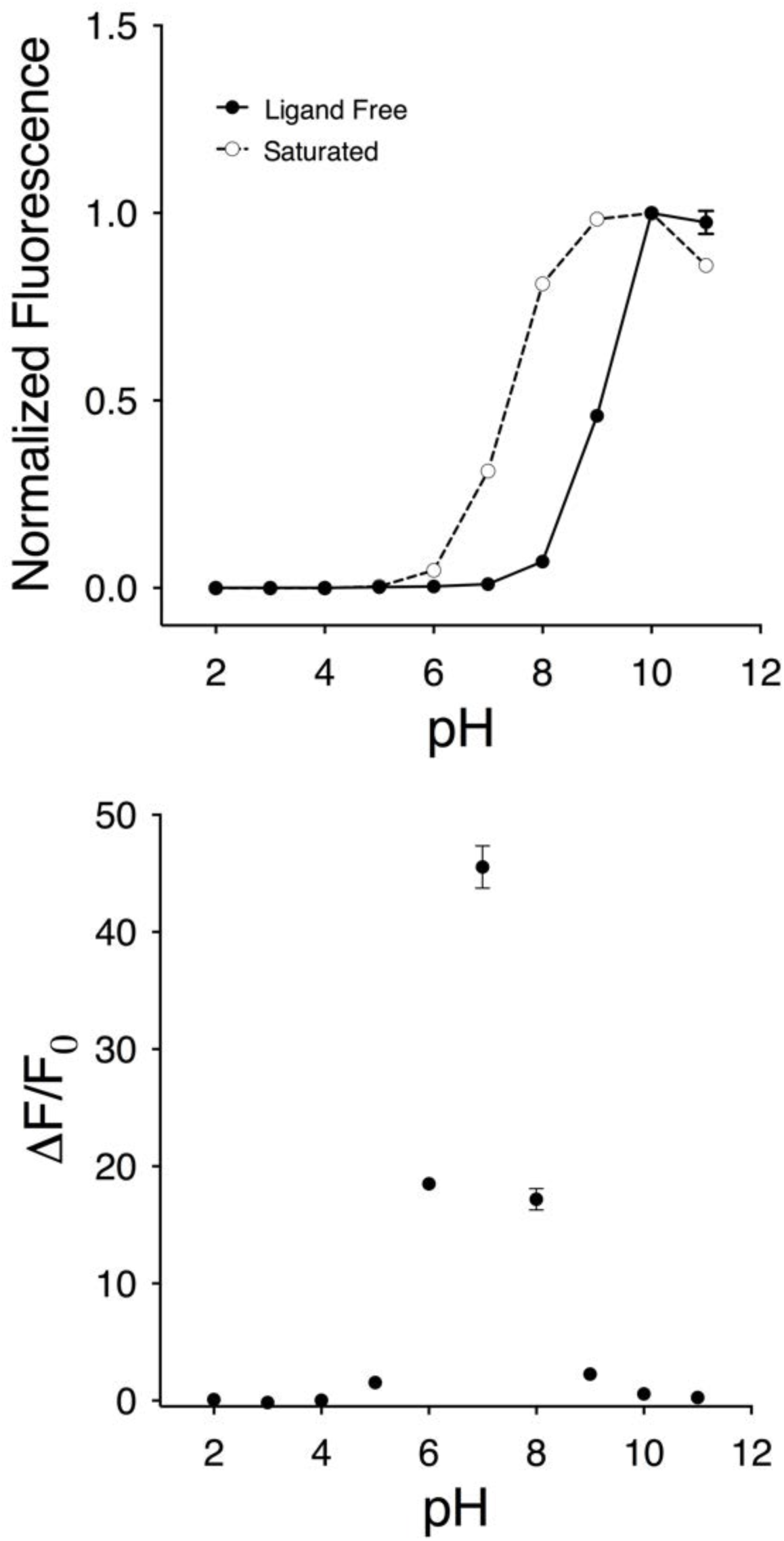
pH titrations of iAChSnFR. Top, fluorescence of the acetylcholine-saturated (open circles, 100 μM acetylcholine) and -free (circles) normalized to peak fluorescence of iAChSnFR0.7. Bottom, measurement of ΔF/F_0_ as a function of pH. Measurements were made using 200 nM sensor, in colorless Hydrion buffers. Mean of n=3 technical replicates, error bars show std.dev.

**Supplementary Fig. S10.**
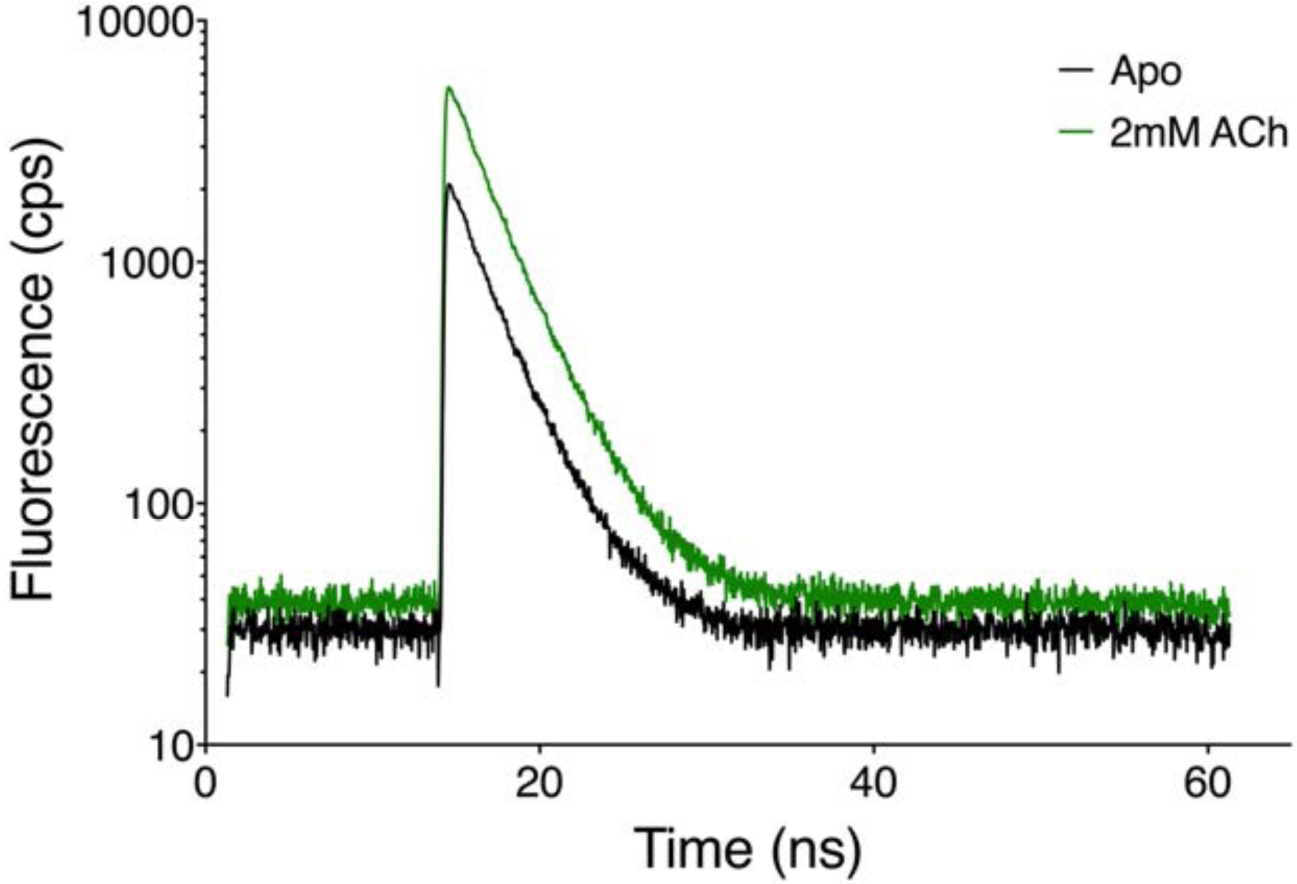
Fluorescence lifetime characterization of iAChSnFR. Two-photon fluorescence lifetime measurement of iAChSnFR in both apo and acetylcholine-saturated states. Single exponential curve fits show a lifetime of 2.59 nanoseconds for the apo form and 2.58 nanoseconds for the acetylcholine-bound form. Lifetime measurements were taken at 950 nm excitation wavelength at 10 mW power with a neutral density (ND 0.5) filter placed in the excitation path.

**Supplementary Fig. S11.**
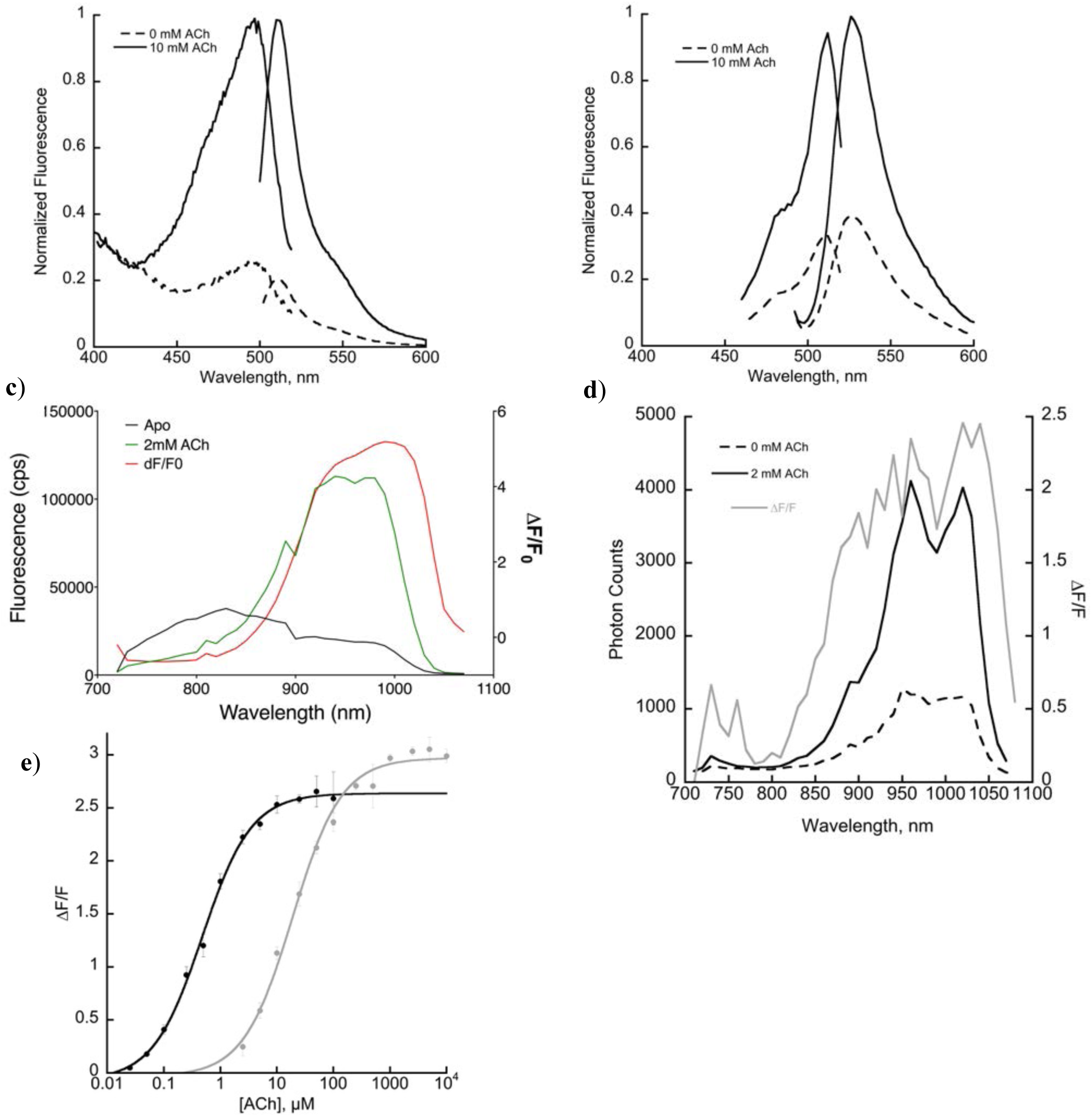
**Top**. One-photon fluorescence spectra of iAChSnFR and y-iAChSnFR. **a)** iAChSnFR: Excitation (5 nm bandwidth) was measured at an emission wavelength of 650 nm/20 nm bandwidth; emission (5 nm bandwidth) was measured at excitation wavelength of 485 nm/2.5 nm bandwidth, with 1 μM iAChSnFR in PBS + 1 mM acetylcholine (solid traces) or PBS alone (dashed traces). **b)** y-iAChSnFR: Excitation (5 nm bandwidth) was measured at an emission wavelength of 555 nm (5 nm bandwidth); emission (5 nm bandwidth) was measured at excitation of 470 nm/5 nm bandwidth, with 1 μM iAChSnFR in PBS + 10 mM acetylcholine (solid traces) or PBS alone (dashed traces). **Middle.** Two-photon peak brightness spectra, along with calculated ΔF/F at each wavelength. 2P spectra obtained at 1 mW power across 720-1080 nm, fluorescence collected with a 550BP88 filter. **c)** iAChSnFR: apo (dashed) and acetylcholine-bound (solid) sensor. **d)** y-iAChSnFR: apo (dashed) and ACh-bound (solid) sensor. (**e)** Titration of y-iAChSnFR with ACh (black, *K*_d_ = 500 nM) and choline (gray, *K*_d_ = 20 μM). Mean of n=3 technical replicates, error bars show std.dev.

**Supplementary Fig. S12.**
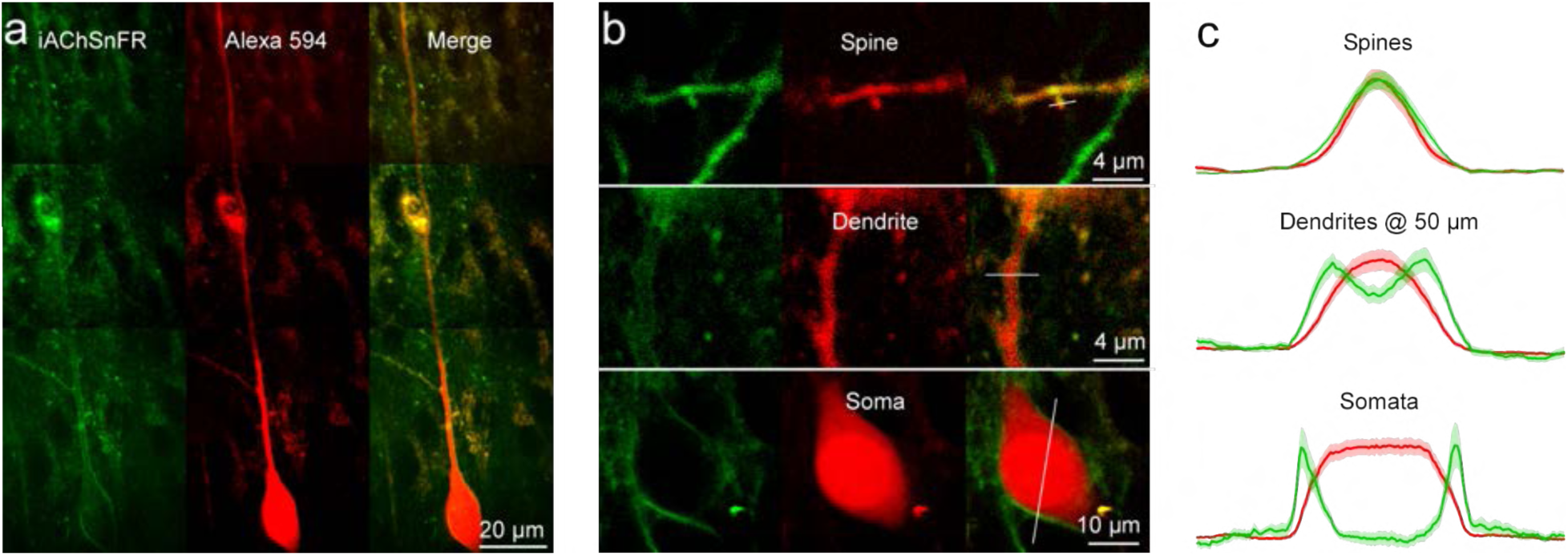
iAChSnFR displays well on the membrane of CA1 neurons in brain slice. (**a**) Two-photon montage of images (green, EGFP channel; red, AlexaFluor 594 channel) of an iAChSnFR-expressing CA1 neuron pipette-loaded with 5 µM AlexaFluor 594. Slice thickness is 400 µm. (**b**) Representative two-photon images of: top) a dendrite and spine, middle) the dendrite 50 µm away from the soma, and bottom) the soma of another iAChSnFR-expressing CA1 neuron. Also shown are the sites of line scans for quantification in (c). (**c**) Means and std. err. of normalized green and red fluorescence measured from line scans across somata (*n* = 14 from 14 neurons of 12 animals), dendrites ∼50 µm away from the soma (*n* = 17 from 17 neurons of 12 animals), and spines (*n* = 25 from 15 neurons of 12 animals) of iAChSnFR-expressing CA1 neurons.

**Supplementary Fig. S13.**
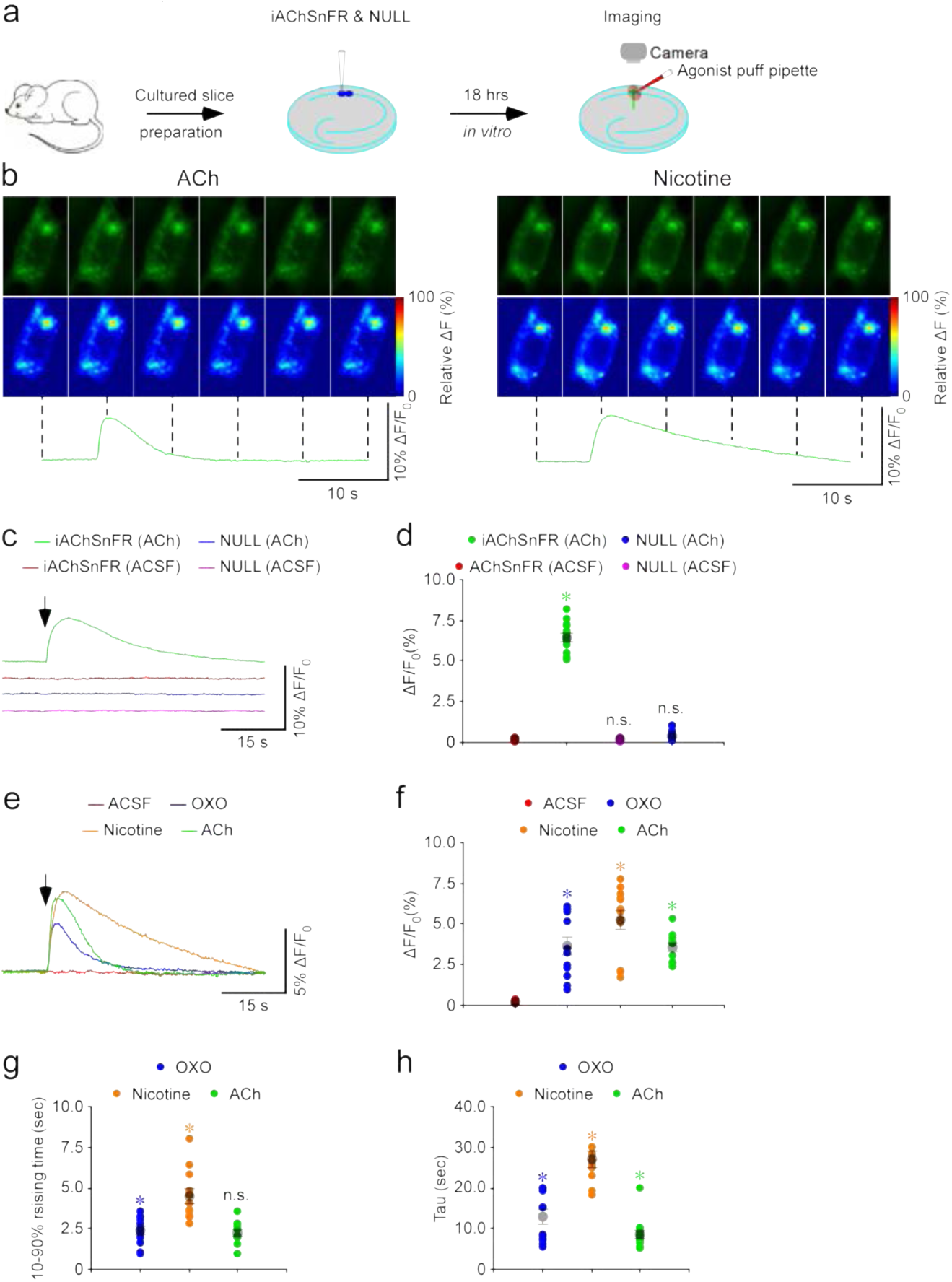
Response of iAChSnFR in hippocampal CA1 slice to puffs of ACh. (**a**) Schematic of the experiments. (**b**) Snapshots of fluorescence images (upper) and relative fluorescence responses in heat map format (lower) of iAChSnFR-expressing CA1 neurons to a 500-ms puff of ACh (1 mM, left) or nicotine (1 mM, right). (**c**) Fluorescence responses (offset vertically, all have same baseline) of iAChSnFR and iAChSnFR-NULL to a 500-ms puff of ACSF or 1 mM acetylcholine (ACh). (**d**) Values for fluorescence responses from (c) (ACh: 6.4±0.2%, ACSF: 0.2±0.02%, *Z=*3.7, *p*<0.001 for iAChSnFR; ACh: 0.3±0.06%, ACSF: 0.1±0.02%, *Z=*3.0, *p*<0.005 for iAChSnFR-NULL*; n*=18 from 5 animals). (**e**) Waveforms of fluorescence responses of iAChSnFR to a 500-ms puff of ACSF, 1 mM oxotremorine (Oxo), 1 mM nicotine (Nic) or 1 mM ACh. (**f, g, h**) Analysis of waveforms in **e**. (**f**) Peak fluorescence response (Oxo: 3.6±0.6%, *Z=*3.1, *p<*0.005; Nic: 5.2±0.6%, *Z=*3.1, *p<*0.005; ACh: 3.5±0.3%, *Z=*3.6, *p*<0.005; ACSF: 0.2±0.02%, *n*=12 from 5 animals). (**g**) 10-90% rise time (Nic: 4.6±0.5 s, *Z*=2.8, *p<*0.01; Oxo: 2.2±0.2 s, *Z=*-1.1, *p*=0.26; ACh: 2.4±0.2 s, *n*=12 from 5 animals). (**h**) Decay time constant (tau; Nic: 27.0±2.0 s, *Z=*2.8, *p<*0.01; Oxo: 12.6±1.8 s, *Z=-* 2.1, *p*<0.05; ACh: 8.4±1.1 s, *n*=12 from 5 animals). Large gray dots indicate average parameters; asterisks indicate *p*<0.05 (Wilcoxon rank-sum test).

**Supplementary Fig. S14.**
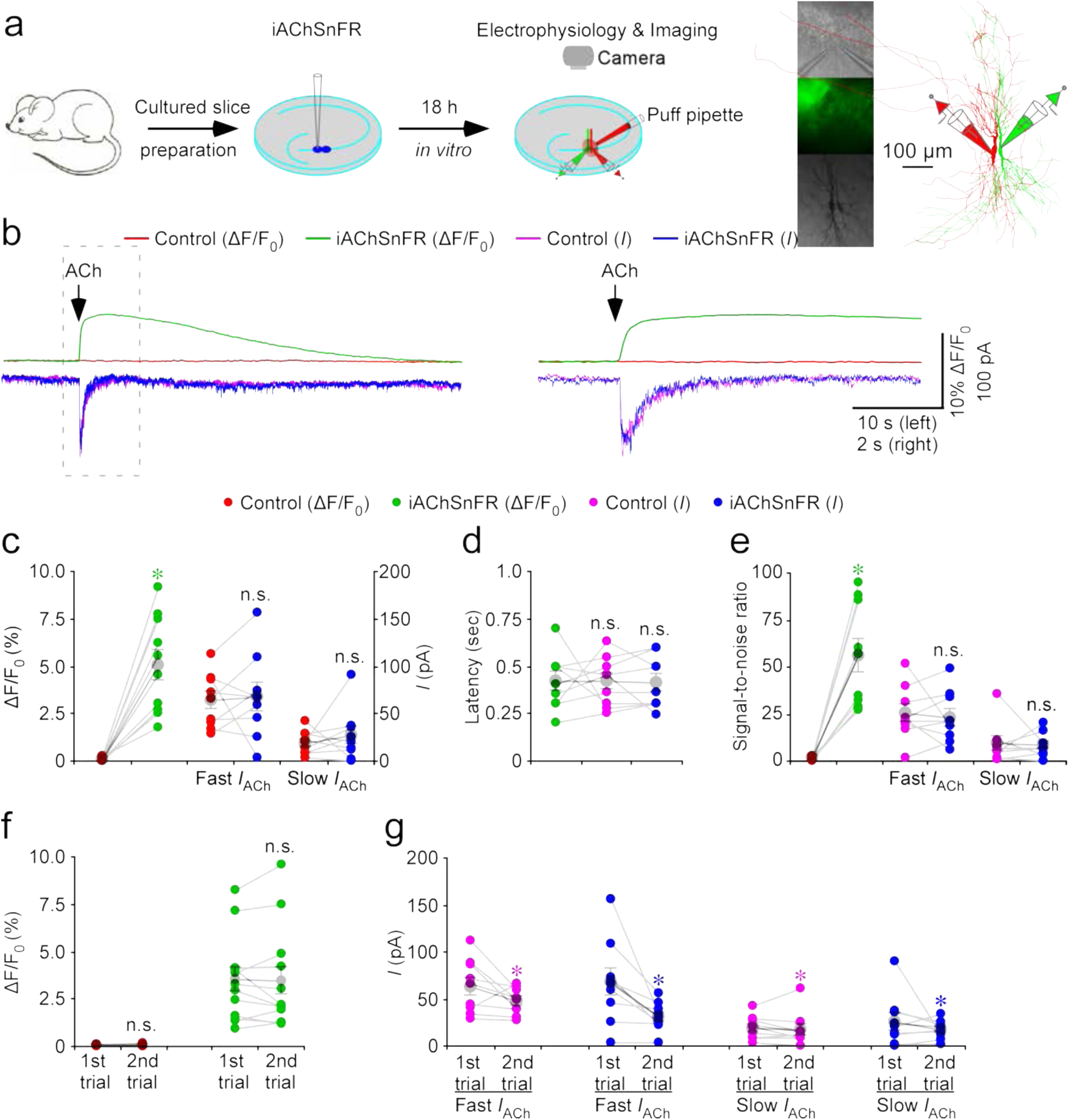
Expression of iAChSnFR does not detectably affect ACh-induced currents in hippocampal CA3 slices. (**a**) Schematic of simultaneous imaging and electrophysiological recording experiments. (**b**) Left, simultaneous fluorescence and current responses of a pair of neighboring CA3 neurons. The neuron at left is fluorescent, showing that it expresses iAChSnFR; the neuron at right is non-fluorescent (showing that it does not express iAChSnFR and serves as a control). A 500-ms puff application of 1 mM ACh was applied. Right, the responses in the left dashed box are replotted in an expanded time scale. Note that both the fluorescence and current waveforms begin with latencies < 300 ms, and that ACh-induced currents appeared to consist of fast and slow components, corresponding presumably to nicotinic and muscarinic currents. (**c**) Values for peak ACh-induced fluorescence (iAChSnFR: 4.0±0.6%; Ctrl: 0.1 ± 0.02%; *Z*=2.8; *p*=0.005), and for peak ACh-induced currents in iAChSnFR-expressing neurons compared to non-expressing neurons, for fast (iAChSnFR: 67.2±15.0 pA; Ctrl: 63.5±9.5 pA; *Z*=0.5; *p*=0.59) and slow (iAChSnFR: 26.2±9.2 pA; Ctrl: 19.2±4.0 pA; *Z*=0.8; *p*=0.44) currents (*n*=10 neuron pairs from 5 animals). (**d**) Latencies of ACh-induced currents in iAChSnFR-expressing (iAChSnFR: 411.7±46.3 ms; *Z*=-0.1; *p*=0.92) and non-expressing CA3 neurons (Ctrl: 418.9±43.5 ms; *Z*=0.7; *p*=0.48) compared to those of fluorescence responses of iAChSnFR-expressing neurons (iAChSnFR: 416.7±50.0 ms) (*n*=9 neuron pairs from 4 animals). (**e**) Values for the signal-to-noise ratio (SNR) of cholinergic fluorescence responses of iAChSnFR expressing CA3 neurons compared to non-expressing neurons (iAChSnFR: 56.3±9.1; Ctrl: 1.5±0.3; *Z*=-2.7; *p*=0.008), and that of fast (iAChSnFR: 23.2±4.6; Ctrl: 25.2±4.7; *Z*=-0.06; *p*=0.95) and slow (iAChSnFR: 8.2±2.1; Ctrl: 9.0±3.5; *Z*=-0.2; *p*=0.86) cholinergic current responses of iAChSnFR-expressing CA3 neurons compared to non-expressing neurons (*n*=9 neuron pairs from 4 animals). Note that SNR of cholinergic fluorescence responses of iAChSnFR-expressing CA3 neurons is larger than both fast (iAChSnFR: *Z*=-2.7; *p*=0.008; Ctrl: *Z*=-2.2; *p=*0.028) and slow (iAChSnFR: *Z*=-2.7; *p*=0.008; Ctrl: *Z*=-2.2; *p=*0.028) cholinergic current responses of iAChSnFR-expressing and non-expressing CA3 neurons. **(f-g)** ACh-induced fluorescence displays less rundown than ACh-induced currents. (**f**) Values for the two consecutive fluorescence responses of iAChSnFR-expressing (1^st^: 0.08±0.01%; 2^nd^: 0.08±0.02%; *Z*=-0.1; *p*=0.88) CA3 neurons (*n*=13 neuron pairs from 5 animals). (**g**) Peak responses for two consecutive fast cholinergic current responses in iAChSnFR-expressing (1^st^: 68.2±12.1 pA; 2^nd^: 34.1±4.7 pA; *Z*=-2.5; *p*=0.011) and control non-expressing (1^st^: 64.7±7.8 pA; 2^nd^: 48.0±3.9 pA; *Z*=-2.0; *p=*0.49) CA3 neurons, and values for the two consecutive slow cholinergic current responses of iAChSnFR-expressing (1^st^: 26.0±7.4 pA; 2^nd^: 16.6±2.8 pA; *Z*=-0.5; *p*=0.59) and control non-expressing (1^st^: 20.4±3.4 pA; 2^nd^: 16.3±4.8 pA; *Z*=-0.9; *p*=0.37) CA3 neurons (*n*=9 neuron pairs from 4 animals). Large gray dots indicate average responses and asterisks indicate *p*<0.05 (Wilcoxon rank-sum tests). Data for panels **c – g** are for *n*=15 cells from 8 animals.

**Supplementary Fig. S15.**
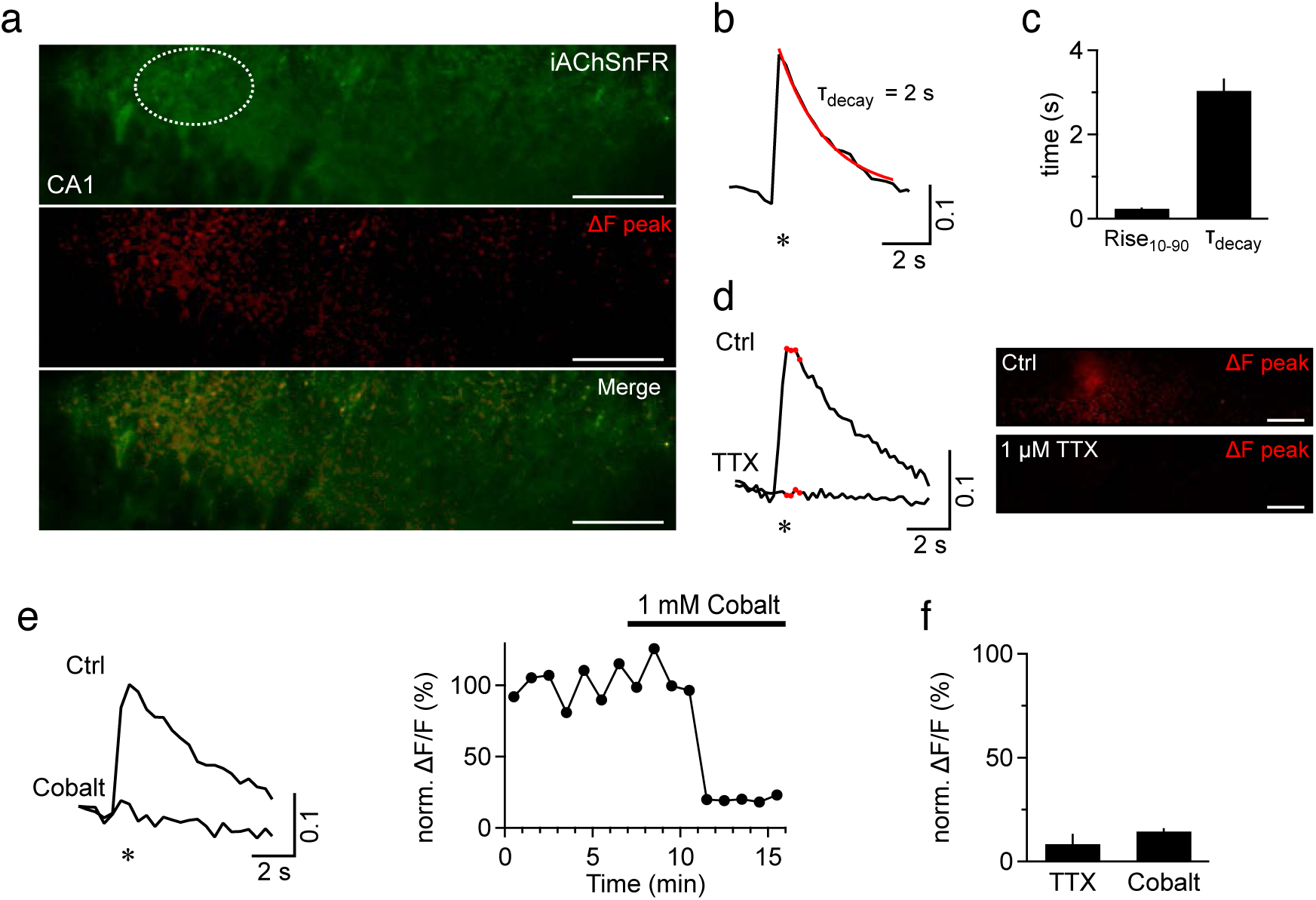
iAChSnFR detects electrically evoked ACh release from presynaptic sites in hippocampal slices. (a) Representative confocal scan of iAChSnFR expression in the CA1 area of an acute hippocampal slice (top panel, mouse 12 weeks). The stimulus was applied from a glass pipette electrode placed in hippocampal CA1 stratum radiatum (2 x 100 µs stimuli, 100 ms interval). The middle panel shows a difference image calculated between the peak and the pre-stimulus fluorescence (after and before the asterisk shown in b)). Note the punctate appearance of the fluorescence increases (ΔF), mostly in stratum radiatum. Lower panel shows an overlay of ΔF and the resting fluorescence shown in the top panel and illustrates that responding puncta localize to structures in the neuropil. Scale bars: 50 µm. (b) Exemplar fluorescence waveform for the dashed ROI shown in a). Asterisk indicates the time of stimulation. The average ΔF/F was 0.24 ± 0.02 (n = 13 from 3 animals). Red trace shows fit to a single exponential. (c) Left, 10-90% rise time (0.7 ± 0.06 s) and right, average decay time constant (3.0 ± 0.3 s) obtained by mono-exponential fitting (n = 13). (d) Left, exemplar waveforms of iAChSnFR response before and after 1 µM TTX application (ΔF peak: average of 4 successive points, red), demonstrating that the responses depend on presynaptic action potential generation. Right, peak ΔF difference image, for control and +TTX. Scale bars: 50 µm. (e) Left panel: representative traces before and after 1 mM CoCl_2_ application to block presynaptic Ca^2+^ entry required for action potential-dependent ACh release. Right panel: time course of the peak of electrically evoked responses showing their reproducibility, stability and responsiveness to the block of Ca^2+^ entry. (f) Summary bar graphs (% of control) of the inhibitory effect of TTX (8.4 ± 4.9%; n = 4) and cobalt (14.6 ± 1.6%; n = 3) on electrically evoked iAChSnFR responses. Error bars show std.err.

**Supplementary Fig. S16.**
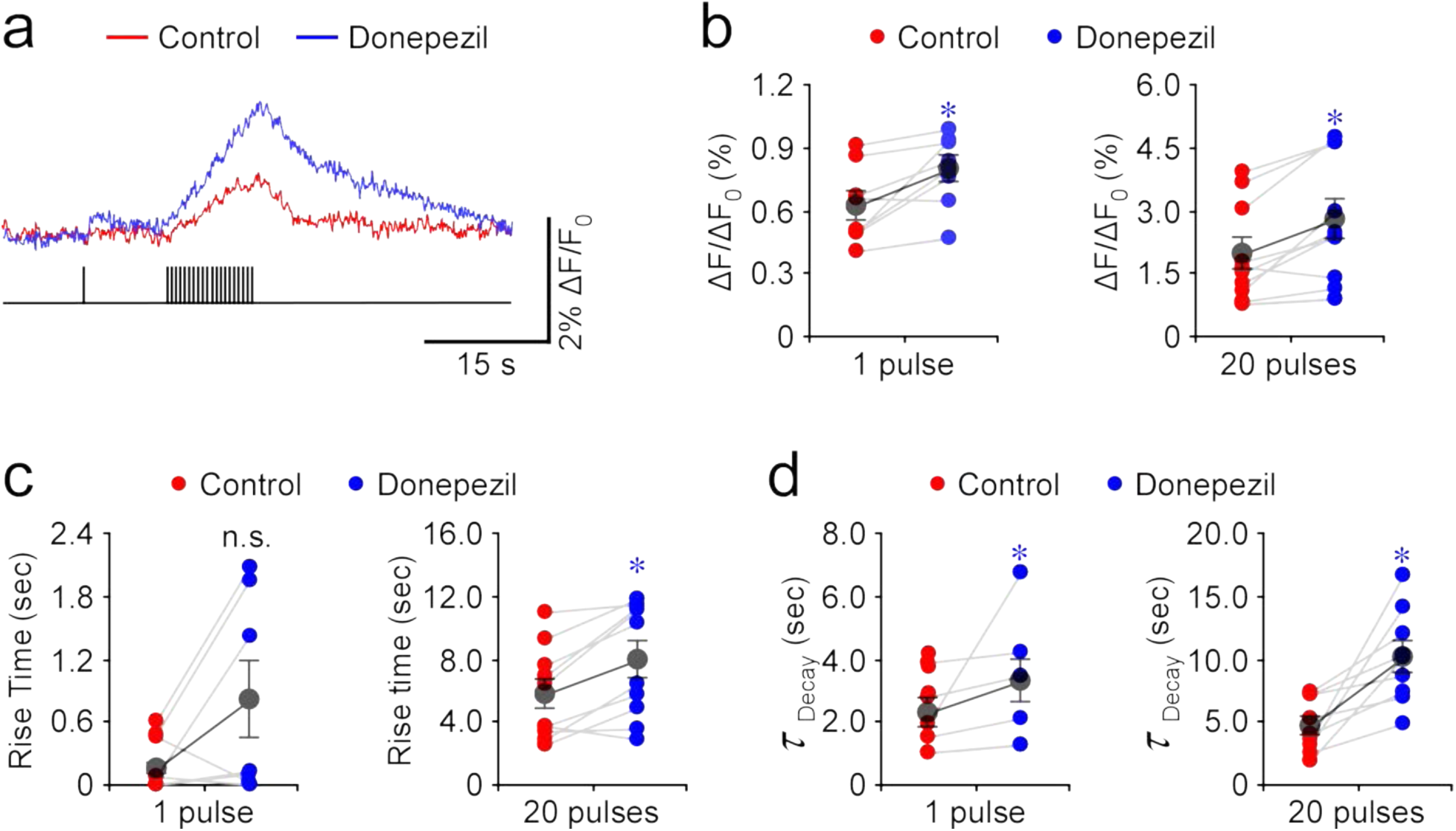
iAChSnFR illustrates the role of AChE in restricting cholinergic transmission. (a) Fluorescence waveforms of iAChSnFR-expressing MEC neurons to a single electrical pulse followed by a train of 20 pulses at 2 Hz in normal ACSF with or without 0.5 μM donepezil. (b) Peak fluorescence responses to single pulses (Ctrl: 0.6±0.07%; Exp: 0.8±0.06%; *Z*=2.380; *p*=0.017; *n*=8 from 8 animals) and 20 pulses (Ctrl: 2.0±0.4%; Exp: 2.8±0.5%; *Z*=2.599; *p*=0.009; *n*=10 from 10 animals) in ACSF with or without 0.5 μM donepezil. (c) The 10-90% rise time of fluorescence responses waveforms for single pulses (Ctrl: 0.2±0.04 ms; Exp: 0.8±0.4 s; *Z*=1.512; *p*=0.128; *n*=7 from 7 animals) and 20 pulses (Ctrl: 5.7±0.9 s; Exp: 7.9±1.1 s; *Z*=2.497; *p*=0.013; *n*=10 from 10 animals) in ACSF with or without 0.5 μM donepezil. (d) The decay time constant of fluorescence responses waveforms for single pulses (Ctrl: 2.2±0.5 s; Exp: 3.3±0.7 s; *Z*=2.203; *p*=0.043; *n*=7 from 7 animals) and 20 electrical pulses (Ctrl: 4.7±0.7 s; Exp: 10.1±1.2 s; *Z*=2.666; *p*=0.008; *n*=9 from 9 animals) in ACSF with or without 0.5 μM donepezil. Large gray dots indicate average responses and asterisks indicate *p*<0.05 (Wilcoxon tests).

**Supplementary Fig. S17.**
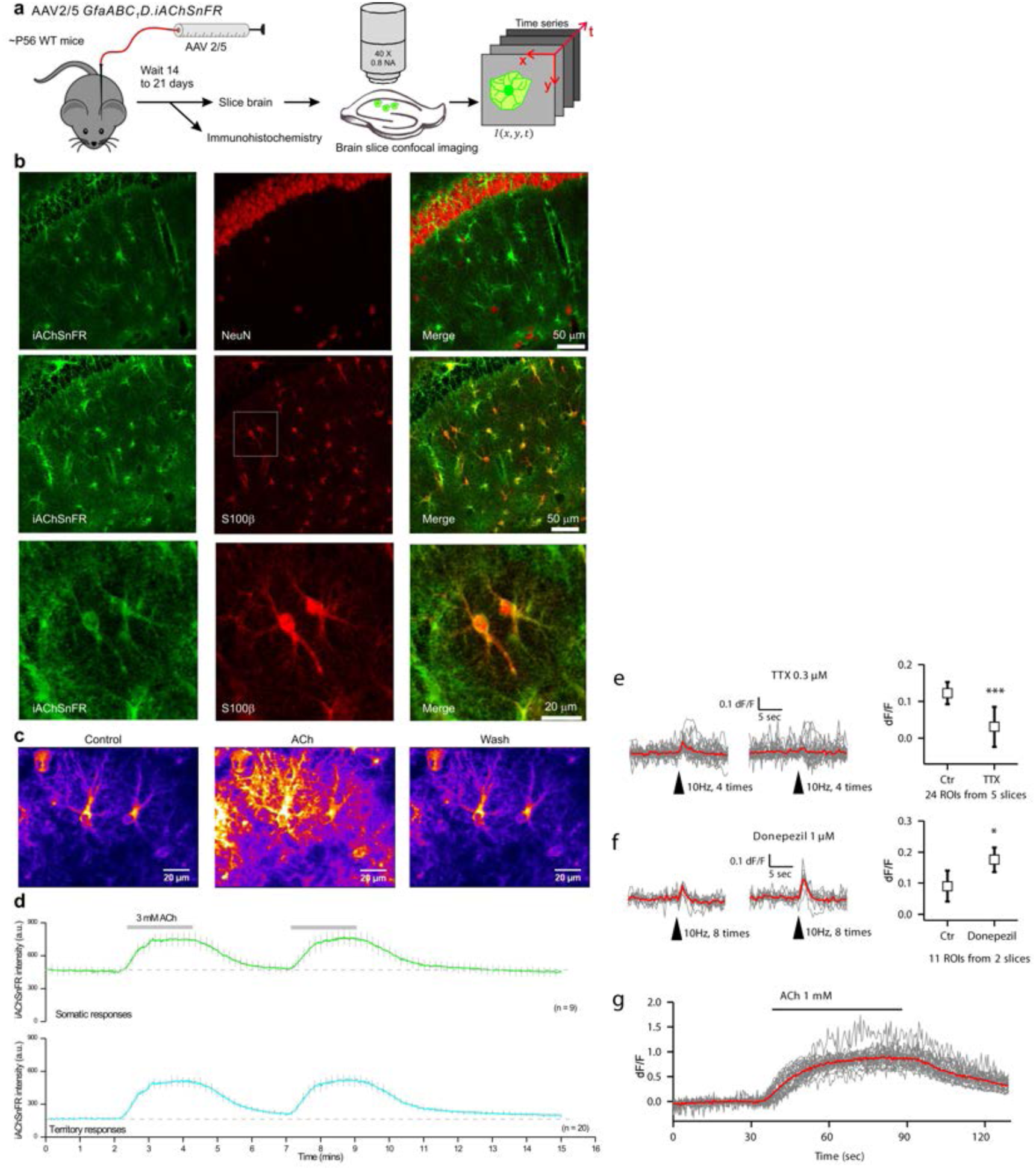
Hippocampal astrocyte expression and responses. (**a**) Schematic illustrates the protocol of astrocyte cell-surface ACh imaging using iAChSnFR. (**b**) Immunohistochemical analysis shows astrocytic expression of iAChSnFR. iAChSnFR was expressed in S100β-positive cells (astrocytes) but not NeuN-positive cells (neurons). (**c**) Representative images of iAChSnFR before (control), during (ACh) and after (wash) application of ACh. The images are shown on an arbitrary “fire” color scale from 0 (black) to 256 (white), with warmer colors indicating higher fluorescence. (**d**) Averaged traces of iAChSnFR signals from soma and processes. (**e**) Electrically evoked ACh increases on striatal astrocytes. Grey traces indicate individual traces. Red traces indicate averaged traces. Evoked signals were abolished by TTX (control, 0.12 ± 0.01; TTX, 0.03 ± 0.03, *t*(23) = 3.93, *p* < 0.001, n = 24, paired Student’s *t*-test). (**f**) Evoked ACh increases were enhanced by donepezil, an AChE inhibitor (control, 0.09 ± 0.02; donepezil, 0.18 ± 0.02, *t*(11) = 2.35, *p*<0.05, n = 12, paired Student’s *t*-test). (**g**) Bath application of ACh induced robust iAChSnFR signals on striatal astrocytes.

**Supplementary Fig. S18.**
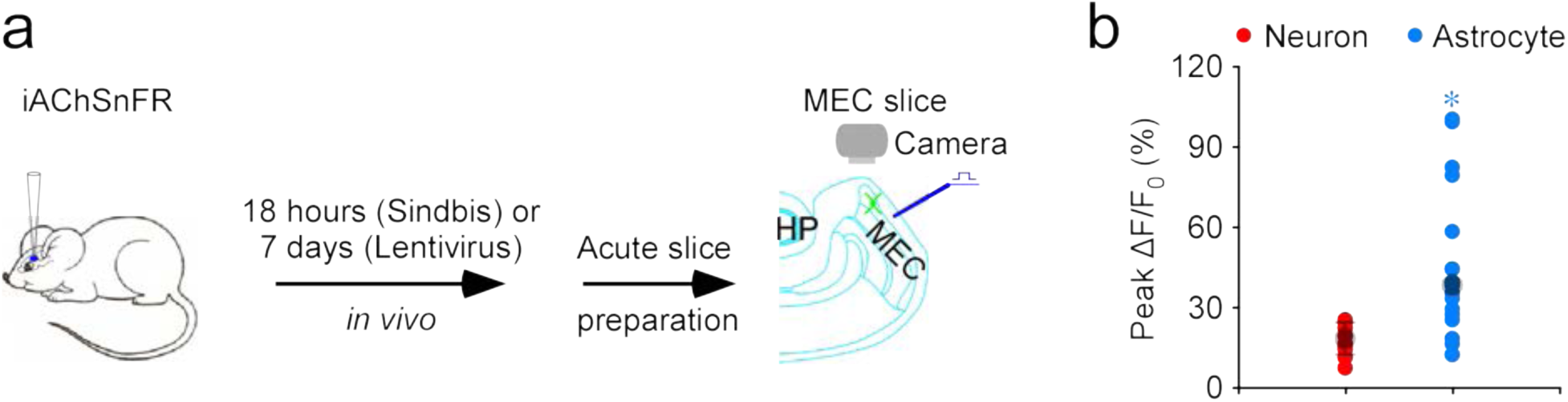
iAChSnFR detects especially large signals at single release sites near entorhinal astrocytes. (a) Schematic drawing of stimulation-imaging experiments on entorhinal neurons and astrocytes in acute mouse MEC slices. HP: hippocampus. (b) Values for peak ΔF/F_0_ responses at single release sites of entorhinal neurons (e.g., see Fig. 2j) and astrocytes (e.g., see Fig. 2o) (Neuron: 17.6±1.5%, *n*=13 from 6 neurons from 6 animals; Astrocyte: 38.4±5.9%, *n*=22 from 10 neurons from 8 animals; *U*=216.0, *p*=0.013). Large gray dots indicate average responses and asterisk indicates *p*<0.05 (Mann-Whitney rank-sum non-parametric test).

**Supplementary Fig. S19.**
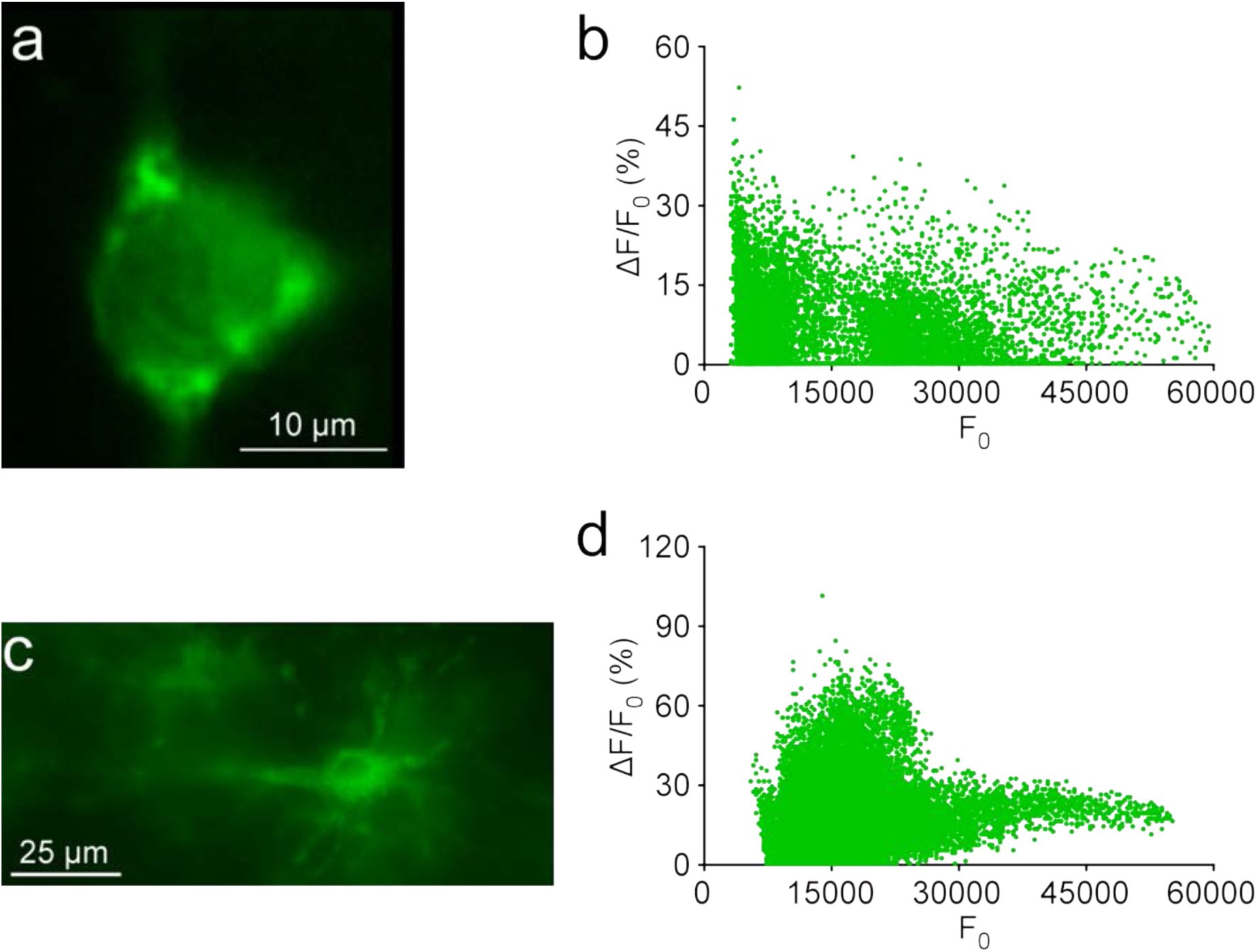
iAChSnFR provides useful data over a wide range of expression levels. (**a**) Analysis on the sub-µm scale. A snapshot of the iAChSnFR-expressing MEC stellate neuron shown in Figure 2h-k (image is 123 x 166 pixels, 0.181 µm/pixel). (**b**) Scatter plot of ΔF/F_0_ against F_0_ of the neuron in (a) at each pixel (20,418 pixels in the ROI). The data show a weak negative correlation between ΔF/F_0_ and F_0_ with a slope value of −0.00014 (*n* = 20,418; normality test *p =* 0.001; constant variance test *p =* 0.001; *r^2^* = 0.047; *F* = 1,004.46; *p <* 0.001; linear regression *t* test). (**c**) Analysis on the µm scale. A snapshot of the iAChSnFR-expressing MEC astrocyte shown in Figure 2m-p (image is 694 x 331 pixels, 0.181 µm/pixel). (**d**) Scatter plot of ΔF/F_0_ against F_0_ of the astrocyte in (c) at each pixel (229,714 pixels in the ROI). a weak negative correlation between ΔF/F_0_ and F_0_ with a slope value of −0.000703 (*n* = 65,536; normality test *p <* 0.001; Constant variance test *p* < 0.001; *r^2^* = 0.0507; *F* = 12,257.27; *p <* 0.001; linear regression *t*-test).

**Supplementary Fig. S20.**
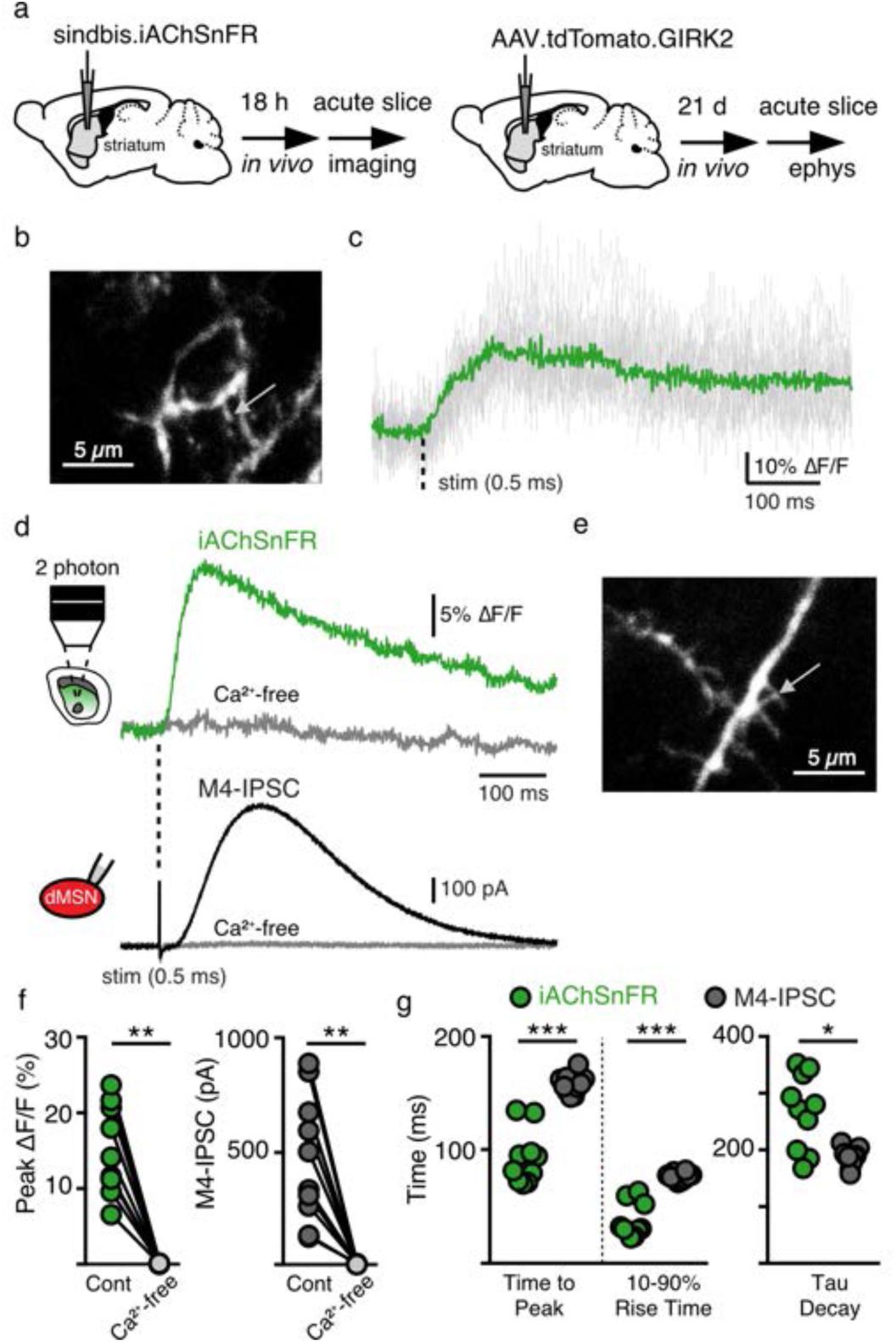
iAChSnFR signals precede ACh-mediated potassium currents in the striatum. (**a**) Schematic illustration of the procedure. (**b**) 2-photon iAChSnFR fluorescence image in a slice from dorsal striatum. Arrow shows location of iAChSnFR hotspot illustrated in panel **c**. (**c**) 2-photon excitation and spot photometry fluorescence waveform for electrical stimulation (30 μA; 0.5 ms). Green trace is the average of 13 sweeps. Single trial fluorescence responses in gray. (**d**) 2-photon excitation and spot photometry fluorescence waveform (average 40 sweeps), compared with a presumptive muscarinic acetylcholine receptor M_4_-mediated inhibitory post-synaptic current (IPSC) from a dorsal medium spiny neuron (dMSN; average 5 sweeps). (**e**) 2-photon fluorescence image of iAChSnFR in the dorsal striatum. Arrow indicates location of iAChSnFR hotspot illustrated in panel **d**. (**f**) Peak amplitude of fluorescence waveforms (ΔF/F %) and M_4_-IPSC amplitude (pA) under control conditions and in the presence of TTX (1 μM) and Ca^2+^-free ACSF (*n* = 10). **: p <0.01, Wilcoxon matched-pairs signed-rank test. (***g***) Summary of fluorescence waveforms and M_4_-IPSC waveforms. Activation kinetics for iAChSnFR (10-90% rise time = 37.2 ± 4.8 ms, time to peak = 92.9 ± 7.5 ms; M_4_-mediated IPSCs: 10-90% rise time = 76.3 ± 1.1 ms. Time-to-peak = 157.5 ± 2.7 ms; *p* < 0.001 for both; *n* = 10. Decay kinetics (iAChSnFR: tau = 258 ± 26.4 ms; M_4_-mediated IPSC: tau = 186.1 ± 5.4 ms, *p* < 0.05, *n* = 10) of fluorescence responses and M_4_-IPSCs. * = p < 0.05; ** = p < 0.01, *** = p < 0.001, Mann-Whitney test.

**Supplementary Figure S21.**
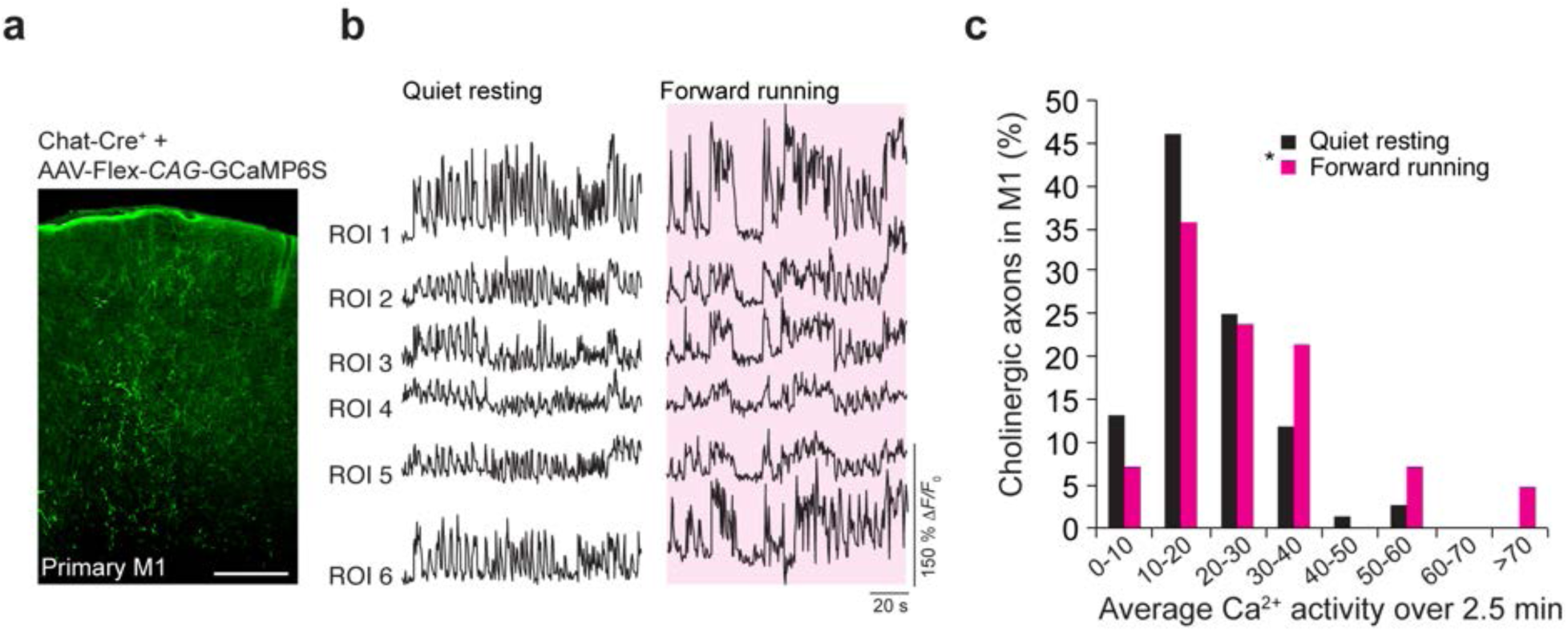
Active cholinergic projections in mouse forelimb motor cortex. (**a**) Coronal section of mouse primary motor cortex from a *ChAT*-Cre mouse infected with AAV2/1-Flex-*CAG*-GCaMP6s in nucleus basalis following 2 weeks of expression. Scale bar, 100 µm. (**b**) Representative fluorescent traces from different axonal segments expressing GCaMP6s (labeled ROIs 1-6) in the motor cortex during quiet resting (left) and forward running (right, magenta shaded). (**c**) Distribution of cholinergic axon Ca^2+^ activity during quiet resting and forward running. \**P* < 0.05, paired Student’s *t*-test.

**Supplementary Fig. S22.**
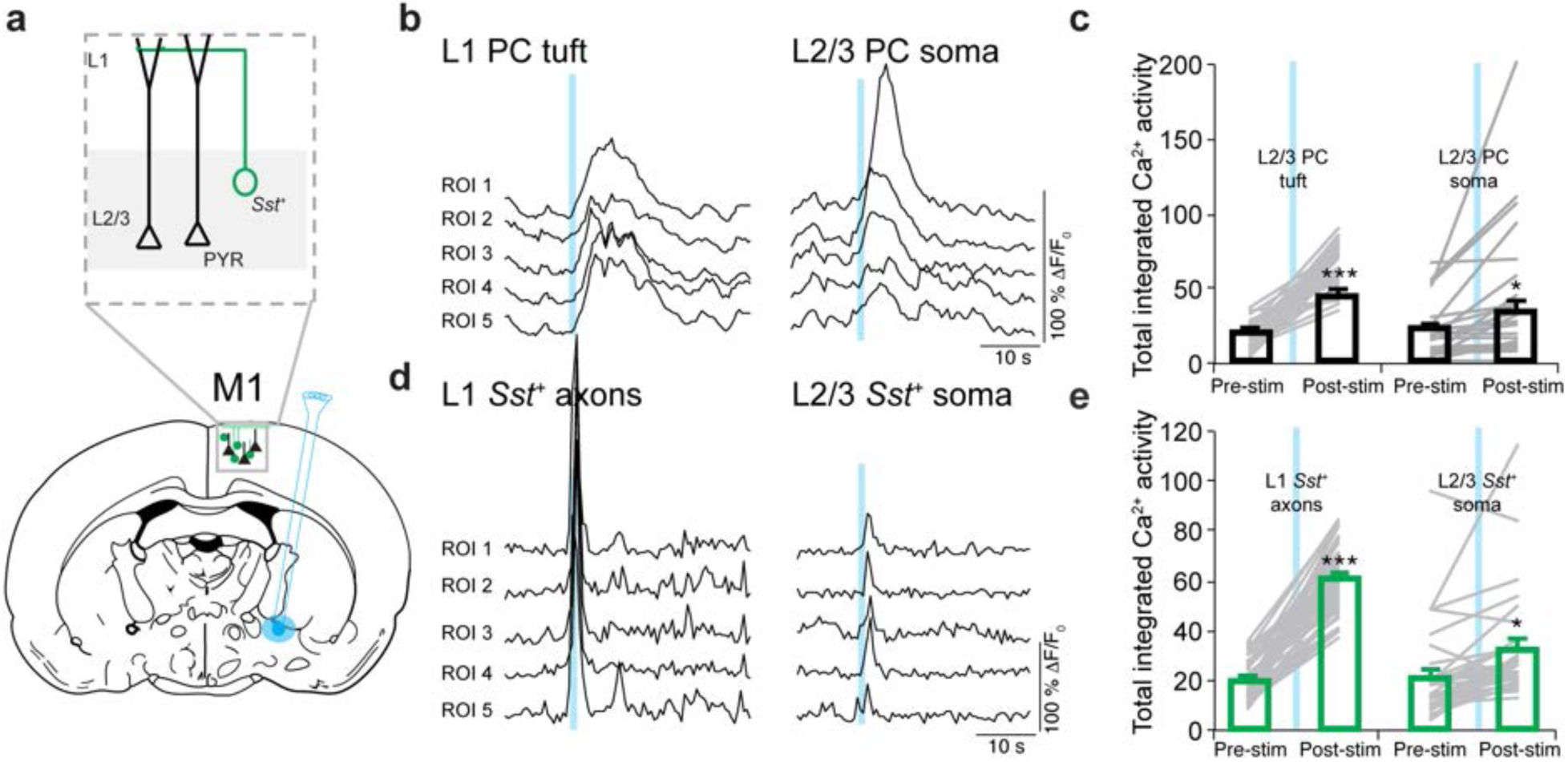
Cortical neuronal activities of layer 2/3 pyramidal cells and *Sst^+^* cells in response to evoked nucleus basalis (NB) activation. (**a**) Schematic of experimental design with stimulating bipolar electrode placed in nucleus basalis and GCaMP6s expression in L2/3 neurons. GCaMP6s was selectively expressed in *Sst^+^* neurons or in PCs (separate animals). Two-photon imaging was performed at the level of PC dendrites (L1) and somata (L2/3) and *Sst^+^* axon fibers (L1) and somata (L2/3) (outlined in dashed box). (**b**) Fluorescence traces from representative L2/3 PC dendritic segments (left) and somata (right) expressing GCaMP6s before and after NB stimulation. (**c**) Average total integrated Ca^2+^ activity detected with GCaMP6s over 10 sec before and after NB stimulation for apical tuft dendrites and L2/3 somata. Apical tuft dendrites (*P* < 0.001, *n* = 37) and somata (*P* < 0.05, *n* = 24) of L2/3 neurons displayed significant enhancement of calcium activity following NB stimulation. (**d**) Fluorescence traces from representative L1 *Sst^+^* axonal segments (left) and L2/3 somata (right) expressing GCaMP6s before and after NB stimulation. (**e**) Average total integrated Ca^2+^ activity detected with GCaMP6s before and after NB stimulation for L1 *Sst^+^* axonal segments and L2/3 *Sst^+^* somata. L1 axonal segments (*P* < 0.001, *n* = 60) and L2/3 *Sst^+^* somata (*P* < 0.05, *n* = 36) displayed significant enhancement of calcium activity following NB stimulation. The peak response of *Sst^+^* cell activation occurred 5.5 sec before the peak response of PCs. Data are presented as means ± s.e.m. \**P* < 0.05, \*\*\**P* < 0.001, paired Student’s *t*-test.

**Supplementary Fig. S23.**
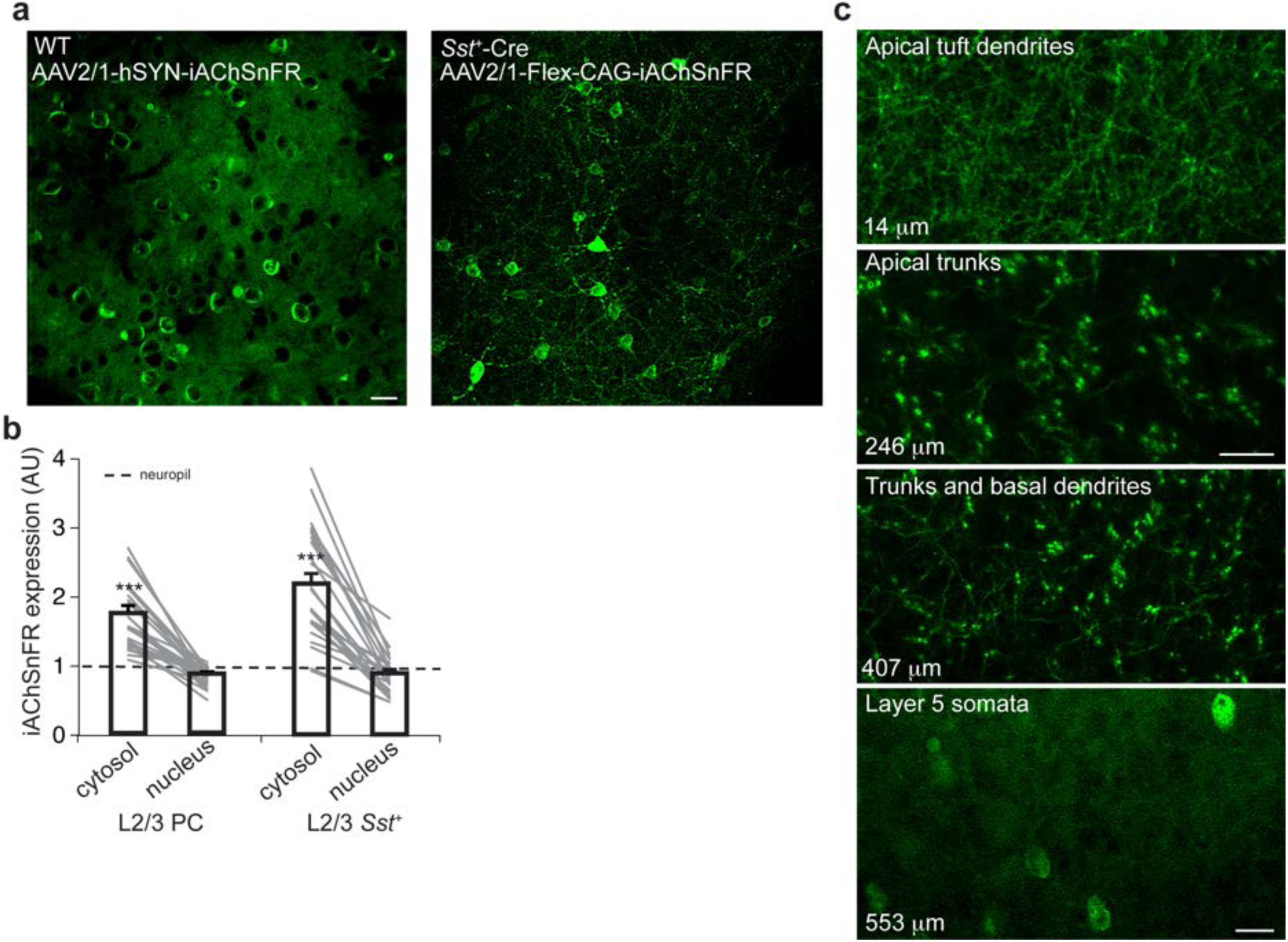
iAChSnFR expression in primary motor cortex. (**a**) Two-photon images of L2/3 neurons expressing iAChSnFR in wild-type (WT; left image) and *Sst-*Cre mice (right image). Scale bar, 20 µm. (**b**) Fluorescence measurements across somata of L2/3 PC (*n* = 23) and *Sst^+^* cells (*n* = 24). Nuclear signals were comparable or less than neuropil signals, demonstrating significant expression in the cytosol. Data are presented as means ± s.e.m. \*\*\**P* < 0.001, unpaired Student’s *t*-test. (**c**) Two-photon images across different cortical depths (thin branches of apical tuft, apical trunk, and deep basal dendrites; depth indicated) of L5 neurons expressing iAChSnFR. To enable L5 expression, AAV2/1-Flex*-CAG*-iAChSnFR was injected into *Rbp4*-Cre mice. Scale bars, 20 µm.

**Supplementary Fig. S24.**
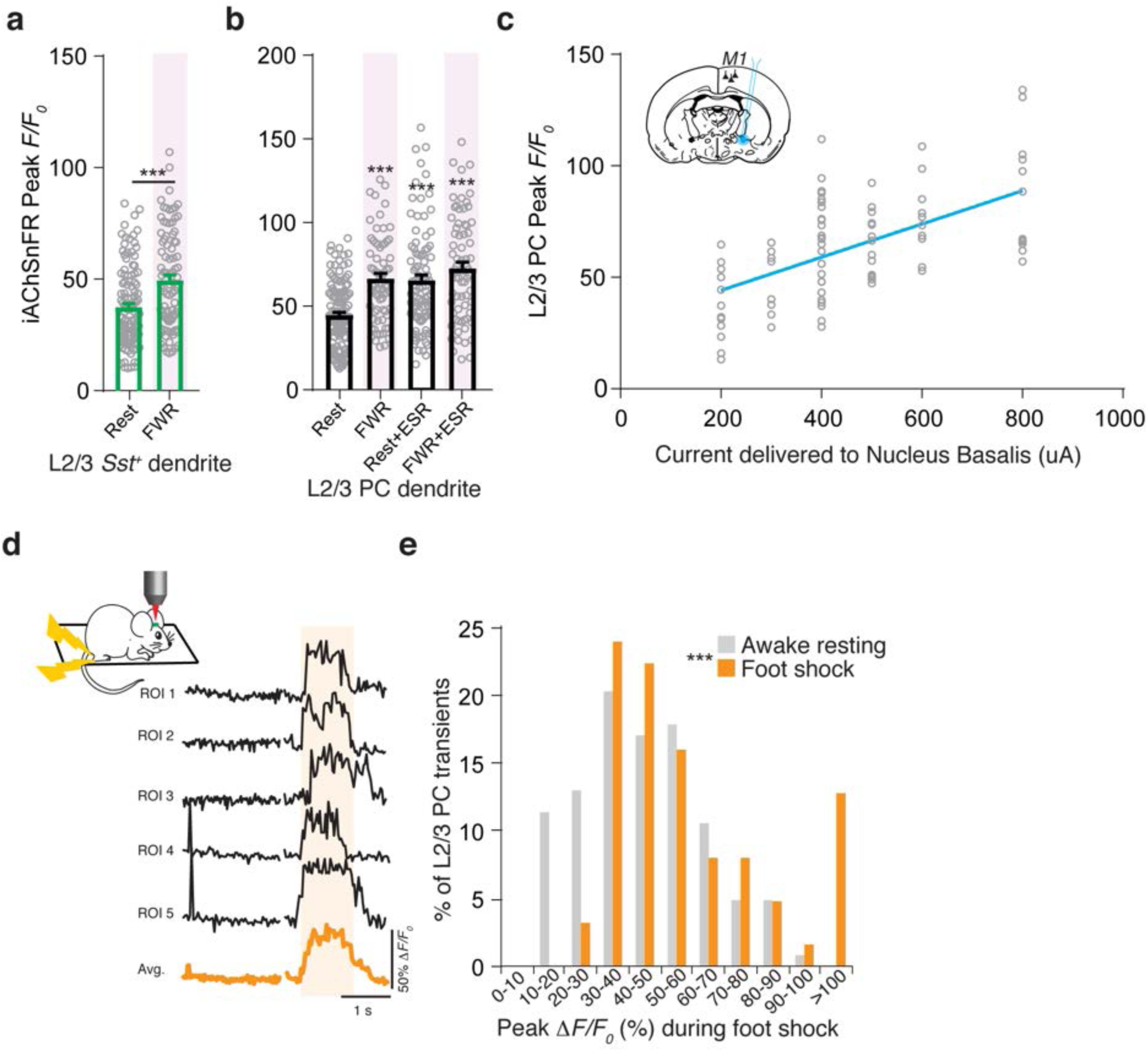
Responses and modulation of iAChSnFR in *Sst^+^* interneurons and pyramidal neurons of mouse motor cortex. (**a**) Average peak iAChSnFR signals from *Sst^+^* dendrites (rest: 38 ± 2%, *n* =112; run: 50 ± 2%, *n* = 89; *P* < 0.001, t=4.293, df=199). (**b**) Average peak iAChSnFR signals from PC dendrites during quiet resting and forward running (rest: 45 ± 2%, *n* = 124; run: 66 ± 3%, *n* = 69, *P* < 0.001, t=5.48, df=353), and in the presence of local eserine (rest: 65 ± 3%, *n* = 93, *P* < 0.001; t=5.71, df=353, run: 73 ± 4%, *n* = 71, *P* < 0.001, t=7.33, df=353). (**c**) Simultaneous activation of NB and 2-photon recording of iAChSnFR signals in L2/3 PC dendrites. Note that peak iAChSnFR signals are proportional to the amount of current delivered to NB (R^2^=0.33, *P* < 0.001). (**d**) Representative fluorescence traces before and during 1 sec foot shock to hindlimb. (**e**) Distribution of peak iAChSnFR signals from dendrites of L2/3 PCs during foot shock as compared to no foot shock (no shock: 45 ± 2%, *n* = 124, shock: 59 ± 4%, *n* = 63, *P* < 0.001). Data are presented as means ± s.e.m. ***P < 0.001, unpaired Student’s *t*-test in **a**, **e**, and two-way ANOVA followed by Bonferroni’s test in **b**.

**Supplementary Fig. S25.**
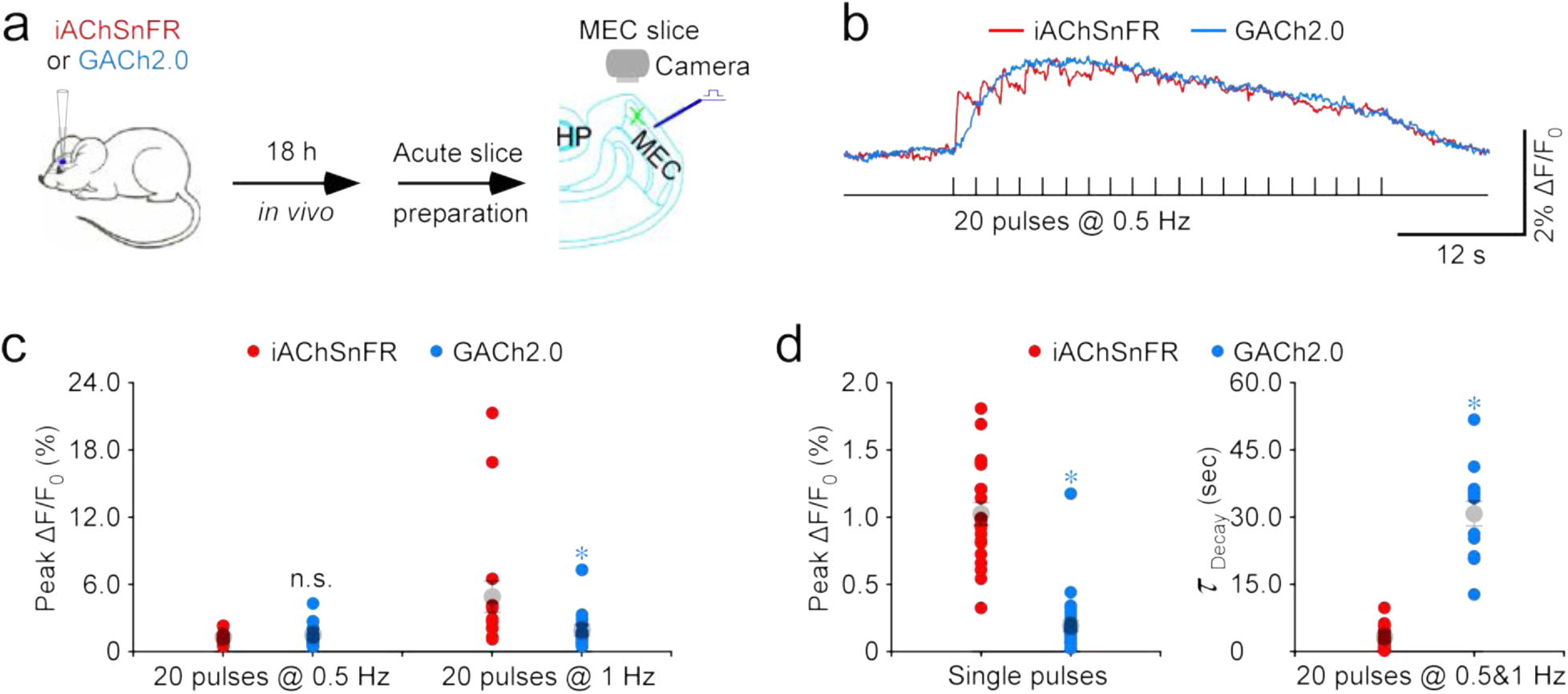
iAChSnFR performs better than GACh2.0 in detecting cholinergic signals. (**a**) Schematic of stimulation-imaging experiments in acute mouse entorhinal cortical (MEC) slices. (**b**) Representative fluorescence ΔF/F_0_ waveforms of iAChSnFR and GACh2.0 expressing MEC neurons to trains of 20 electrical pulses at 0.5 Hz. (**c**) Values for peak ΔF/F_0_ responses evoked by trains of 20 electrical pulses at 0.5 Hz (iAChSnFR: 1.2±0.2%, *n*=15 from 14 animals; GACh2.0: 1.4±0.3%, *n*=14 from 8 animals; *U*=118.0, *p*=0.585) and 1 Hz (iAChSnFR: 4.8±1.4%, *n*=16 from 15 animals; GACh2.0: 1.8±0.5%, *n*=14 from 8 animals; *U*=181.0, *p*=0.004) in iAChSnFR and GACh2.0 expressing entorhinal stellate neurons. (**d**) Left, values for peak ΔF/F_0_ responses evoked by single electrical pulses in iAChSnFR and GACh2.0 expressing entorhinal stellate neurons (iAChSnFR: 1.0±0.1%, *n*=21 from 13 animals; GACh2.0: 0.2±0.04%, *n*=28 from 7 animals; *U*=0.0, *p*<0.001). Right, values for decay time constants of fluorescence waveforms evoked by trains of 20 electrical pulses at 0.5 Hz and 1 Hz of iAChSnFR and GACh2.0 expressing entorhinal stellate neurons (iAChSnFR: 2.9±0.4 sec, *n*=32 from 12 animals; GACh2.0: 30.6±2.7 sec, *n*=14 from 7 animals; *U*=0.0, *p*<0.001). Large gray dots indicate average responses and asterisks indicate *p*<0.05 (Mann-Whitney rank-sum non-parametric tests).

## Supplementary Movie Legends

Movie S1. Imaging responses of a hippocampal CA1 pyramidal neuron to ACh puff application.

Movie S2. Imaging responses of the same CA1 pyramidal neuron to nicotine puff application.

Movie S3. Imaging endogenous cholinergic transmission at an entorhinal stellate neuron.

Movie S4. Imaging endogenous ACh release at individual release sites of an entorhinal stellate neuron.

Movie S5. Imaging endogenous ACh release at an entorhinal astrocyte.

## References

1. Engelhardt von, J., Eliava, M., Meyer, A. H., Rozov, A. & Monyer, H. Functional characterization of intrinsic cholinergic interneurons in the cortex. J. Neurosci. 27, 5633–5642 (2007).

2. Yi, F. et al. Hippocampal ‘cholinergic interneurons’ visualized with the choline acetyltransferase promoter: anatomical distribution, intrinsic membrane properties, neurochemical characteristics, and capacity for cholinergic modulation. Front. Synaptic. Neurosci. 7, 4 (2015).

3. Himmelheber, A. M., Sarter, M. & Bruno, J. P. Increases in cortical acetylcholine release during sustained attention performance in rats. Brain Res. Cogn. Brain. Res. 9, 313–325 (2000).

4. Gadea-Ciria, M., Stadler, H., Lloyd, K. G. & Bartholini, G. Acetylcholine release within the cat striatum during the sleep-wakefulness cycle. Nature 243, 518–519 (1973).

5. Hasselmo, M. E. The role of acetylcholine in learning and memory. Curr. Op. Neurobiology 16, 710–715 (2006).

6. Rosas-Ballina, M. & Tracey, K. J. Cholinergic control of inflammation. J. Intern. Med. 265, 663–679 (2009).

7. Granger, A. J., Mulder, N., Saunders, A. & Sabatini, B. L. Cotransmission of acetylcholine and GABA. Neuropharmacol. 100, 40–46 (2016).

8. Hnasko, T. S. & Edwards, R. H. Neurotransmitter corelease: mechanism and physiological role. Annu. Rev. Physiol. 74, 225–243 (2012).

9. Millar, N. S. & Gotti, C. Diversity of vertebrate nicotinic acetylcholine receptors. Neuropharmacol. 56, 237–246 (2009).

10. Picciotto, M. R., Higley, M. J. & Mineur, Y. S. Acetylcholine as a neuromodulator: cholinergic signaling shapes nervous system function and behavior. Neuron 76, 116–129 (2012).

11. Soreq, H. & Seidman, S. Acetylcholinesterase--new roles for an old actor. Nat. Rev. Neurosci. 2, 294–302 (2001).

12. VanPatten, S. & Al-Abed, Y. The challenges of modulating the ‘rest and digest’ system: acetylcholine receptors as drug targets. Drug Discovery Today 22, 97–104 (2017).

13. Tansey, E. M. Henry Dale and the discovery of acetylcholine. Comptes rendus biologies 329, 419–425 (2006).

14. Figl, A., Labarca, C., Davidson, N., Lester, H. A. & Cohen, B. N. Voltage-jump relaxation kinetics for wild-type and chimeric beta subunits of neuronal nicotinic receptors. J. Gen. Physiol. 107, 369–379 (1996).

15. Jenden, D. J., Hanin, I. & Lamb, S. I. Gas chromatographic microestimation of acetylcholine and related compounds. Anal. Chem. 40, 125–128 (1968).

16. Parikh, V. & Sarter, M. Cortical choline transporter function measured *in vivo* using choline-sensitive microelectrodes: clearance of endogenous and exogenous choline and effects of removal of cholinergic terminals. J. Neurochem. 97, 488–503 (2006).

17. Ziegler, N., Bätz, J., Zabel, U., Lohse, M. J. & Hoffmann, C. FRET-based sensors for the human M1-, M3-, and M5-acetylcholine receptors. Bioorg. Med. Chem. 19, 1048–1054 (2011).

18. Schena, A. & Johnsson, K. Sensing acetylcholine and anticholinesterase compounds. Angew. Chem. Int. Ed. Engl. 53, 1302–1305 (2014).

19. Nguyen, Q.-T. et al. An *in vivo* biosensor for neurotransmitter release and *in situ* receptor activity. Nat. Neurosci. 13, 127–132 (2010).

20. Jing, M. et al. A genetically encoded fluorescent acetylcholine indicator for *in vitro* and *in vivo* studies. Nat. Biotechnol. 50, 279 (2018).

21. Tubio, M. R. et al. Expression of a G protein-coupled receptor (GPCR) leads to attenuation of signaling by other GPCRs: experimental evidence for a spontaneous GPCR constitutive inactive form. J. Biol. Chem. 285, 14990–14998 (2010).

22. Wang, L. et al. Overexpression of G protein-coupled receptor GPR87 promotes pancreatic cancer aggressiveness and activates NF-κB signaling pathway. Mol. Cancer 16, 61 (2017).

23. Gargiulo, L. et al. A Novel Effect of β-Adrenergic Receptor on Mammary Branching Morphogenesis and its Possible Implications in Breast Cancer. J Mammary Gland Biol Neoplasia 22, 43–57 (2017).

24. Marvin, J. S. et al. An optimized fluorescent probe for visualizing glutamate neurotransmission. Nat. Methods 10, 162–170 (2013).

25. Marvin, J. S. et al. Stability, affinity, and chromatic variants of the glutamate sensor iGluSnFR. Nat. Methods 15, 936–939 (2018).

26. Marvin, J. S. et al. A genetically encoded fluorescent sensor for *in vivo* imaging of GABA. Nat. Methods 16, 763–770 (2019).

27. Keller, J. P. et al. *In vivo* glucose imaging in multiple model organisms with an engineered single-wavelength sensor. bioRxiv, doi:doi/10.1101/571422 (2019).

28. Díaz-García, C. M. et al. Quantitative *in vivo* imaging of neuronal glucose concentrations with a genetically encoded fluorescence lifetime sensor. J. Neurosci. Res. 97, 946–960 (2019).

29. Marvin, J. S., Schreiter, E. R., Echevarría, I. M. & Looger, L. L. A genetically encoded, high-signal-to-noise maltose sensor. Proteins 79, 3025–3036 (2011).

30. Alicea, I. et al. Structure of the *Escherichia coli* phosphonate binding protein PhnD and rationally optimized phosphonate biosensors. J. Mol. Biol. 414, 356–369 (2011).

31. Oswald, C., Smits, S. H. J., Höing, M., Bremer, E. & Schmitt, L. Structural analysis of the choline-binding protein ChoX in a semi-closed and ligand-free conformation. Biol. Chem. 390, 1163–1170 (2009).

32. Pittelkow, M., Tschapek, B., Smits, S. H. J., Schmitt, L. & Bremer, E. The crystal structure of the substrate-binding protein OpuBC from *Bacillus subtilis* in complex with choline. J. Mol. Biol. 411, 53–67 (2011).

33. Pédelacq, J.-D., Cabantous, S., Tran, T., Terwilliger, T. C. & Waldo, G. S. Engineering and characterization of a superfolder green fluorescent protein. Nat. Biotechnol. 24, 79–88 (2006).

34. Du, Y. et al. Structures of the substrate-binding protein provide insights into the multiple compatible solute binding specificities of the *Bacillus subtilis* ABC transporter OpuC. Biochem. J. 436, 283–289 (2011).

35. Marvin, J. S. & Hellinga, H. W. Manipulation of ligand binding affinity by exploitation of conformational coupling. Nat. Struct. Biol. 8, 795–798 (2001).

36. Shivange, A. V. et al. Determining the pharmacokinetics of nicotinic drugs in the endoplasmic reticulum using biosensors. J. Gen. Physiol. 151, 738–757 (2019).

37. Nagai, T. et al. A variant of yellow fluorescent protein with fast and efficient maturation for cell-biological applications. Nat. Biotechnol. 20, 87–90 (2002).

38. Kazemipour, A. et al. Kilohertz frame-rate two-photon tomography. Nat. Methods 16, 778–786 (2019).

39. Jakubík, J., Bačáková, L., El-Fakahany, E. E. & Tuček, S. Positive cooperativity of acetylcholine and other agonists with allosteric ligands on muscarinic acetylcholine receptors. Molec. Pharmacol. 52, 172 (1997).

40. Shen, J. & Barnes, C. A. Age-related decrease in cholinergic synaptic transmission in three hippocampal subfields. Neurobiol. Aging 17, 439–451 (1996).

41. Dani, J. A. & Bertrand, D. Nicotinic acetylcholine receptors and nicotinic cholinergic mechanisms of the central nervous system. Annu. Rev. Pharmacol. Toxicol. 47, 699–729 (2007).

42. Mamaligas, A. A. & Ford, C. P. Spontaneous Synaptic Activation of Muscarinic Receptors by Striatal Cholinergic Neuron Firing. Neuron 91, 574–586 (2016).

43. Simon, A. P., Poindessous-Jazat, F., Dutar, P., Epelbaum, J. & Bassant, M.-H. Firing properties of anatomically identified neurons in the medial septum of anesthetized and unanesthetized restrained rats. J. Neurosci. 26, 9038–9046 (2006).

44. Duque, A., Tepper, J. M., Detari, L., Ascoli, G. A. & Zaborszky, L. Morphological characterization of electrophysiologically and immunohistochemically identified basal forebrain cholinergic and neuropeptide Y-containing neurons. Brain Struct. Funct. 212, 55–73 (2007).

45. Zhu, P. K. et al. Nanoscopic visualization of restricted non-volume cholinergic and monoaminergic transmissions. Nano Letters in press, 999 (2020).

46. Duffy, A. M. et al. Acetylcholine α7 nicotinic and dopamine D2 receptors are targeted to many of the same postsynaptic dendrites and astrocytes in the rodent prefrontal cortex. Synapse 65, 1350–1367 (2011).

47. Zhang, L. et al. Ras and Rap Signal Bidirectional Synaptic Plasticity via Distinct Subcellular Microdomains. Neuron 98, 783–800.e4 (2018).

48. Hersch, S. M., Gutekunst, C. A., Rees, H. D., Heilman, C. J. & Levey, A. I. Distribution of m1-m4 muscarinic receptor proteins in the rat striatum: light and electron microscopic immunocytochemistry using subtype-specific antibodies. J. Neurosci. 14, 3351–3363 (1994).

49. Cai, Y. & Ford, C. P. Dopamine Cells Differentially Regulate Striatal Cholinergic Transmission across Regions through Corelease of Dopamine and Glutamate. Cell Rep. 25, 3148–3157.e3 (2018).

50. Richmond, J. E. & Jorgensen, E. M. One GABA and two acetylcholine receptors function at the C. elegans neuromuscular junction. Nat. Neurosci. 2, 791–797 (1999).

51. Collins, K. M. et al. Activity of the *C. elegans* egg-laying behavior circuit is controlled by competing activation and feedback inhibition. eLife 5, 13819 (2016).

52. Korn, H. & Faber, D. S. The Mauthner cell half a century later: a neurobiological model for decision-making? Neuron 47, 13–28 (2005).

53. Liu, K. S. & Fetcho, J. R. Laser ablations reveal functional relationships of segmental hindbrain neurons in zebrafish. Neuron 23, 325–335 (1999).

54. Pujala, A. & Koyama, M. Chronology-based architecture of descending circuits that underlie the development of locomotor repertoire after birth. eLife 8, 471 (2019).

55. Fetcho, J. R. & Faber, D. S. Identification of motoneurons and interneurons in the spinal network for escapes initiated by the mauthner cell in goldfish. J. Neurosci. 8, 4192–4213 (1988).

56. Zheng, Z. et al. A Complete Electron Microscopy Volume of the Brain of Adult *Drosophila melanogaster*. Cell 174, 730–743.e22 (2018).

57. Stocker, R. F., Heimbeck, G., Gendre, N. & de Belle, J. S. Neuroblast ablation in Drosophila P[GAL4] lines reveals origins of olfactory interneurons. J. Neurobiol. 32, 443–456 (1997).

58. Laissue, P. P. et al. Three-dimensional reconstruction of the antennal lobe in *Drosophila melanogaster*. J. Comp. Neurol. 405, 543–552 (1999).

59. Kirkpatrick, R. B. et al. Heavy chain dimers as well as complete antibodies are efficiently formed and secreted from *Drosophila* via a BiP-mediated pathway. J. Biol. Chem. 270, 19800–19805 (1995).

60. Laurent, G. Olfactory network dynamics and the coding of multidimensional signals. Nat. Rev. Neurosci. 3, 884–895 (2002).

61. Wang, J. W., Wong, A. M., Flores, J., Vosshall, L. B. & Axel, R. Two-photon calcium imaging reveals an odor-evoked map of activity in the fly brain. Cell 112, 271–282 (2003).

62. Bhandawat, V., Olsen, S. R., Gouwens, N. W., Schlief, M. L. & Wilson, R. I. Sensory processing in the *Drosophila* antennal lobe increases reliability and separability of ensemble odor representations. Nat. Neurosci. 10, 1474–1482 (2007).

63. Stopfer, M., Jayaraman, V. & Laurent, G. Intensity versus identity coding in an olfactory system. Neuron 39, 991–1004 (2003).

64. Hallem, E. A. & Carlson, J. R. Coding of odors by a receptor repertoire. Cell 125, 143–160 (2006).

65. McCormick, D. A. & Prince, D. A. Two types of muscarinic response to acetylcholine in mammalian cortical neurons. Proc. Natl. Acad. Sci. U.S.A. 82, 6344–6348 (1985).

66. McCormick, D. A. & Prince, D. A. Mechanisms of action of acetylcholine in the guinea-pig cerebral cortex *in vitro*. J. Physiol. 375, 169–194 (1986).

67. Kawaguchi, Y. Selective cholinergic modulation of cortical GABAergic cell subtypes. J. Neurophysiol. 78, 1743–1747 (1997).

68. Arroyo, S., Bennett, C., Aziz, D., Brown, S. P. & Hestrin, S. Prolonged disynaptic inhibition in the cortex mediated by slow, non-α7 nicotinic excitation of a specific subset of cortical interneurons. J. Neurosci. 32, 3859–3864 (2012).

69. Delmas, P. & Brown, D. A. Pathways modulating neural KCNQ/M (Kv7) potassium channels. Nat. Rev. Neurosci. 6, 850–862 (2005).

70. Wall, N. R. et al. Brain-Wide Maps of Synaptic Input to Cortical Interneurons. J. Neurosci. 36, 4000–4009 (2016).

71. Chen, T.-W. et al. Ultrasensitive fluorescent proteins for imaging neuronal activity. Nature 499, 295–300 (2013).

72. Harris, J. A. et al. Anatomical characterization of Cre driver mice for neural circuit mapping and manipulation. Front Neural Circuits 8, 76 (2014).

73. Blokland, A. Acetylcholine: a neurotransmitter for learning and memory? Brain Res. Brain Res. Rev. 21, 285–300 (1995).

74. Mattinson, C. E. et al. Tonic and phasic release of glutamate and acetylcholine neurotransmission in sub-regions of the rat prefrontal cortex using enzyme-based microelectrode arrays. Journal of Neuroscience Methods 202, 199–208 (2011).

75. Bloom, F. E., Costa, E. & Salmoiraghi, G. C. Anesthesia and the responsiveness of individual neurons of the caudate nucleus of the cat to acetylcholine, norepinephrine and dopamine administered by microelectrophoresis. J. Pharmacol. Exp. Ther. 150, 244 (1965).

76. Shichino, T. et al. Effects of inhalation anaesthetics on the release of acetylcholine in the rat cerebral cortex *in vivo*. Br. J. Anaesth. 80, 365–370 (1998).

77. Kristt, D. A. in Cerebral Cortex: Further Aspects of Cortical Function, Including Hippocampus (eds. Jones, E. G. & Peters, A.) 187–236 (Springer US, 1987).

78. Orsetti, M., Casamenti, F. & Pepeu, G. Enhanced acetylcholine release in the hippocampus and cortex during acquisition of an operant behavior. Brain Research 724, 89–96 (1996).

79. Teles-Grilo Ruivo, L. M. et al. Coordinated acetylcholine release in prefrontal cortex and hippocampus is associated with arousal and reward on distinct timescales. Cell Rep. 18, 905–917 (2017).

80. Nyakas, C., Luiten, P. G., Spencer, D. G. & Traber, J. Detailed projection patterns of septal and diagonal band efferents to the hippocampus in the rat with emphasis on innervation of CA1 and dentate gyrus. Brain Res. Bull. 18, 533–545 (1987).

81. Jing, M. et al. An optimized acetylcholine sensor for monitoring in vivo cholinergic activity. bioRxiv https://doi.org/10.1101/861690 (2019).

82. Bao, H. et al. Exocytotic fusion pores are composed of both lipids and proteins. Nat. Struct. Biol. 23, 67–73 (2016).

83. Keller, J. P. & Looger, L. L. The Oscillating Stimulus Transporter Assay, OSTA: Quantitative Functional Imaging of Transporter Protein Activity in Time and Frequency Domains. Molec. Cell 64, 199–212 (2016).

84. McGirr, A., LeDue, J., Chan, A. W., Xie, Y. & Murphy, T. H. Cortical functional hyperconnectivity in a mouse model of depression and selective network effects of ketamine. Brain 140, 2210–2225 (2017).

85. Cui, Q. et al. Blunted mGluR activation disinhibits striatopallidal transmission in Parkinsonian mice. Cell Rep. 17, 2431–2444 (2016).

86. Unger, E. K. et al. Directed evolution of a selective and sensitive serotonin biosensor *via* machine learning. Cell doi:dx.doi.org/10.2139/ssrn.3498571

87. Bera, K. et al. Biosensors Show the Pharmacokinetics of S-Ketamine in the Endoplasmic Reticulum. Front. Cell. Neurosci. 13, 499 (2019).

88. Muthusamy, A. K. et al. Microscopy Using Fluorescent Drug Biosensors for Inside-Out Pharmacology. Biophys. J. 114, 358a (2018).

89. Oswald, C. et al. Crystal structures of the choline/acetylcholine substrate-binding protein ChoX from Sinorhizobium meliloti in the liganded and unliganded-closed states. J. Biol. Chem. 283, 32848–32859 (2008).

90. Wathey, J. C., Nass, M. M. & Lester, H. A. Numerical reconstruction of the quantal event at nicotinic synapses. Biophys. J. 27, 145–164 (1979).

91. Ashford, J. W. Treatment of Alzheimer’s disease: the legacy of the cholinergic hypothesis, neuroplasticity, and future directions. J. Alzheimers Dis. 47, 149–156 (2015).

92. Zemek, F. et al. Outcomes of Alzheimer’s disease therapy with acetylcholinesterase inhibitors and memantine. Expert Opinion on Drug Safety 13, 759–774 (2014).

93. Barbour, B. & Häusser, M. Intersynaptic diffusion of neurotransmitter. Trends in Neurosciences 20, 377–384 (1997).

94. Satin, L. S. & Kinard, T. A. Neurotransmitters and their receptors in the islets of Langerhans of the pancreas: what messages do acetylcholine, glutamate, and GABA transmit? Endocrine 8, 213–223 (1998).

95. Dantzer, R. Neuroimmune Interactions: From the Brain to the Immune System and Vice Versa. Physiol. Rev. 98, 477–504 (2018).

96. Magnon, C. et al. Autonomic nerve development contributes to prostate cancer progression. Science 341, 1236361–1236361 (2013).

97. Mauffrey, P. et al. Progenitors from the central nervous system drive neurogenesis in cancer. Nature 569, 672–678 (2019).

98. Hildebrand, D. G. C. et al. Whole-brain serial-section electron microscopy in larval zebrafish. Nature 545, 345–349 (2017).

99. Kunkel, T. A., Bebenek, K. & McClary, J. Efficient site-directed mutagenesis using uracil-containing DNA. Meth. Enzymol. 204, 125–139 (1991).

100. Studier, F. W. Protein production by auto-induction in high density shaking cultures. Protein Expr. Purif. 41, 207–234 (2005).

101. Battye, T. G. G., Kontogiannis, L., Johnson, O., Powell, H. R. & Leslie, A. G. W. iMOSFLM: a new graphical interface for diffraction-image processing with MOSFLM. Acta Crystallogr. D Biol. Crystallogr. 67, 271–281 (2011).

102. McCoy, A. J., et al. Phaser crystallographic software. J. Appl. Crystallogr. 40, 658– 674 (2007).

103. Winn, M. D. et al. Overview of the CCP4 suite and current developments. Acta Crystallogr. D Biol. Crystallogr. 67, 235–242 (2011).

104. Emsley, P., Lohkamp, B., Scott, W. G. & Cowtan, K. Features and development of Coot. Acta Crystallogr. D Biol. Crystallogr. 66, 486–501 (2010).

105. Adams, P. D. et al. PHENIX: a comprehensive Python-based system for macromolecular structure solution. Acta Crystallogr. D Biol. Crystallogr. 66, 213– 221 (2010).

106. McPhillips, T. M. et al. Blu-Ice and the Distributed Control System: software for data acquisition and instrument control at macromolecular crystallography beamlines. J. Synchrotron Radiat. 9, 401–406 (2002).

107. Kabsch, W. XDS. Acta Crystallogr. D Biol. Crystallogr. 66, 125–132 (2010).

108. Wang, G. et al. An optogenetics- and imaging-assisted simultaneous multiple patch-clamp recording system for decoding complex neural circuits. Nat. Protoc. 10, 397– 412 (2015).

109. Canto, C. B. & Witter, M. P. Cellular properties of principal neurons in the rat entorhinal cortex. II. The medial entorhinal cortex. Hippocampus 22, 1277–1299 (2012).

110. Lim, C.-S. et al. BRaf signaling principles unveiled by large-scale human mutation analysis with a rapid lentivirus-based gene replacement method. Genes Dev. 31, 537–552 (2017).

111. Ding, X. et al. Silencing IFN-γ binding/signaling in astrocytes versus microglia leads to opposite effects on central nervous system autoimmunity. J. Immunol. 194, 4251–4264 (2015).

112. Sauer, M. Localization microscopy coming of age: from concepts to biological impact. J. Cell Sci. 126, 3505–3513 (2013).

113. Thompson, R. E., Larson, D. R. & Webb, W. W. Precise nanometer localization analysis for individual fluorescent probes. Biophys. J. 82, 2775–2783 (2002).

114. Small, A. & Stahlheber, S. Fluorophore localization algorithms for super-resolution microscopy. Nat. Methods 11, 267–279 (2014).

115. Brenner, S. The genetics of Caenorhabditis elegans. Genetics 77, 71–94 (1974).

116. Kim, E., Sun, L., Gabel, C. V. & Fang-Yen, C. Long-term imaging of *Caenorhabditis elegans* using nanoparticle-mediated immobilization. PLoS ONE 8, e53419 (2013).

117. Chai, H. et al. Neural Circuit-Specialized Astrocytes: Transcriptomic, Proteomic, Morphological, and Functional Evidence. Neuron 95, 531–549.e9 (2017).

118. Schindelin, J. et al. Fiji: an open-source platform for biological-image analysis. Nat. Methods 9, 676–682 (2012).

119. Seelig, J. D. et al. Two-photon calcium imaging from head-fixed *Drosophila* during optomotor walking behavior. Nat. Methods 7, 535–540 (2010).

120. Olsen, S. R., Bhandawat, V. & Wilson, R. I. Excitatory interactions between olfactory processing channels in the *Drosophila* antennal lobe. Neuron 54, 89–103 (2007).

121. Thévenaz, P., Ruttimann, U. E. & Unser, M. A pyramid approach to subpixel registration based on intensity. IEEE Trans Image Process 7, 27–41 (1998).

122. Koyama, M., Kinkhabwala, A., Satou, C., Higashijima, S.-I. & Fetcho, J. Mapping a sensory-motor network onto a structural and functional ground plan in the hindbrain. Proc. Natl. Acad. Sci. U.S.A. 108, 1170–1175 (2011).

123. Ono, F., Higashijima, S., Shcherbatko, A., Fetcho, J. R. & Brehm, P. Paralytic zebrafish lacking acetylcholine receptors fail to localize rapsyn clusters to the synapse. J. Neurosci. 21, 5439–5448 (2001).

124. Lu, R., Tanimoto, M., Koyama, M. & Ji, N. 50 Hz volumetric functional imaging with continuously adjustable depth of focus. Biomed. Opt. Express 9, 1964–1976 (2018).

